# The quality of dissolved organic matter shapes the biogeography of the active bathypelagic microbiome

**DOI:** 10.1101/2021.05.14.444136

**Authors:** Marta Sebastián, Pablo Sánchez, Guillem Salazar, Xosé A. Álvarez-Salgado, Isabel Reche, Xosé Anxelu G Morán, M Montserrat Sala, Carlos M. Duarte, Silvia G. Acinas, Josep M. Gasol

## Abstract

The bathypelagic ocean (1000-4000 m depth) is the largest aquatic biome on Earth but it is still largely unexplored. Due to its prevalent low dissolved organic carbon concentrations, most of the prokaryotic metabolic activity is assumed to be associated to particles. The role of free-living prokaryotes has thus been mostly ignored, except that of some chemolithoautotrophic lineages. Here we used a global bathypelagic survey of size-fractionated metagenomic and 16S (genes and transcripts) data and performed a differential abundance analysis to explore the functional traits of the different prokaryotic life-strategies, their contribution to the active microbiome, and the role that the quality of the dissolved organic matter (DOM) plays in driving this contribution. We found that free-living prokaryotes have limited capacity to uplift their metabolism in response to environmental changes and display comparatively lower growth rates than particle associated prokaryotes, but are responsible for the synthesis of vitamins in the bathypelagic. Furthermore, their contribution to the active prokaryotic microbiome increased towards waters depleted of labile DOM, which represented a large fraction of the tropical and subtropical ocean sampled stations. This points to a relevant yet overlooked role of free-living prokaryotes in DOM cycling in the vast bathypelagic desert.

## Introduction

The bathypelagic ocean (1000-4000 m) contains ca. 35% of the ocean’s prokaryotes (Bacteria and Archaea) [1] but remains poorly explored. The energy needs of bathypelagic prokaryotes must be met by intermittent pulses of labile carbon introduced via surface-derived rapid sinking particles [2] or actively transported by zooplankton and vertebrates [3], downward transport of dissolved organic carbon in deep water formation areas [4], suspended small particles from unknown origin [5], or *in situ* chemolithoautotrophy [6]. Due to the prevalent low organic carbon concentrations found in the bathypelagic [4], together with the extreme environmental conditions of low temperature and high pressure typical of this realm, the activity of bathypelagic microbes has traditionally been assumed to be very low. Evidence of relatively high respiration rates [7] and exoenzymatic activities in bathypelagic waters [8], and a relatively modest decrease of prokaryotic growth rate with depth despite the corresponding sharp decrease in prokaryotic cell abundance [1] challenged this view. Yet, there are still a lot of unknowns on the ecology of deep ocean prokaryotes, and how the structure of prokaryotic communities changes across environmental gradients in the bathypelagic realm.

Initial metagenomic studies pointed to a prevalence of the particle-associated life style in bathypelagic waters [9]. This view is currently supported by recent findings using combined-omics approaches hinting that most of the bathypelagic heterotrophic metabolic activity is mediated by particle-associated prokaryotes [10, 11]. However, there is experimental evidence that bathypelagic assemblages are very resilient to the long-term absence of external sources of carbon [12] and contain a seedbank that ensures the long term functionality of the community in the absence of external carbon inputs. These findings, together with observations of potential metabolic versatility in single-amplified-genomes of deep ocean prokaryotes [13–15] and in metagenome-assembled genomes reconstructed from a global bathypelagic metagenomic study [16], question the view that prokaryotic life in the bathypelagic is limited to particles and the initial assumption that prokaryotes inhabiting the free-living fraction live on the DOM released by particle-colonizers during particle degradation [17]. Furthermore, the flux of particles reaching the bathypelagic is variable over time [2, 18, 19], and thus bathypelagic communities must have adapted to these intermittent carbon inputs.

Particle-attached and free-living communities in the bathypelagic differ in dominant phyla and/or classes [20], suggesting that these lifestyles have been strongly conserved through the evolutionary history of deep-sea prokaryotes. One possible driver of this evolution is the nature of the organic carbon sources available to these two communities. Whereas free-living prokaryotes likely face an environment composed mostly of recalcitrant [21] and/or diluted organic compounds [22], particle-associated prokaryotes have access to a more concentrated and fresher organic pool [23, 24]. Size-fractionation approaches have shown that free-living and particle associated prokaryotes in the global bathypelagic have distinct metabolic capabilities [16], but the functional traits of free-living prokaryotes and how they contribute to the active microbiome are still far for being understood, except for some chemolithoautotrophic lineages [6].

Most of the knowledge about the diversity and spatial structuring of deep ocean prokaryotes has been obtained through metagenomics [9, 25] or massive sequencing of the ribosomal 16S RNA gene (hereafter ‘rDNA’) of deep water samples collected in regional and global surveys [26, 27]. Even though these studies have provided very valuable information on the structure of deep ocean communities and possible factors driving this structure, rDNA-based techniques do not discern metabolically active from dead or inactive cells, which is fundamental to understand the role of different members of the prokaryotic communities in the function of the ecosystem. An alternative to delineate the structure of the metabolically active populations is the high-throughput sequencing of the 16S gene cDNA (synthesized from the rRNA, hereafter ‘rRNA’). This approach is not free of criticism, because some taxa are more efficient than others in the use of ribosomes for growth, and some others accumulate ribosomes during dormancy to be able to respond swiftly to sudden changes in resource availability (see [28] for a review on this topic). Detection of rRNA from a given microbe is thus strictly indicative of its “capacity for protein synthesis” [28] and therefore of its potential activity. Despite these caveats, the simultaneous characterization of the 16S-rDNA and rRNA communities has provided interesting insights into the ecology of the potentially active members of microbial communities in lakes [29], marine waters [30], sediments [31], and soils [32], but it has never been attempted in the bathypelagic ocean at a global scale.

Here, we explored the structure of prokaryotic communities with potential for protein synthesis (hereafter referred to as the active prokaryotic microbiome) in the global tropical and subtropical bathypelagic ocean with samples collected during the Malaspina Expedition, and assessed the role that the quality of dissolved organic matter (DOM) plays in driving this structure. To this end, we categorized the different prokaryotic taxa (16S rDNA ASVs) into free-living, particle-attached or dual life-style (i.e. able to thrive in both the free-living and particle associated fraction) lifestyle categories based on differential abundance analyses of these ASVs in the rDNA obtained from the 0.2-0.8 µm and 0.8-20 µm size fractions, respectively. The distinct genetic repertoire of free-living and particle-attached prokaryotes was also delineated using the same differential abundance approach on metagenomic data available from the same samples from both size fractions [16]. We then explored how the contribution of these lifestyle categories to the active bathypelagic microbiome changed with varying dissolved organic matter (DOM) quality, inferred from the optical properties of fluorescent DOM. We hypothesized that the relative contribution of the different lifestyles to the active bathypelagic microbiome will be largely influenced by the composition of the dissolved organic matter.

## Materials and Methods

A total of 101 water samples were collected during the Malaspina 2010 expedition corresponding to 30 different sampling stations globally distributed across the subtropical and tropical region of the world’s oceans between 2,150 and 4,000 m depth (median depth: 4,000 m, Table S1). Samples were obtained for 16S genes (rDNA) and 16S transcripts (rRNA) sequencing of communities from the 0.2-0.8 and the 0.8-20 µm size fraction to discern typical free-living communities from particle associated ones. The 16S rDNA samples were reported before [20, 27] and consist of 60 samples from 30 different stations for which both size fractions are available. The 16S rRNA dataset consists on 41 samples (27 from the 0.2-0.8 µm size fraction and 14 from the 0.8-20 µm size fraction). In this study we used the rDNA samples for the delineation of the different lifestyles and assessing the rRNA: rDNA relationships of the different lifestyles. Details on sample collection, and nucleic acids extraction, sample and sequencing processing can be found in the Supplementary methods. Exact amplicon sequence variants were obtained with DADA2 v1.8 [33].

All raw sequences used in this study are publicly available at the European Nucleotide Archive (ENA, https://www.ebi.ac.uk/ena/browser/home) under Study accession numbers SRP031469 and SRP079340 for the rDNA and rRNA datasets respectively.

### Differential abundance analyses

Studies exploring free-living and particle-attached prokaryotic lifestyles using size-fractionation approaches [34, 35] may be affected by detachment of cells during sample processing [36]. As a result, the free-living fraction may be contaminated by bacteria with a particle associated lifestyle [37]. Thus, with the aim of delineating the different lifestyles in global bathypelagic waters, we performed a differential abundance analysis of individual ASVs in the rDNA across both size fractions using the *corncob* package (v. 0.1.0) [38]. The analysis was performed on rDNA sequences because the amount of ribosomes and transcriptional activity vary strongly among prokaryotes [39], and copiotrophic taxa containing a large number of ribosomes could mask the rRNA sequences of slow growers. This differential analyses allowed us to tackle those ASVs that exhibited significant changes in abundance between the 0.2-0.8 µm size fraction and the 0.8-20 µm size fraction. *Corncob* uses a beta-binomial regression to model ASV abundances, and takes into account zero counts, differences in the sequencing depth between samples, as well as the variability in relative abundances among samples [38]. Wald tests with a false discovery rate (FDR) threshold of 0.05 were applied. Those ASVs predominantly found in the 0.2-0.8 µm size fraction were categorized as having a ‘free-living lifestyle’ (FL-ls). ASVs that did not present significant differences in their distributions in both size-fractions were categorized as ‘dual lifestyle’, and those ASVs predominantly found in the 0.8-20 µm size fraction were considered as ASVs with a ‘particle-associated lifestyle’ (PA-ls). Given the large difference in the sequencing effort for the rDNA and the rRNA (cDNA) pool (Table S1), for downstream analyses the ASV table was sampled down to the minimum number of reads (17,824 reads/sample) to avoid artifacts due to the uneven sequencing effort among samples, using the *rrarefy* function in the *vegan* package. This process was repeated 100 times and the mean number of reads (rounded to integers) from the 100 rarefactions was used.

Functional metagenomic information (KEGG orthology table) for the same sampled stations and size fractions was retrieved from the companion website of [16] https://malaspina-public.gitlab.io/malaspina-deep-ocean-microbiome/page/data/. All the information of how the metagenomes were processed can also be found in that companion website. *Corncob* was also used to delineate the functional traits unique to FL-ls and PA-ls prokaryotes. The KOs shared between the 0.2-0.8 and 0.8-20 µm size fractions were categorized as ‘shared’ since they comprised both core genes shared by the FL-ls and PA-ls lifestyle categories and genes harboured by dual-ls prokaryotes. Raw functional tables were used for the differential analysis and the sub-sampled functional tables of the differentially abundant KOs were used to generate the figures.

All data treatment and statistical analyses were conducted with the R Statistical Software using version 3.2.4.

The rRNA:rDNA relationships of individual ASVs were assessed after a log-centered transformation (CLR) of the rDNA and the rRNA ASV tables, adequate for compositional data [40], using major axis regression with 10000 permutations (R package lmodel2).

### Categorization of the stations based on their FDOM properties

We used a K-means algorithm (*vegan* package) to classify the different sampled stations based on the quality of DOM using as variables for the clustering the amount of recalcitrant (C1+ C2) and labile (C3 +C4) fluorescent DOM, and the labile to recalcitrant ratio (L/R ratio), and forcing the partition into three clusters. This resulted in three clusters of stations whose DOM characterization can be found in Table S2, and comprised waters with high L/R ratios (labile cluster), waters with medium-low L/R ratios (intermediate cluster) and waters with very low L/R ratios (recalcitrant cluster).

### Broad niche preference of bathypelagic prokaryotes

Association of prokaryotic families or individual ASVs in the rRNA pool with the different fluorescent components of the DOM and other biotic and abiotic variables were explored using a Sparse partial least squares (sPLS) regression analysis (see supplementary methods for details).

## Results and discussion

### Different lifestyles contribute to the active bathypelagic prokaryotic microbiome

The differential abundance analysis of individual ASVs across both size fractions (see methods) identified 867 FL-ls ASVs, 532 PA-ls ASVs and 2041 dual-ls ASVs in the entire dataset, as well as 627 very rare ASVs for which we were not able to fit the beta-binomial model (representing only from 0.08 to 5% of the reads, median 0.4%). FL-ls ASVs generally dominated the assemblages of the 0.2-0.8 µm size fraction, with a contribution ranging from 24 to 77% of the reads (mean 55%), although they were sometimes outcompeted by the dual-ls taxa, which represented from 13 to 58% of the reads (mean 33%) (Figure 1a, upper panel). In the 0.8-20 µm size fraction, as expected, FL-ls ASVs only represented 8% of the reads (Figure 1a, upper panel). Conversely, PA-ls ASVs accounted on average for 11% of the reads in the 0.2-0.8 µm size fraction, but represented between 26 and 87% of the community (mean 54%) in the 0.8-20 µm size fraction (Figure 1a, upper panel), while the dual-ls taxa ranged from 10 to 60% in this size fraction. Taxa with a particle-associated lifestyle usually form complex communities on particles, with cell densities orders of magnitude higher than in the surrounding water [41]. However, as particles are degraded, particle-attached communities undergo a succession of taxa that either live on by-product metabolites of the first-colonizers or are specialized on degrading different polymeric material [42]. Thus, taxa associated to particles likely undergo cycles of attachment and detachment [43, 44]. The PA-ls ASV sequences detected in the 0.2-0.8 µm size fraction (∼11%) likely reflect taxa that have recently detached from particles. On the other hand, taxa with a dual-ls likely have the ability to utilize a broad spectrum of substrates and various terminal electron acceptors, allowing them to thrive between encounters with particles and displaying fast growth upon favorable conditions, as it has been suggested previously for successful opportunistic taxa [45, 46]. This probably explains why they can contribute up to 60% of the rDNA sequences in both the 0.2-0.8 µm and the 0.8-20 µm size fractions (Figure 1a, upper panel).

**Figure 1.**
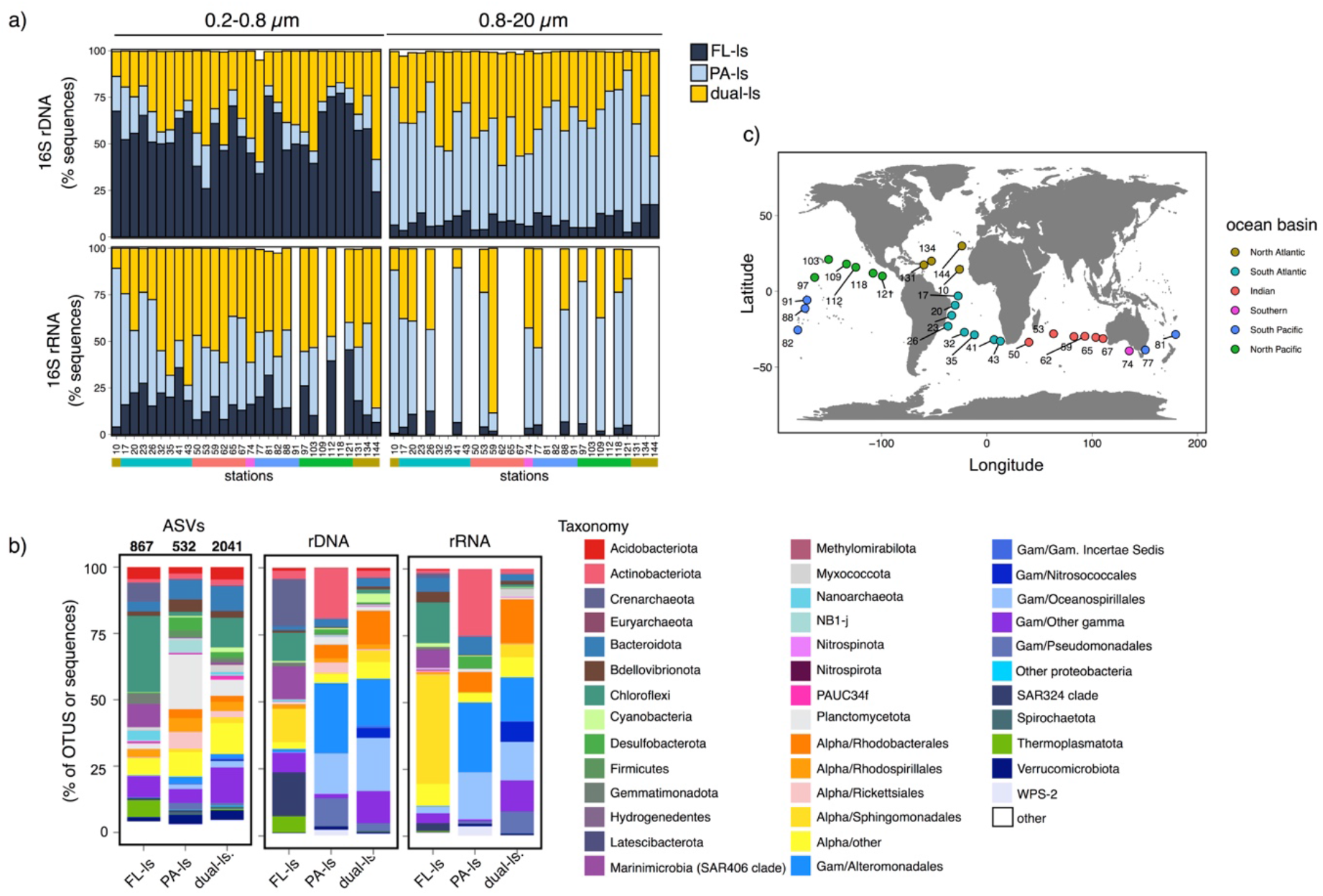
Different life strategies contribute to the total and active bathypelagic prokaryotic microbiome. **a)** Contribution of the life-style categories (based on differential abundance analyses) to bathypelagic prokaryotic communities in the rDNA and rRNA pool from the 0.2-0.8µm and 0.8-20µm size fractions across the global ocean. The white space in top of the bars represent the contribution of those ASVs for which we were not able to fit the model. The color bar below denotes the ocean basins the samples belong to. **b)** Taxonomic composition (proportion of ASVs, and proportion of sequences in the rDNA and rRNA pools) of the three lifestyle categories (Free-living, FL-ls, Particle-associated, PA-ls, and dual lifestyle) shown at the phylum level except for the Proteobacteria, which are divided into classes, and Alphaproteobacteria and Gammaproteobacteria, which are divided into orders. **c)** World map showing the location of the sampled stations. Dots are color-coded based on the ocean basin the samples belong to.

We then explored the contribution of the different lifestyle-resolved categories to the rRNA pool. Regardless of the size fraction, the rRNA-based communities were dominated by dual-ls and PA-ls taxa (Figure 1a, lower panels). FL-ls ASVs contribution to the 0.2-0.8 µm size fraction rRNA pool represented on average 19% (range 4 to 45%), whereas the PA-ls and dual-ls taxa contribution ranged from 8 to 85 and from 11 to 86% (on average 33 and 48%) respectively. PA-ls are thought to be copiotrophic organisms with high ribosome content [47], which probably explains the large contribution of PA-ls taxa to the rRNA of the 0.2-0.8 µm size fraction despite their small contribution in the rDNA. In the 0.8-20 µm size fraction, the rRNA was dominated by the PA-ls assemblages, which accounted on average for 60% of the reads, followed by the dual-ls assemblages, although the number of samples available was half the ones in the 0.2-0.8 µm size fraction. The dominance of PA and dual-ls in the rRNA could suggest that prokaryotes with these lifestyles rule the metabolic potential of the bathypelagic prokaryotic microbiome. Nonetheless, it is also possible that this large contribution in the rRNA reflects higher ribosomal content in prokaryotes with PA and dual lifestyle, which allows them to swiftly respond to the presence of particles or resource patches.

Within each of the lifestyle categories the taxonomic assignation of the ASVs was phylogenetically diverse (Figure 1b), but there were notable differences in how these ASVs contributed to each category in the rDNA and the RNA pool. FL-ls assemblages in the rDNA were largely represented by sequences of the putatively chemolithotrophic Crenarchaeota [48] and SAR324 [14], and by Chloroflexi (mostly SAR202), Marinimicrobia and the Alphaproteobacteria Sphingomonadales, which may have the ability to use recalcitrant compounds [15, 49, 50]. In contrast, in the rRNA pool, Sphingomonadales took over and accounted for half of the reads (Figure 1b). PA-ls rDNA communities were dominated by Actinobacteriota and Gammaproteobacteria belonging to the orders Alteromonadales, Oceanospirillales and Pseudomonadales (Figure 1b), whereas in the rRNA pool the contribution of Pseudomonadales was negligible and instead that of Bacteroidota, Desulfobacterota and Rhodobacterales increased. The dual-ls category was largely represented in both the rDNA and the rRNA by Gammaproteobacteria (mostly Alteromonadales, Nitrosococcales, and Pseudomonadales, Figure 1b), which accounted for >60% of the reads, and Alphaproteobacteria (Rhodobacterales and other less abundant orders). Although this was the average taxonomic composition for the three categories in the entire dataset, taxonomic shifts in the total and active microbiome were also observed across stations (Figure S1 and S2). For example, the rRNA of the PA-ls communities in the 0.8-20 µm size fractions had a large representation of Actinobacteriota in the southernmost stations of the South Atlantic and in the North Pacific, whereas the Gammaproteobacteria Alteromonadales and Oceanospirillales dominated the active PA-ls taxa in the tropical Atlantic. Active FL-ls and PA-ls Bacteroidota ASVs were present in the Indian Ocean in the 0.2-0.8 µm size fraction, and the Gammaproteobacteria Nitrosococcales dominated the active dual-ls ASVs of both size fractions at station 59, in the Indian ocean (Figure S2). Interestingly, PA-ls taxa belonging to the candidate phylum WPS-2 [51] represented up to 20% of the rRNA reads in the 0.2-0.8 µm size fraction in some stations of the Indian Ocean and more than 15% of the 0.8-20 µm size fraction at a station in the Pacific Ocean (Figure S2). This candidate phylum is abundant in organic matter poor soil environments and is thought to encompass high metabolic versatility [52]. The high contribution of WPS-2 to the rRNA pool in stations from different oceans basins points to a relevant but yet unknown role of these bacteria in the deep ocean.

There was an overall positive correlation between the rRNA and rDNA sequences of the individual ASVs belonging to each of the lifestyle categories, but the relationship was weak (Figure 2). rRNA-rDNA relationships have been used to estimate growth rates of individual taxa [53], even though their use has been subject of controversy given the capacity of some bacteria to accumulate ribosomes during dormancy [28]. However, recent findings of positive relationships between RNA content and growth (or activity) of marine bacteria [47, 54] support their use. The slope of the rRNA-rDNA relationship of the ASVs belonging to the FL-ls category was significantly lower than the ones for the dual-ls and PA-ls categories (no overlapping 95% confidence interval of the model II Major Axis regression slopes, 10000 permutations, Figure 2). This would suggest overall comparatively lower growth rates of FL-ls taxa than taxa with a dual or PA lifestyle.

**Figure 2.**
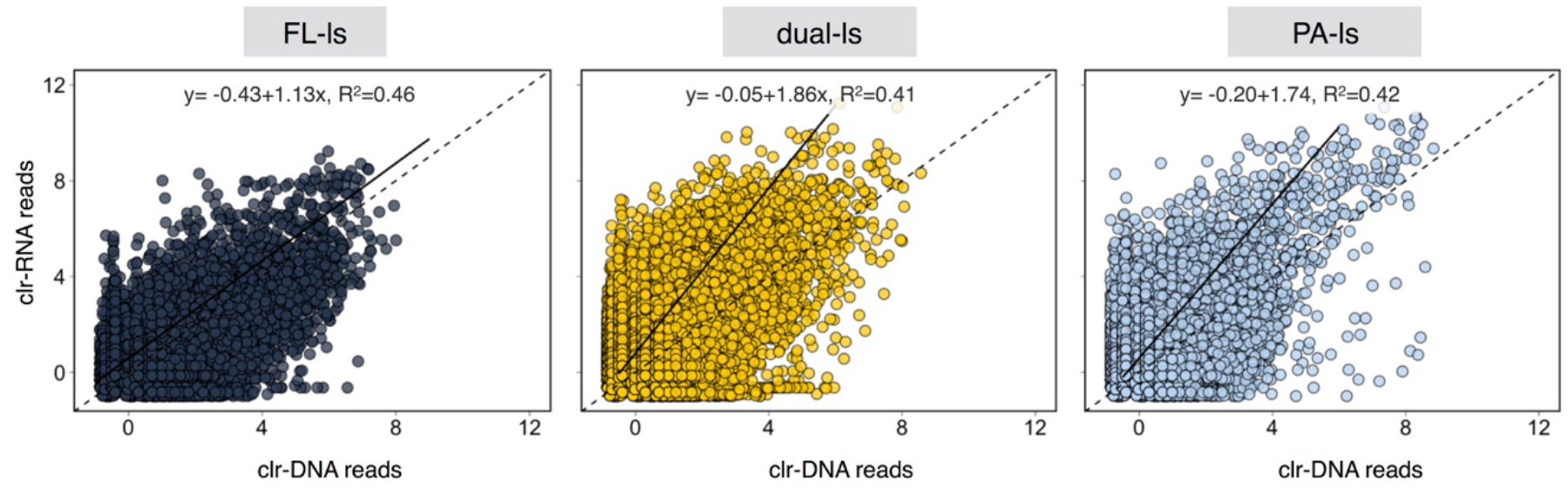
Relationship between RNA and DNA centered-log-ratio normalized reads for the three lifestyle categories. Each data point represents an individual ASV. The dashed line indicates the 1:1 relationship and the solid line the major axis regression. The slope from the FL-ls category is significantly smaller than the slope from the other lifestyle categories (p<0.001, no overlap between the 95% confidence interval for the slope, MA regression, 10000 permutations).

### Differential functional traits of prokaryotes with a free-living and particle associated lifestyle

Using metagenomic data available from the same sampled stations [16], the functional traits of the different lifestyle categories were explored. The same differential abundance analysis that we used to categorize the ASVs was used to categorize the genetic repertoire, as the 0.2-0.8 µm size fraction contained a notable proportion of PA-ls and dual-ls taxa (Figure 1). This approach yielded three categories, one including the genes (based on KO annotations) significantly enriched in the 0.2-0.8 µm size-fraction (FL-ls repertoire,), another one including the genes significantly enriched in the 0.8-20 µm size-fraction (PA-ls gene repertoire), and another one that included both the repertoire of dual-ls taxa, and core functional genes that are shared among the three lifestyles categories. Since we could not differentiate between the core genes and genes harbored by dual-ls taxa, we focused only in the differences in the functional capabilities of FL-ls and PA-ls prokaryotes.

The most striking differences in the gene repertoire of FL-ls and PA-ls taxa were in the contrasting number of genes devoted to environmental and genetic information processing (Figure 3). PA-ls genetic content included a high number of genes coding for two-component systems, substrate binding proteins and permeases that were not detected in the FL-ls gene repertoire (Figure S3). Two-component systems are regulatory systems that enable the cell to sense and respond to changes in environmental cues [55], and they are regarded as the principal mechanism for signal transduction in bacteria [56]. Low numbers of two-component systems have been found in streamlined genomes [55], and in the sunlit ocean the low amount of these genes has been described as a distinctive feature of an oligotrophic lifestyle [57]. The comparatively lower number of two-component systems in FL-ls prokaryotes implies they have a limited capability to regulate gene expression in response to environmental variation, suggesting FL-ls taxa are slow-growers with a relatively uniform metabolism regardless of the external environmental cues, similar to what has been described for dominant oligotrophs in the surface ocean (e.g. SAR11 bacteria [58]). This is in agreement with our finding that FL-ls taxa displayed on average significantly lower rRNA:rDNA values than dual-ls and PA-ls taxa (Figure 2). In contrast, FL-ls prokaryotes contained a large number of genes involved in genetic information processing that were not detected in the PA-ls prokaryotes (Figure 3, Data S1), and proteasome genes, that code for proteases that degrade intracellular proteins and have a role in stress responses [59]. FL-ls communities also displayed a relatively higher number of differentially abundant genes involved in energy metabolism, particularly in ATP synthesis (Figure 3, Figure S4), but genes involved in carbon fixation were mostly shared between both lifestyle categories (Figure S4). Furthermore, genes involved in the synthesis of most vitamins, particularly of vitamins B5, B7 and B12, were found in FL-ls communities, but not in PA-ls communities, indicating that PA-ls prokaryotes are not generally able to synthesize these vitamins and likely obtain them from the particles or the FL-ls communities (Figure 3, and Figure S5). This is supported by the observation that vitamin B7 (Biotin) and B12 transporters were significantly enriched in PA-ls communities (Figure S3). It is known that members of Crenarchaeota (that were numerically important members of the FL-ls communities, Figure 1) encode and express biosynthetic pathways for several B vitamins [60], and release some of these vitamins as a by-product of their metabolism [61]. Yet, we found that the capability to synthesize different B-vitamins was widely taxonomically distributed within FL-ls assemblages (Figure S6). This points to a crucial role of FL-ls prokaryotes in supplying B vitamins for bathypelagic metabolism.

**Figure 3.**
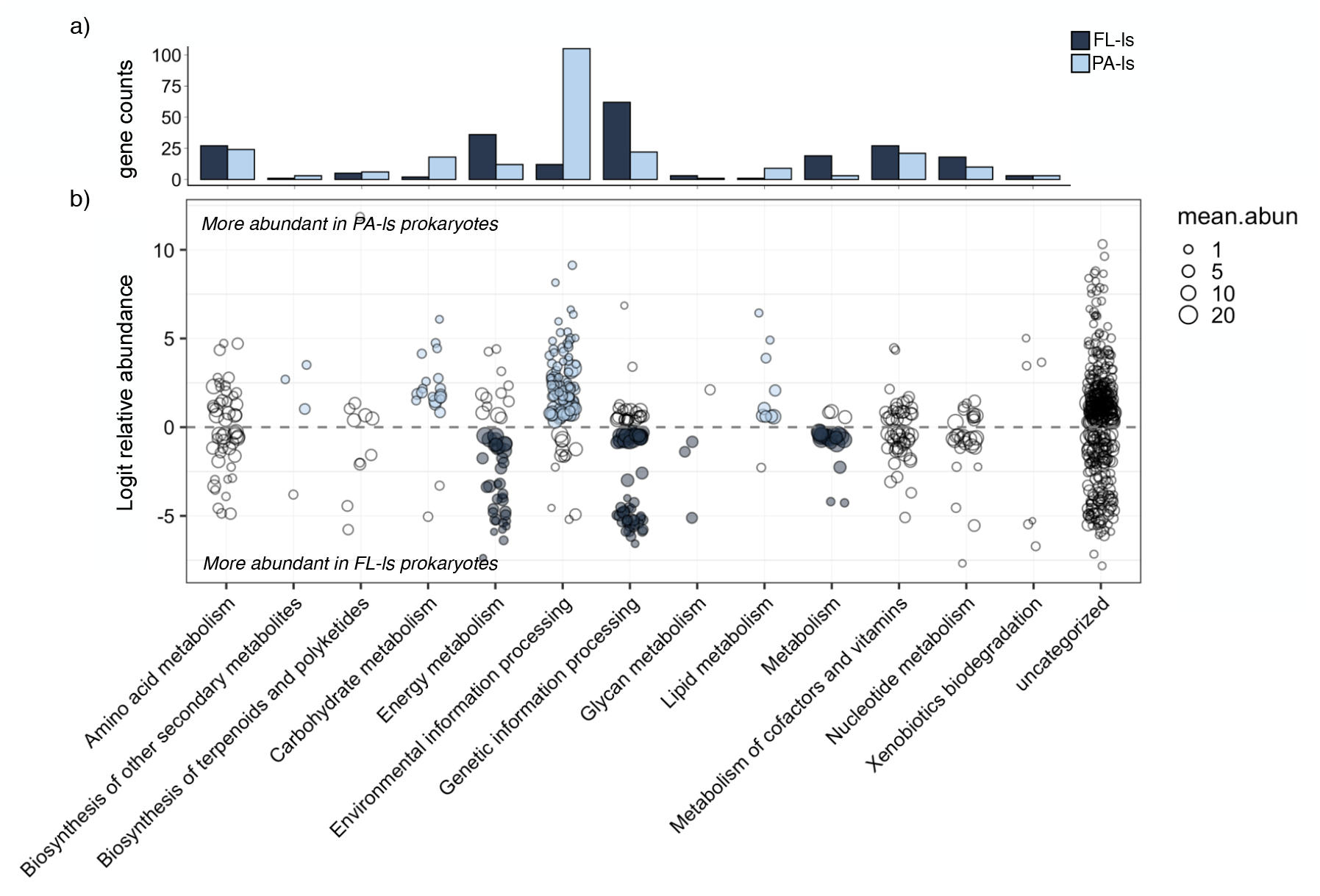
Functional characterization of differentially abundant genes (based on KO annotations) between metagenomes from the 0.2-0.8 µm size fraction and metagenomes from the 0.8-20 µm size fraction. **a)** number of KOs belonging to each of the functional categories that are differentially found in both size fractions. **b)** Details of the differential abundance tests for each size fraction. Each circle represents a differentially abundant KO. Positive values imply that a KO was more abundant in the 0.8-20 µm size fraction (i.e. PA-ls KOs), whereas negative values indicate that the KO was more abundant in the 0.2-0.8 µm size fraction (i.e. FL-ls KOs). Circle size represents the mean relative abundance of each KO in the bathypelagic samples. Colour of circles highlight those functional categories that were clearly enriched in either the FL-ls or the PA-ls prokaryotes (i.e. the number of KOs in a given size fraction doubled the ones detected in the other size fraction).

Besides a broad array of transporters (Figure S3), PA-ls taxa harbored Type II secretion systems (T2SS), which are involved in the export of hydrolytic enzymes like chitinases or proteases [62], as well as in the secretion of adhesins, motility and biofilm formation [63]. The increase in T2SS abundance towards bathypelagic waters has been used to suggest that deep ocean prokaryotes have a preferential particle-associated lifestyle [11]. PA-ls taxa also had a number of genes devoted to carbohydrate metabolism that were not present in the FL-ls communities (Figure 3, Figure S7).

Altogether our analyses indicate that deep ocean communities are formed by an assembly of i) slow growing taxa with a true free-living lifestyle that respond little to environmental cues and are likely adapted to the prevalent DOM scarcity and recalcitrance in the bathypelagic realm, ii) copiotrophic taxa with a pure particle associated lifestyle that probably decrease rapidly in abundance when they are not associated to particles, and iii) taxa with a dual-ls that can live associated to particles, but they can also thrive between successful encounters with particles.

### The quality of dissolved organic matter drives the structure of the bathypelagic prokaryotic microbiome

To further confirm that the quality of DOM plays a role in the structuring of the active bathypelagic prokaryotic microbiome, we classified the sampled stations according to their DOM characteristics. To assess the quality of DOM we looked at the optical properties of the fluorescent fraction of DOM (FDOM), which provide insight into the nature of the refractory DOM pool (see [64, 65] for a complete characterization of the FDOM during the Malaspina expedition). FDOM was characterized by two humic-like recalcitrant components, C1 and C2, and two comparatively more labile components, C3 and C4, attributed to the amino acids tryptophan and tyrosine respectively [64]. Using a standard K-means classification algorithm we grouped the stations into three different clusters based on the maximum fluorescence of the recalcitrant (C1+C2) and labile (C3+C4) components, as well as the labile to recalcitrant FDOM (L/R) ratio (Figure 4a). Most of the waters in the stations sampled were characterized by an intermediate or low L/R ratio (Figure 4a,b), and only two stations in the central Atlantic were characterized by high L/R ratios. The three clusters of stations also presented contrasting values of apparent oxygen utilization (AOU) (Figure 4b), with significant differences between the low and high L/R ratio clusters (pairwise Wilcoxon test, p<0.05). AOU represents the amount of oxygen consumed since the last contact of a given water parcel with the atmosphere, and thus integrates respiratory processes and provides an idea of water ageing [66]. A decrease in DOM lability was also accompanied by a decrease in average specific prokaryotic growth rates, although in this case the differences between clusters were not significant (Kruskal-Wallis test, p> 0.05, Figure 4b). Regarding the contribution of the different life styles to the active microbiome (rRNA), no clear trend was observed in the 0.8-20 µm size fraction, with a dominance of PA-ls taxa in the three clusters, as expected (Figure 4c). However, we found a decreasing trend in the contribution of PA-ls taxa to the rRNA of the 0.2-0.8 µm size fraction with decreasing levels of DOM lability (Figure 4c), which went from ∼70% to <20%. The labile amino acid-like DOM is linked to surface water production [67] and thus mostly originates from solubilization of the proteinaceous material of large sinking particles rapidly reaching the deep ocean. Indeed, sinking particles are known to produce a plume of labile DOM during their transport to deep waters [17], which may fuel the activity of the copiotrophic PA-ls taxa detached from particles, explaining the high contribution of PA-ls taxa in the 0.2-0.8 µm rRNA of the stations enriched in labile material.

**Figure 4.**
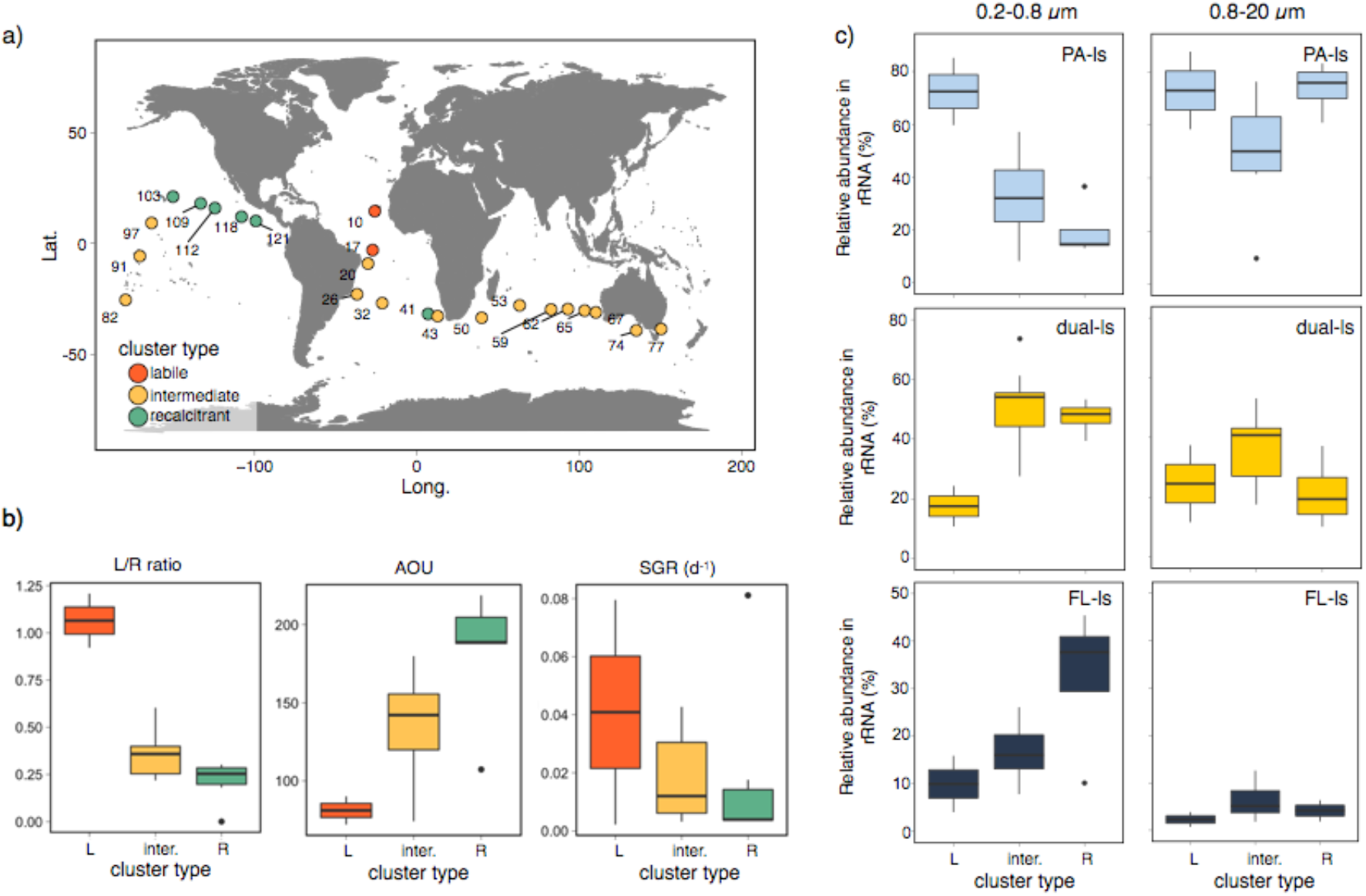
Varying contribution of free-living, dual and particle-attached lifestyles to the active prokaryotic microbiome in waters with different dissolved organic matter quality. K-means clustering of stations based on the amount of the different FDOM components and the labile to recalcitrant (L/R) FDOM ratio. **a**) Map showing the distribution of the stations colored by the cluster they belong to: labile (high values of L/R ratio, see Table S2 for details), intermediate and recalcitrant (low values of L/R ratio). **b)** Boxplots showing the variation of the L/R FDOM ratio, the apparent oxygen utilization (AOU), which integrates respiratory processes, and the average specific prokaryotic growth rates in the labile (L), intermediate (inter.) and recalcitrant (R) cluster of stations. c) Percent contribution of the different lifestyle categories to the rRNA in each of the cluster types, in both size fractions.

The enhanced specific growth rates observed in these stations where PA-ls taxa dominate agrees with the observation that particle associated prokaryotes display higher growth rates than the prokaryotes in the free-living realm [68] and the higher rRNA:rDNA ratios of PA-ls taxa relative to the FL-ls (Figure 2). In contrast, despite their presumably slow growth rates, FL-ls ASVs represented over 30% of the rRNA-based communities in the waters depleted in labile DOM, and together with dual-ls taxa they made up to 75% of the rRNA in these waters (Figure 4c).

Even though particle-associated prokaryotes are assumed to dominate deep sea metabolism [1, 10, 11, 69], their contribution to the active microbiome will rely largely on their abundance, their per-cell activity, and the occurrence of their particulate habitats [70], which is variable over temporal and spatial scales. Bottle samplers are known to underestimate particle abundances [71], but particle-associated prokaryotes are thought to generally represent less than 10% of the total prokaryotic abundance in the bathypelagic [72, 73]. This percentage could decrease further in waters with reduced particle supply, such as the tropical and subtropical ocean, given the presumably lower abundance of particles. In these oligotrophic waters, the older particles impoverished in nutrients may be comparatively less populated by prokaryotes, and free-living assemblages (i.e the 0.2-0.8 µm size fraction) may largely dominate bathypelagic prokaryotic abundance. We found that intermediate and recalcitrant DOM clusters made up 90% of the sampled stations across the tropical and subtropical bathypelagic ocean (Figure 4a). Hence, the enhanced contribution of FL-ls and dual-ls taxa to the 0.2-0.8 µm active microbiome in these waters with decreasing proportion of labile DOM, implies that the view that life in the bathypelagic is mostly limited to particle-attached prokaryotes is oversimplified. In fact, we found that the dominant contributors to the rRNA in the FL-ls category were Sphingomonadales, which seem to cope well with the long-term absence of fresh organic carbon, as in an experiment with bathypelagic communities with no external organic carbon input they represented up to a third of the rRNA based prokaryotic community during the first 5 months ([12], Figure S8). The exploration of the metabolic profiles of prokaryotic communities by means of Biolog GN2 plates^®^ showed that communities from waters depleted in labile DOM displayed higher capability in the utilization of polymers relative to carbohydrates than communities in waters with a higher proportion of labile DOM (Figure 5). This indicates that communities adapted to conditions of low labile DOM enhance their ability to use recalcitrant compounds, which agrees with the observed increase in the ratio of polymers to carbohydrates utilization from the surface to the deep ocean [74] due to the increase in the recalcitrant nature of DOM [64].

**Figure 5.**
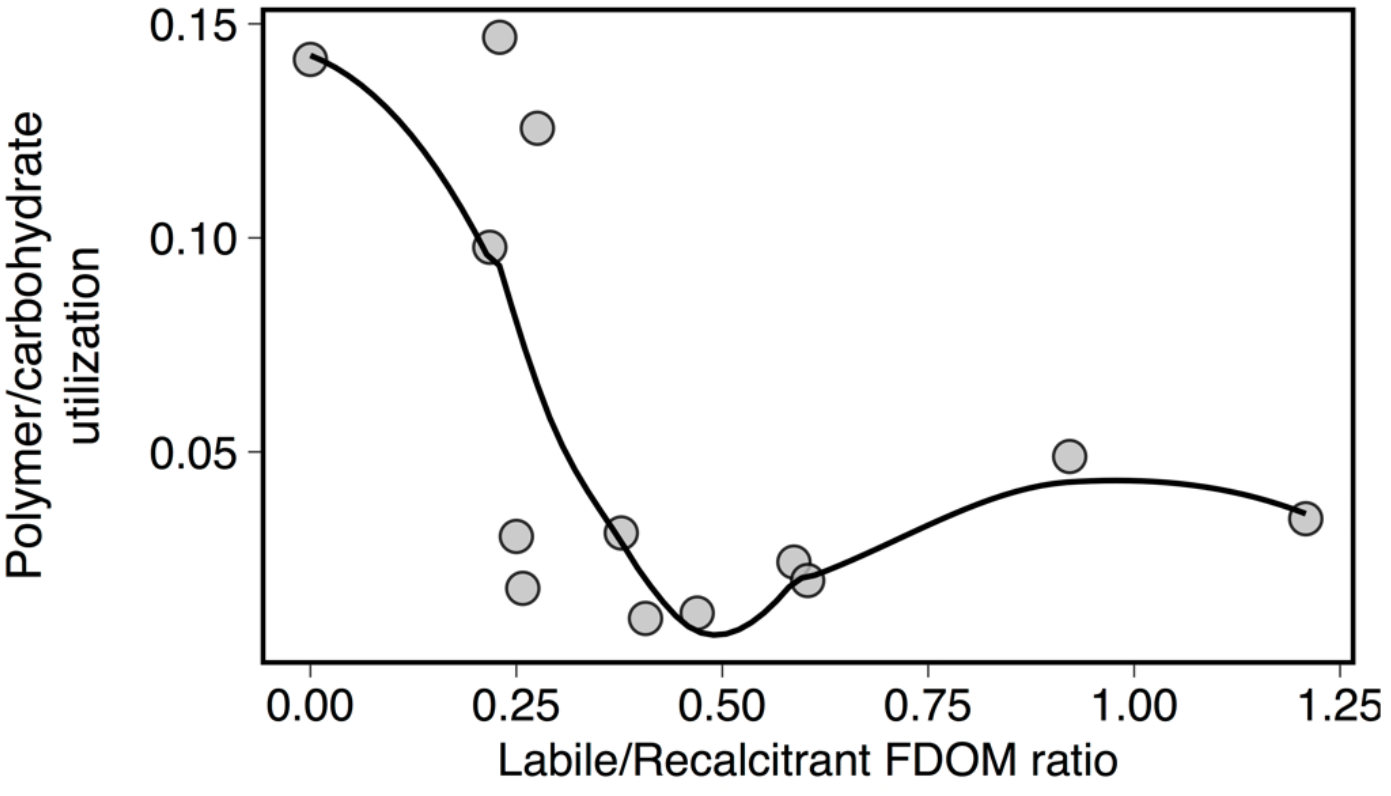
Relationship between the labile to recalcitrant FDOM ration and the ratio of utilization of polymers versus carbohydrates by bathypelagic prokaryotic communities. Prokaryotic substrate use was estimated using Biolog GN2 plates. The line is a best fit smooth curve through the centre of the data calculated using weighted least squares to highlight the relationship between the two variables.

DOM quality also had a significant effect on the contribution of distinct prokaryotic families or individual ASVs to the rRNA pool of both size fractions (Figure 6 and Figure S9, S10), regardless their lifestyle strategy (Figure S9, S10). Spare Partial Least Square (sPLS) regression analyses showed a subset of prokaryotic families (or individual ASVs) that appeared associated with younger waters (i.e. low AOU, [64]) enriched in labile compounds (high labile to recalcitrant ratio), and other prokaryotic families associated with aged waters (high AOU) and with an enrichment in the recalcitrant components C1 and C2, in both size fractions (Figure 6 and Figure S9, S10). This suggests that despite the activity of prokaryotes in the 0.8-20 µm size fraction is likely more insensitive to changes in the surrounding water than in the 0.2-0.8 µm size fraction [75], particles from older waters are distinct to those from younger waters, probably because they have gone through longer remineralization and transport histories, leading to the presence of different active ribotypes. This hypothesis is supported by recent findings showing that in productive regions, where higher particle flux is expected, bathypelagic particle-associated communities are similar to particle-associated communities in the surface, and show different community structure than particle-associated communities from less productive regions [76]. The broad niche partitioning observed among bathypelagic prokaryotes at the global scale (see supplementary information for further discussion on the niche preference of particular taxonomic groups), point to a strong linkage between the relative activity of different bathypelagic prokaryotes and the quality of the DOM pool. The study of the relationship of DOM and members of the ocean prokaryotic microbiome is slowly gaining momentum [77–80] and it seems a necessary step to improve our understanding of microbial processes in the ocean and the key players involved.

**Figure 6.**
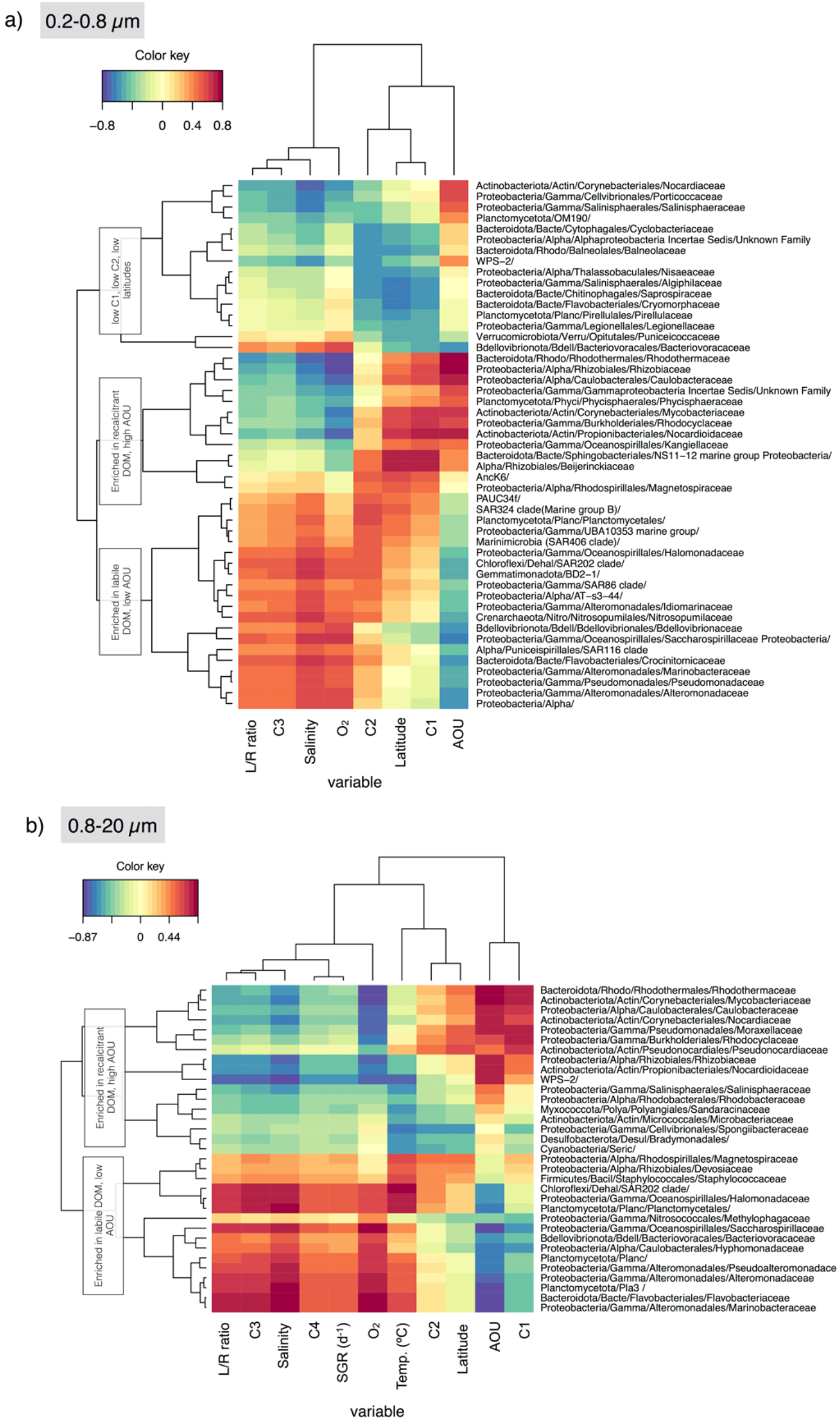
Broad niche partitioning of bathypelagic prokaryotic families. Pair-wise associations between the relative abundance in the rRNA pool of different prokaryotic families and the absolute fluorescence of the FDOM components, the labile to recalcitrant FDOM ratio and other abiotic and biotic variables. Only those families that reached 100 sequences in the rRNA pool were considered for the analysis. **a)** 0.2-0.8 µm size fraction, **b)** 0.8-20 µm size fraction. The color key represents the strength of the positive or negative association.

Our work provides compelling evidence that microbial life in the bathypelagic is not limited to particles. We demonstrate that there is a fraction of prokaryotic communities thriving as free-living organisms and well adapted to scarce DOM availability. These prokaryotes comprise both slow growers that have a free-living lifestyle and limited metabolic flexibility to respond to environmental changes, and dual lifestyle taxa, which are capable of fast growth and also able to cope with low carbon inputs, perhaps by using various terminal electron acceptors, as reported for other marine opportunistic taxa [46]. The contribution of these free-living and dual-ls taxa to the active microbiome increases when the availability of labile DOM decreases, the conditions prevailing in the global bathypelagic desert. Given the scarcity of particles in the bathypelagic realm, these findings suggest an unrecognized role of obligate and optional free-living prokaryotes in this ecosystem, which may have strong implications for its functioning.

## Supporting information

Supplementary DataS1

## Acknowledgments

This research was funded by the Spanish Ministry of Economy and Competitiveness Science and Innovation through the Consolider-Ingenio programme (project Malaspina 2010 Expedition, ref. CSD2008–00077). We thank F.M. Cornejo-Castillo, E. Borrull, C. Díez-Vives, E. Lara, and D. Vaqué for their help in sampling. We also thank our fellow scientists and the crew and chief scientists of the different cruise legs for the smooth operation. Sequencing at the JGI was supported by U.S. Department of Energy (DOE) JGI 2011 Microbes Program grant CSP 387 grant to S.G.A. The work conducted by the U.S. Department of Energy Joint Genome Institute is supported by the Office of Science of the U.S. Department of Energy under Contract No. DE-AC02-05CH11231. Additional funding was provided by MAGGY (CTM2017-87736-R) funded to SGA and through the Spanish Ministry of Economy and Competitivity grants MALASPINOMICS (CTM2011-15461-E), DOREMI (CTM2012-34294), ANIMA (CTM2015-65720-R) and MIAU (RTI2018-101025-B-I00). GS held a Ph.D. JAE-Predoc (CSIC). This work has been carried out with the institutional support of the ‘Severo Ochoa Centre of Excellence’ accreditation (CEX2019-000928-S). We thank Clara Ruiz-González and Francisco M Cornejo Castillo for their comments on a previous version of the manuscript.

## Author Contributions

GS contributed to the collection of microbial samples. MS performed RNA extractions. GS and MS performed the sequence data processing and the statistical analyses. CMD led the Malaspina 2010 Expedition and JMG led the microbial block of the expedition and both provided the ancillary data. SGA obtained the JGI grant that made possible the sequencing of the DNA and RNA samples. PS helped with the bioinformatics work. MMS provided the BIOLOG data and IR and XAAS provided the FDOM data. JMG and XAGM provided prokaryotic specific growth rates. MS conceived and wrote the paper. All authors contributed to the final version of the paper.

