## Supplemental information for "The quality of dissolved organic matter shapes the biogeography of the active bathypelagic microbiome"

**This PDF file includes:**
Supplementary methods
Supplementary discussion
Figure S1 to S10
Tables S1 to S2
Supplementary references

**Other Supplementary Materials for this manuscript include the following:**

Dataset 1

### Supplementary Methods

*Sample collection*

Water was collected with Niskin bottles mounted on a rosette sampler equipped with conductivity–temperature–depth (CTD) (Seabird SBE 911) and dissolved oxygen (SBE 43) profilers. Apparent oxygen utilization (AOU) was calculated as the difference between the saturation and measured dissolved oxygen concentrations.

For nucleic acids collection, samples were pre-filtered through a 200 µm and a 20 µm mesh to remove large plankton. For the DNA analysis, samples were collected and extracted as described elsewhere (1, 2). For the rRNA analyses, 12 L were sequentially filtered (within ~ 15 minutes) through a 142 mm polycarbonate membrane filter of 0.8 µm (Merk Millipore, Isopore polycarbonate) and a 0.2 µm pore size (Merk Millipore, Isopore polycarbonate). The filtration was done with a peristaltic pump (Masterflex, EW-77410-10). The filters were then flash-frozen in liquid $N_2$ and stored at -80ºC until extraction. RNA was extracted with the RNEasy kit (Qiagen) procedure following manufacturer's instructions. Residual DNA was removed using the Turbo DNA-free kit (Applied Biosystems, Austin, TX, USA) and the absence of rDNA in the RNA sample was verified through PCR. RNA was reverse transcribed using random hexamers and the SuperScriptIII kit (Invitrogen) according to the manufacturer's instructions.

*16S Sample sequencing and processing*

Prokaryotic diversity in the rDNA and the rRNA pool was assessed through amplicon sequencing of the V4 region of the 16S rDNA gene in the Illumina MiSeq platform using paired-end reads (2 X 250 bp). All library construction and sequencing was carried out at the JGI (www.jgi.doe.gov) following an standard protocol (3). The variable region V4 of the 16S rDNA gene was amplified using primers F515/R806 (5'-GTG CCA GCM GCC GCG GTA A-3' / 5'-GGA CTA CHV GGG TWT CTA AT-3'). Before sequencing, PhiX spike-in shotgun library reads were added to the amplicons pool for a final concentration of about 20-25% of the pair-end reads library as an internal standard. Although primer 806R has been shown to underestimate the abundance of SAR11 (4), a good agreement was previously found for the rDNA dataset with data derived from metagenomes, and thus not dependent on primers (2). Exact amplicon sequence variants were obtained with DADA2 v1.8 (5).

Primers and spurious sequences were trimmed using cutadapt (6). For the rDNA sequences it was necessary to run it twice using the following parameters: --pair-filter=any –interleaved --minimum-length=32. DADA2 v1.8 was used to differentiate exact sequence variants (5). DADA2 resolves ASVs (amplicon sequence variants) by modelling the errors in Illumina-sequenced amplicon reads. The approach is threshold free, inferring exact variants up to 1 nucleotide of difference using the quality scores distribution in a probability model. After filtering through DADA2, 75% of the total rDNA reads and 90% of the rRNA reads were retained for further analyses. Taxonomic assignation was performed using the function 'assignTaxonomy' against SILVA v.138.

*Free-living prokaryotic abundances and bulk heterotrophic activity*

Prokaryotic abundance was determined by flow cytometry as described in (7). Prokaryotic biovolume was estimated using the SSC flow cytometry signal as in (8) assuming a spherical shape and converted to biomass using the empirical equation described in (9) and prokaryotic abundance. Bulk prokaryotic heterotrophic production was estimated using $^3$H-leucine incorporation as follows: two 40-ml aliquots and two formaldehyde-killed controls (2% final conc.) were incubated at *in situ* temperature with $^3$H-leucine (160 Ci mmol$^{-1}$, 5 nM final conc.) for 6 up to 14 hours depending on the expected activity. Incubations were stopped with formaldehyde (2% final conc.), samples were filtered through 0.2 µm filters, and rinsed three times with 5 ml of cold TCA (5%). The filters were then introduced in scintillation vials, 1 mL of liquid scintillation cocktail (Optimal HiSafe) was added to each of the vials, and the radioactivity was counted on a Beckman scintillation counter after a minimum of 24 h storage. Conversion of leucine to carbon units (prokaryotic heterotrophic production) was done with the theoretical factor 1.5 kg C mol Leu$^{-1}$ (10). Specific growth rates were calculated using the following equation: ln(1+ Prokaryotic het. production/Prokaryotic biomass).

*Analysis of the fluorescent dissolved organic matter (FDOM)*

The optical properties of fluorescent DOM were used to evaluate the quality of dissolved organic matter. These optical properties were described on the basis of fluorescence excitation/emission matrices (EEMs) as explained in (11, 12). Four main fluorescence components were recovered from the EEMs using parallel factor analysis (PARAFAC): Components C1 and

C2, previously related to refractory, humic-like material, and C3 and C4, that represent more biodegradable and fresher microbially-produced FDOM (for more details see (11, 12). The labile (C3 + C4) to recalcitrant (C1 + C2) fluorescence (L/R) ratio was used as an index of the relative lability of the DOM pool.

*Substrate utilization capabilities*

The prokaryotic capability to use different DOM substrates was evaluated by means of Biolog GN2 plates, as fully described in (13).

*Broad niche preference of bathypelagic prokaryotes*

Association of prokaryotic families or individual ASVs in the rRNA pool with the different fluorescent components of the DOM and other biotic and abiotic variables were explored using a Sparse partial least squares (sPLS) regression analysis. To that end, we used the 'mixOmics' package (14) in R in 'regression' mode with families that at least displayed 100 rRNA reads in the dataset or individual ASVs representing more than 1% of the rRNA-based community in at least one sample. Only those families/ASVs detected also in the rDNA pool were considered for the analysis. We run the analysis for each of the size fractions independently using the clr-transformed reads and included the following variables: Latitude, Temperature, Salinity, oxygen concentration, apparent oxygen utilization (AOU), heterotrophic prokaryotic production (estimated as leucine incorporation), prokaryotic biomass, prokaryotic cell abundance, specific prokaryotic growth rates, the FDOM components C1, C2, C3 and C4 , and the labile to recalcitrant ratio. Additionally we used the mean annual chlorophyll-*a* concentration and primary production associated to the Longhurst province to which each station belonged to as proxies for the annual average productivity conditions within the sunlit layer of each station. These data were downloaded from the Marine Regions site (https://www.marineregions.org/downloads.php), and represent mean annual chlorophyll-*a* concentrations (mg $m^{-2}$) and mean daily rates of net primary production (gC $m^{-2}$ $d^{-1}$) integrated for the photic zone. Leave-one-out cross-validation was performed to evaluate model performance (15). sPLS allowed the simultaneous selection of meaningful variables in both the rRNA and the environmental dataset, and the resulting pair-wise associations were visualized using clustered image maps (16).

### Supplementary discussion

*Broad niche-partitioning of bathypelagic prokaryotes*

The rRNA contribution of members of the SAR324 clade, the candidate phylum PAUC34f, the phyla Gemmatimonadota and Myxococcota, the Nitrosopumilaceae family, and the SAR202 clade to the 0.2-0.8 µm size fraction showed significant associations with waters enriched in labile compounds (Figure and Figure S9, S10). Both SAR324 and Nitrosopumilaceae have the potential for chemolithotrophy [14, 48], but they rely on the oxidation of sulfur compounds and ammonia, respectively, as sources of energy to fix inorganic carbon, and these reduced compounds are also released through particle solubilization. This could explain why they are associated with the labile DOM components. SAR202 bacteria encode proteins to use recalcitrant organic compounds [15], and it was recently experimentally demonstrated that SAR202 populations respond to the addition of lignin [77]. However, SAR202 bacteria did not appear associated with the ageing waters enriched in recalcitrant DOM (Figure 6 and Figure S9), suggesting that other factors like microbial interactions or extra carbon or energy needs play a role in controlling the activity of this prokaryotic group. Indeed, SAR202 cells have also been shown to actively participate in amino acid consumption in deep waters using single-cell approaches [78], suggesting they can benefit from both labile and recalcitrant compounds.

In the 0.8-20 µm size fraction, two Planctomycetota families (Pla3 lineage and Planctomycetes), Flavobacteriaceae and some Gammaproteobacteria families, mostly belonging to the Alteromonadales order, were associated with labile DOM and with high specific prokaryotic growth rates (Figure 6). This is consistent with previous findings showing that Alteromonadales display fast growth rates in the ocean [53], play a pivotal role in the processing of labile DOM [79], and respond swiftly to inputs of organic carbon throughout the water column [80].

Conversely, aged waters (i.e. characterized by high AOU) with high signatures of the humic-like components and low specific prokaryotic growth rates were significantly associated with the rRNA sequences of other families of Gammaproteobacteria (Rhodocyclaceae, Moraxellaceae and Kaniellaceae) and Alphaproteobacteria (Caulobacteraceae, Rhizobiaceae, Beijerinckiaceae) in both size fractions (Figure 6), as well as several families of Actinobacteriota mainly in the 0.8-20 μm size fraction.

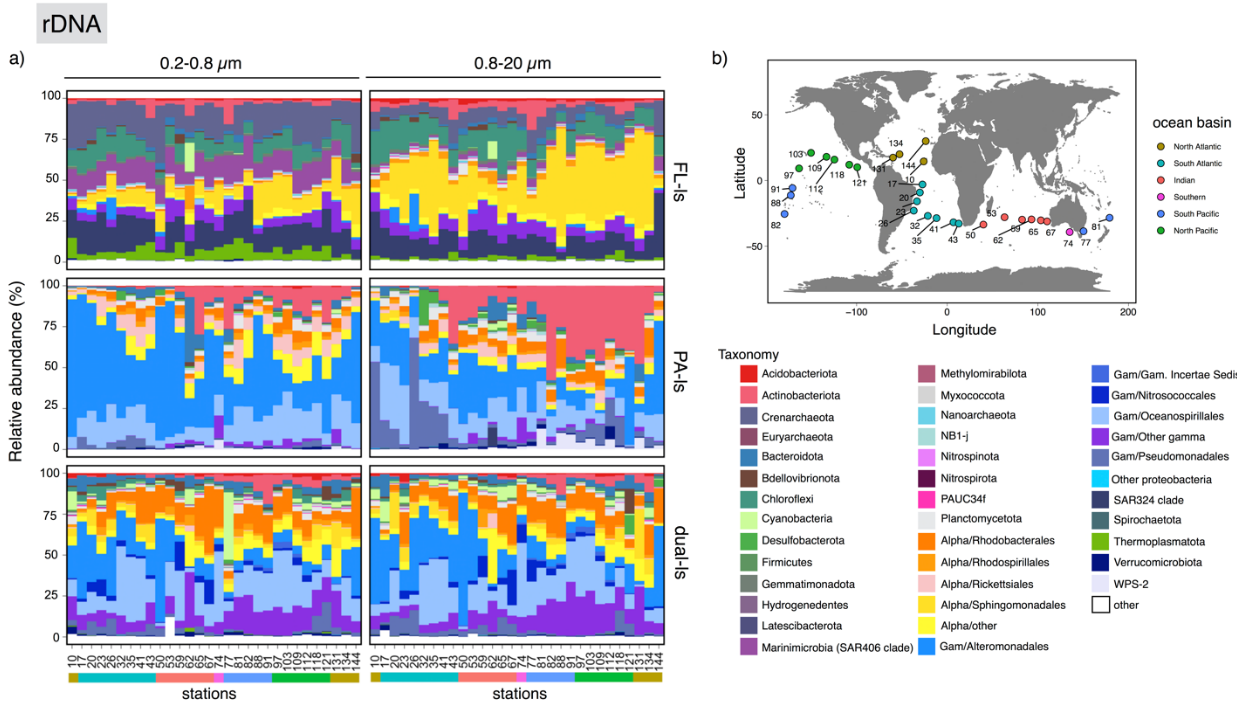


**Figure S1. a)** Contribution of the different taxonomic groups to the rDNA pool from the 0.2-0.8μm (left panels) and 0.8-20μm (right panels) size fractions across the global bathypelagic ocean. Each panel represent the DNA reads of the ASVs within each of the different lifestyle categories (FL-ls, PA-ls and dual-ls). The color bar below denotes the ocean basins the samples belong to. **b)** map of the stations (colored based on ocean basin) to facilitate interpretation.

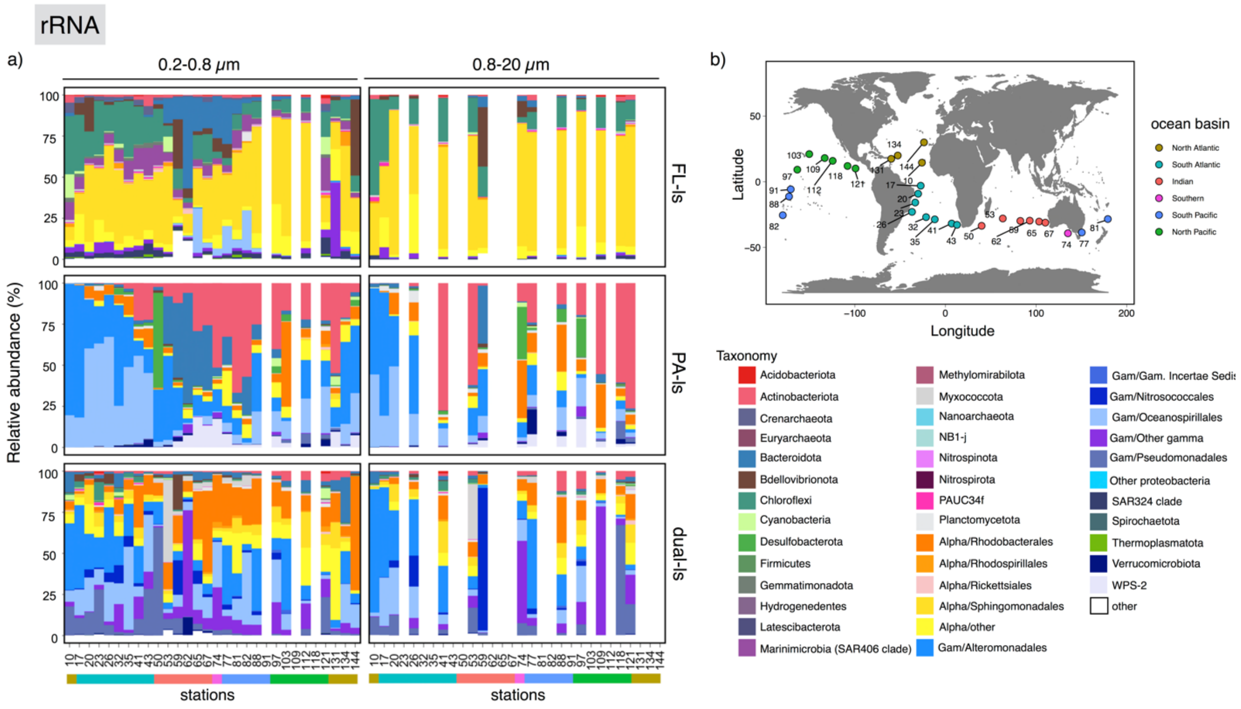

**Figure S2. a)** Contribution of the different taxonomic groups to the rRNA pool from the 0.2-0.8µm (left panels) and 0.8-20µm (right panels) size fractions across the global bathypelagic ocean. Each panel represent the RNA reads of the ASVs within each of the different lifestyle categories (FL-ls, PA-ls and dual-ls). The color bar below denotes the ocean basins the samples belong to. **b)** map of the stations (colored based on ocean basin) to facilitate interpretation.

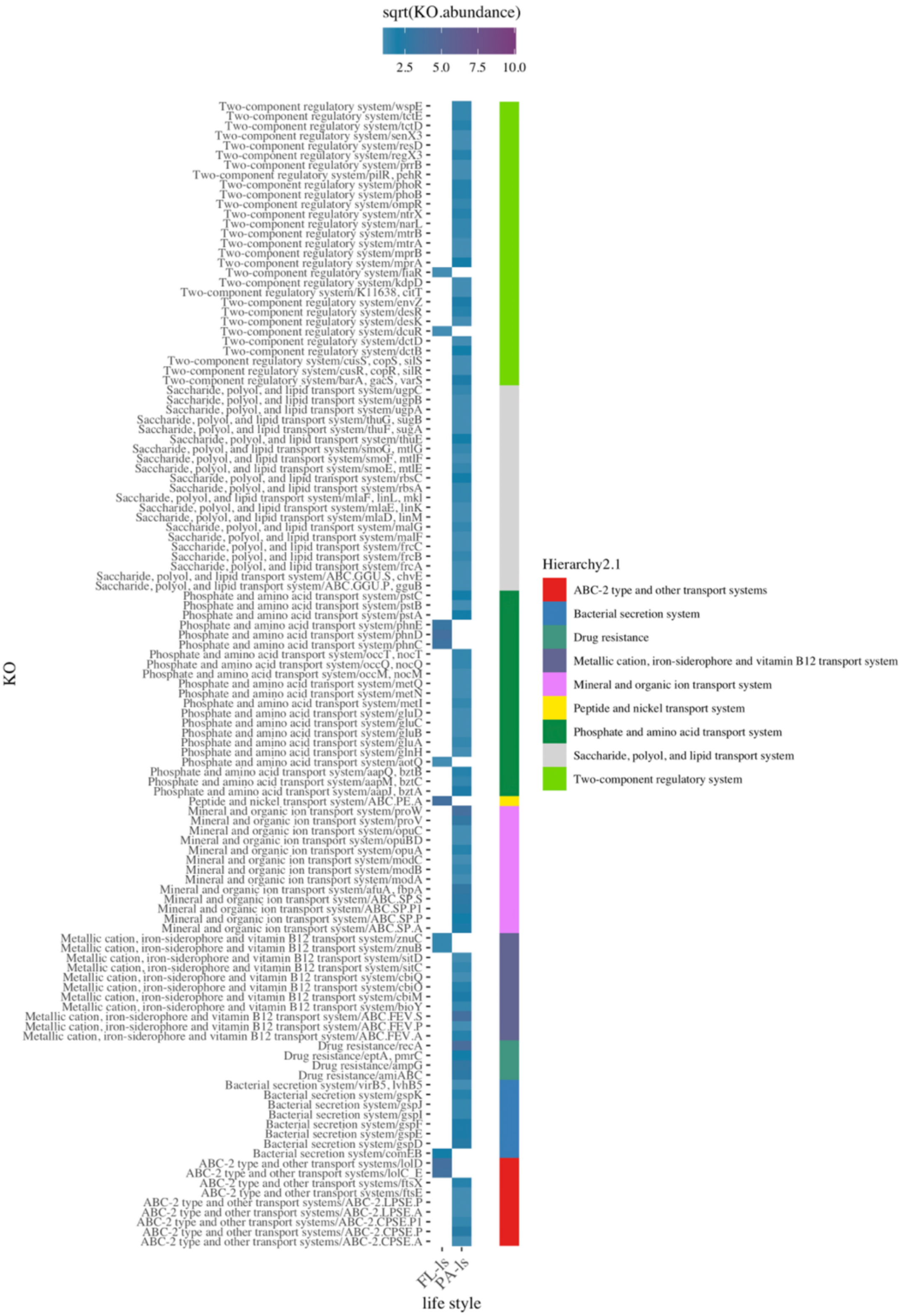


**Figure S3.** Differentially abundant genes (based on KO annotations) in environmental processing in the FL-ls and PA-ls categories.

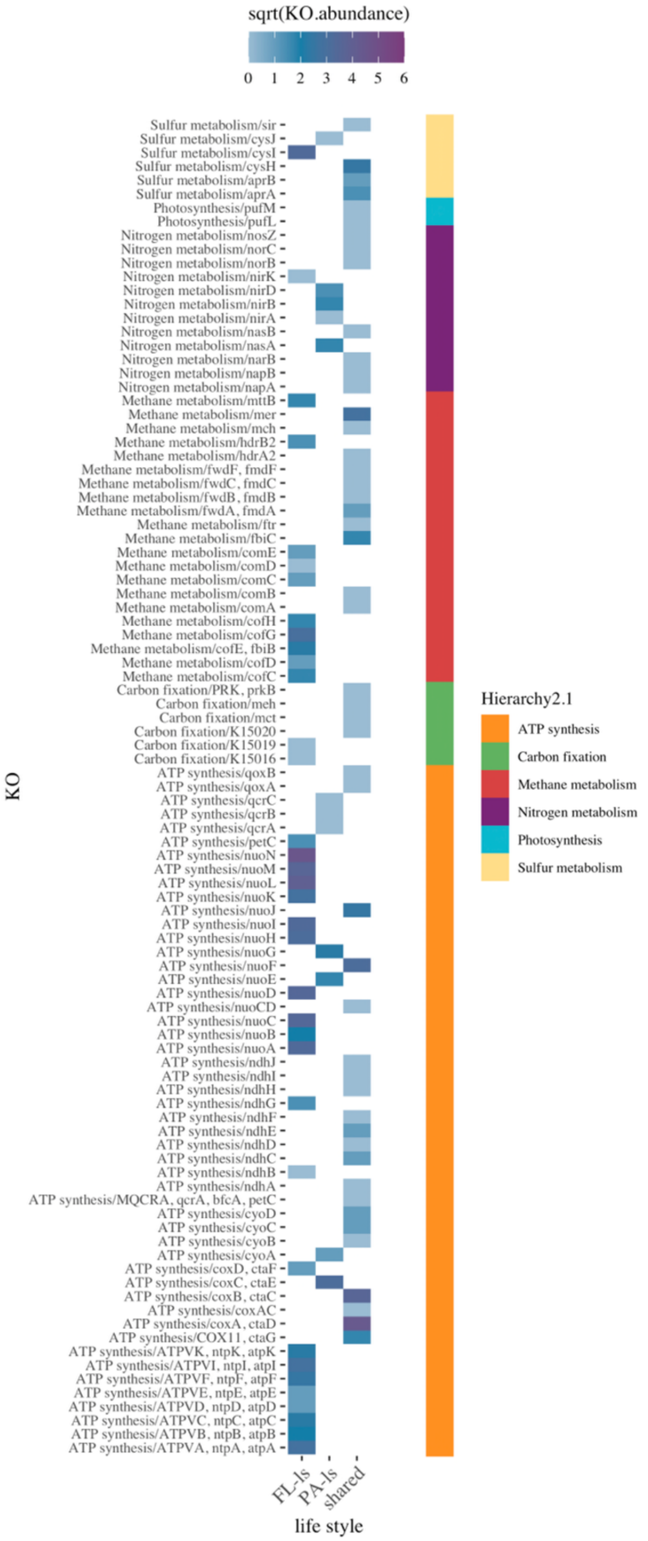


**Figure S4.** Differentially abundant genes (based on KO annotations) in energy metabolism in the FL-ls and PA-ls categories. The shared category includes both the genetic repertoire of dual-ls taxa and KOs shared by the FL-ls and PA-ls categories.

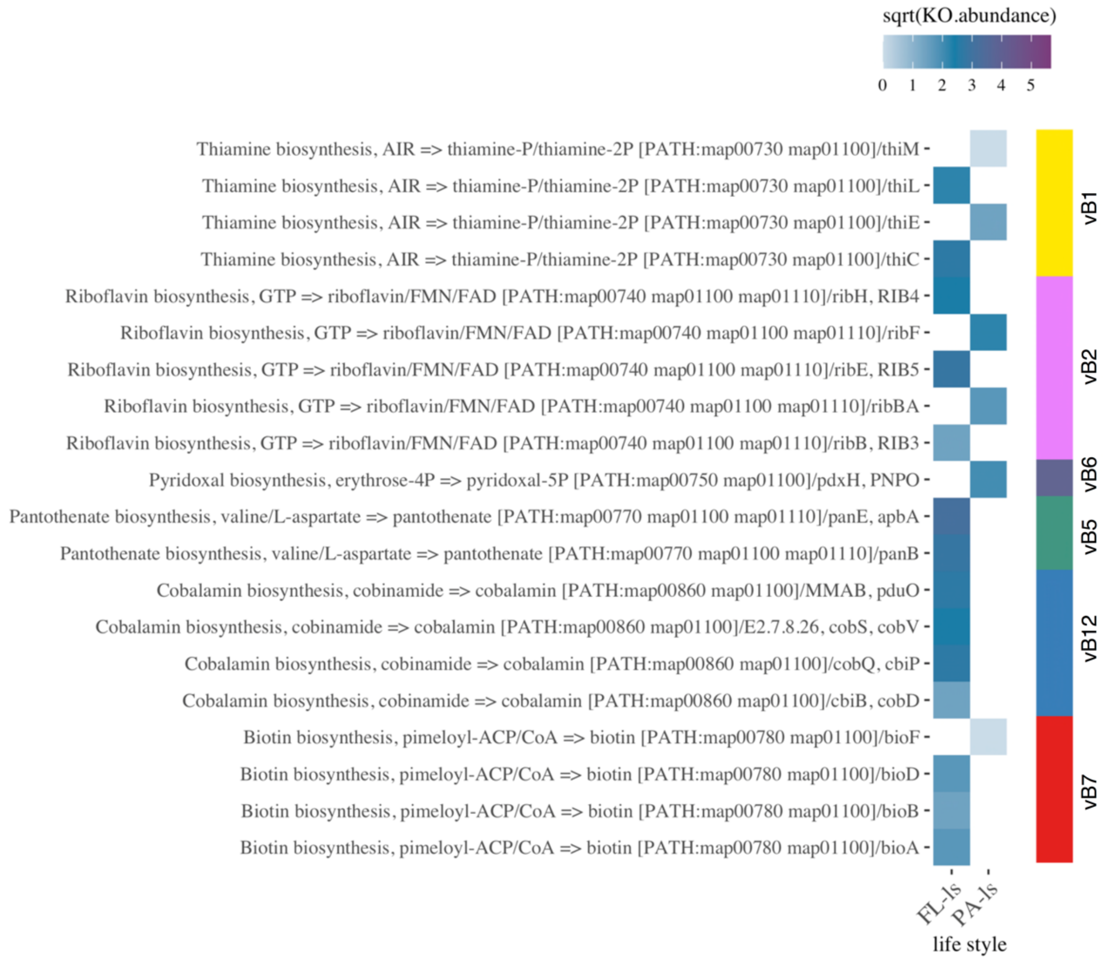


**Figure S5.** Differentially abundant genes (based on KO annotations) involved in vitamin synthesis in the FL-ls and PA-ls categories. Color bar represents the vitamin in which the different genes are involved.

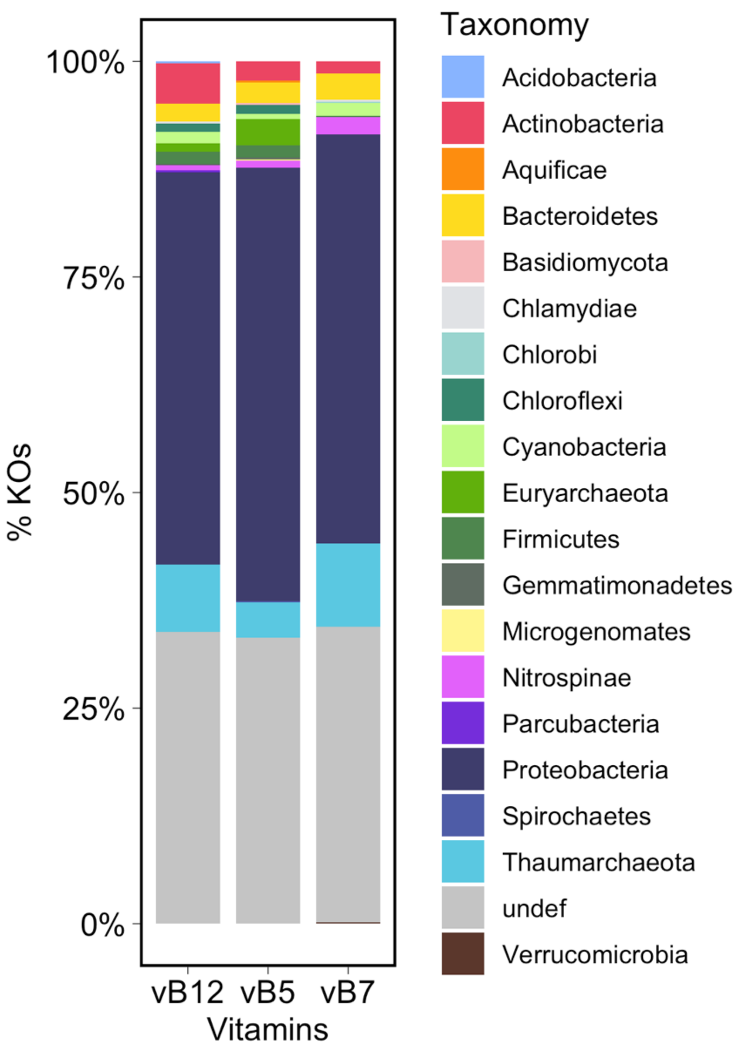


**Figure S6.** Taxonomic assignation of the vitamin synthesis genes that were preferentially found in the FL-ls genetic repertoire. The assignation was done by homology search against UniRef100 (least common ancestor).

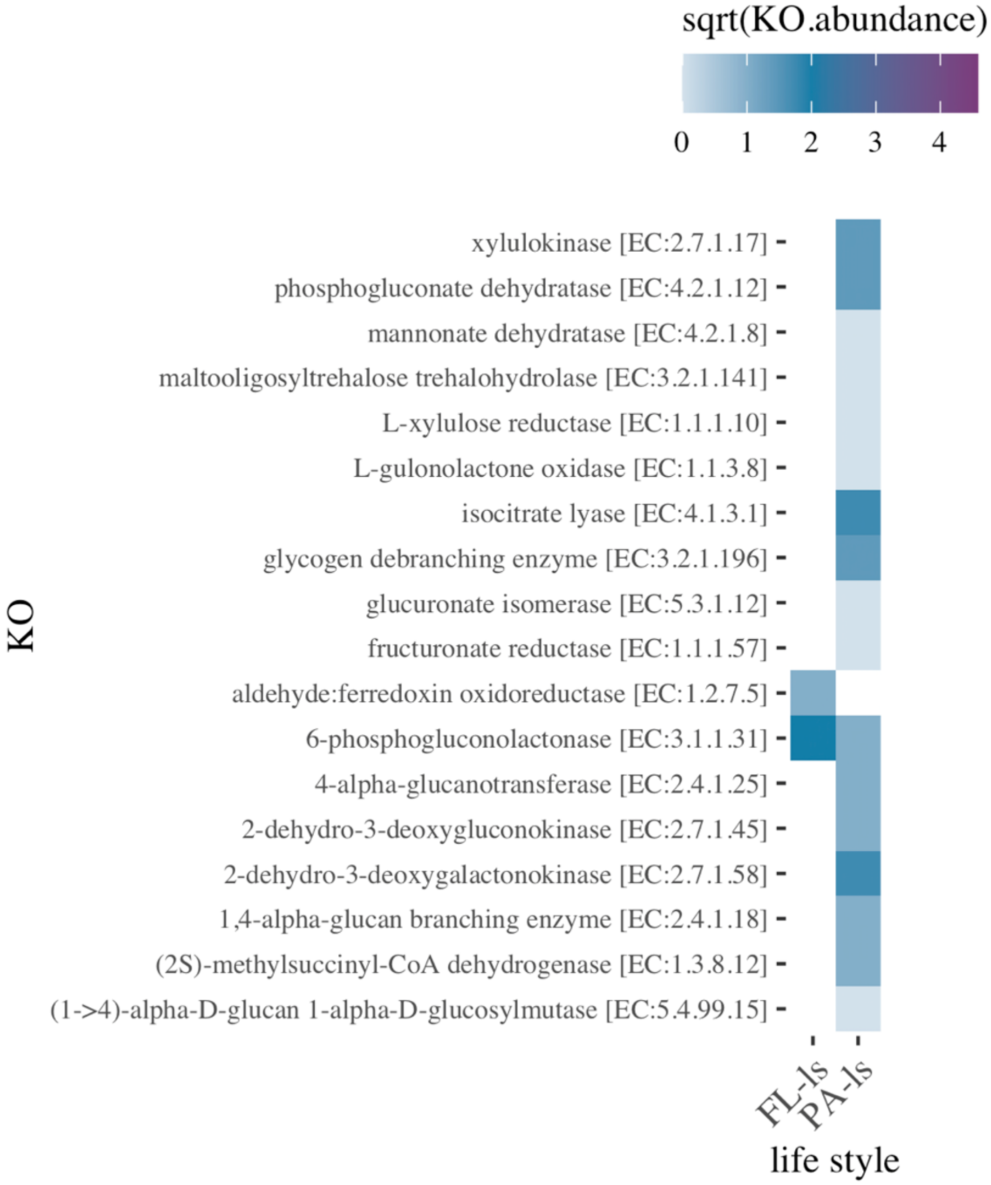


**Figure S7.** Differentially abundant genes (based on KO annotations) involved in carbohydrate metabolism in the FL-ls and PA-ls categories.

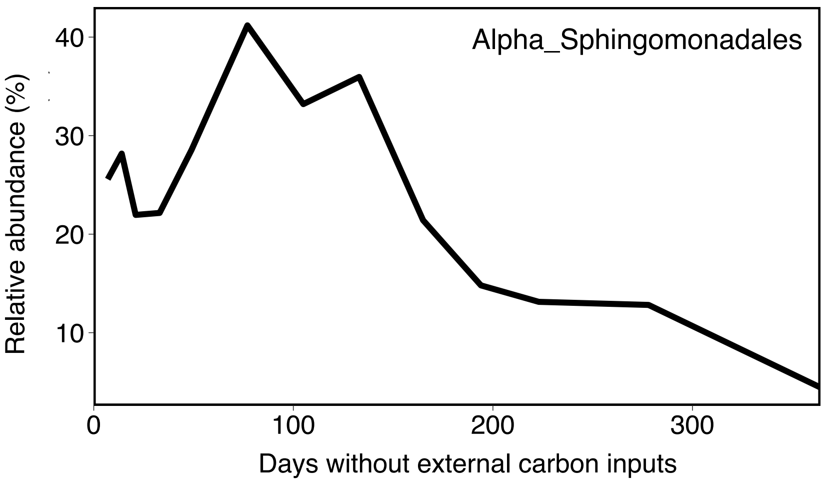


**Figure S8.** Contribution of the Alphaproteobacteria Sphingomonadales to the 16S rRNA pool of an enclosed bathypelagic community during 1 year with no external organic carbon inputs. See Sebastian et al. (2018) for more details of this experiment**.**

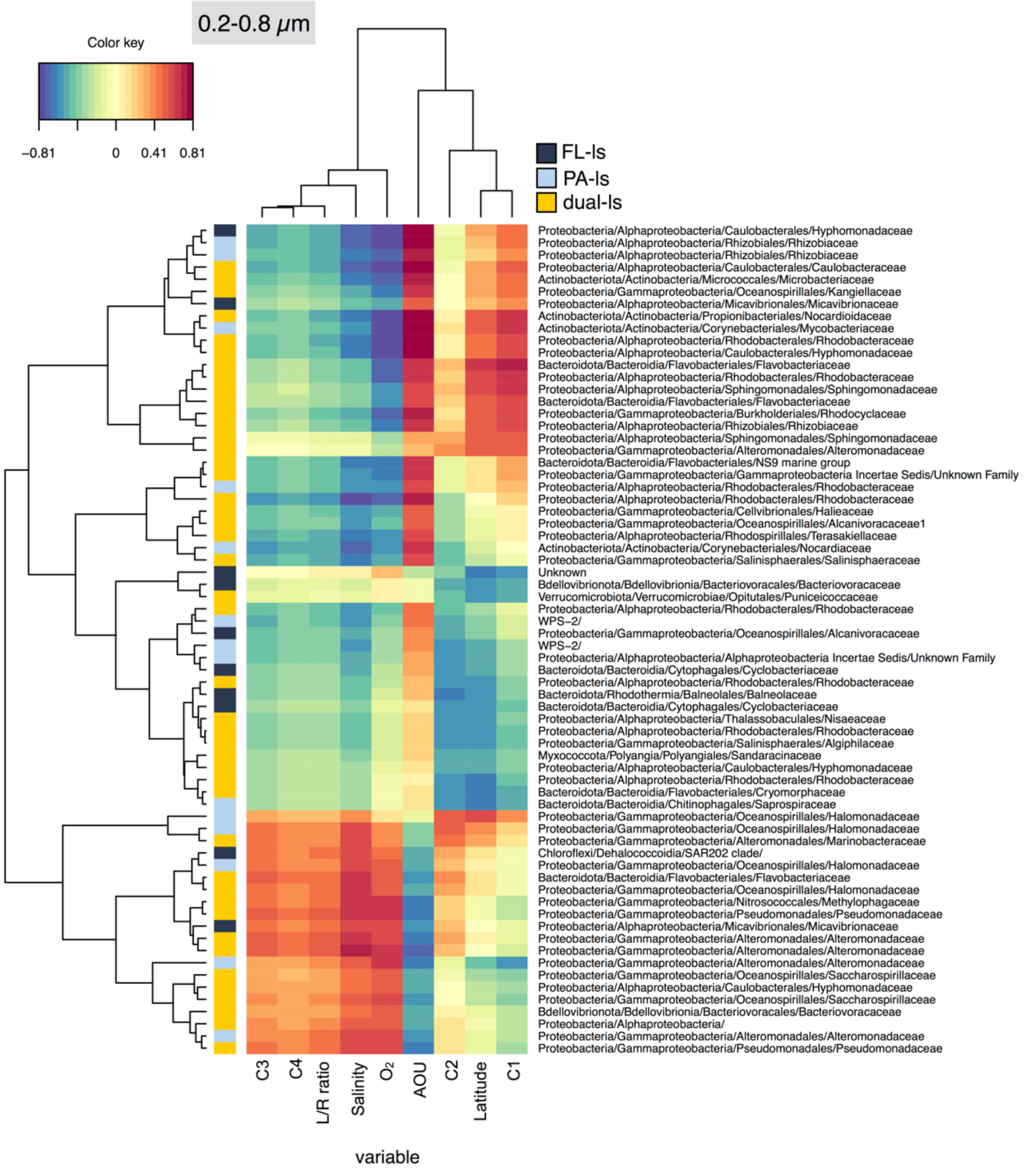


**Figure S9** Pair-wise associations between the relative abundance of individual prokaryotic ASVs in the rRNA pool of the 0.2-0.8 μm size fraction, and the absolute fluorescence of the FDOM components, the labile to recalcitrant FDOM ratio and other abiotic and biotic variables. Only those ASVs that represented >1% in relative abundance in the rRNA were considered for the analysis. Left color bar indicates the lifestyle categorization of each of the ASVs (FL-ls: free-living taxa, PA-ls, taxa with a particle associated lifestyle, and dual lifestyle taxa). Only those ASVs that displayed correlations R>0.5 with any variable are shown. Since dual-ls and PA-ls ASVs dominated the RNA pool, they represented a notable fraction of the abundant ASVs of the 0.2-0.8 μm size fraction. Interestingly FL-ls taxa appear associated to different qualities of FDOM, although in terms of percent contribution of this category to the total RNA they were overrepresented in the waters with higher values of component C1 (Figure 4, main text).

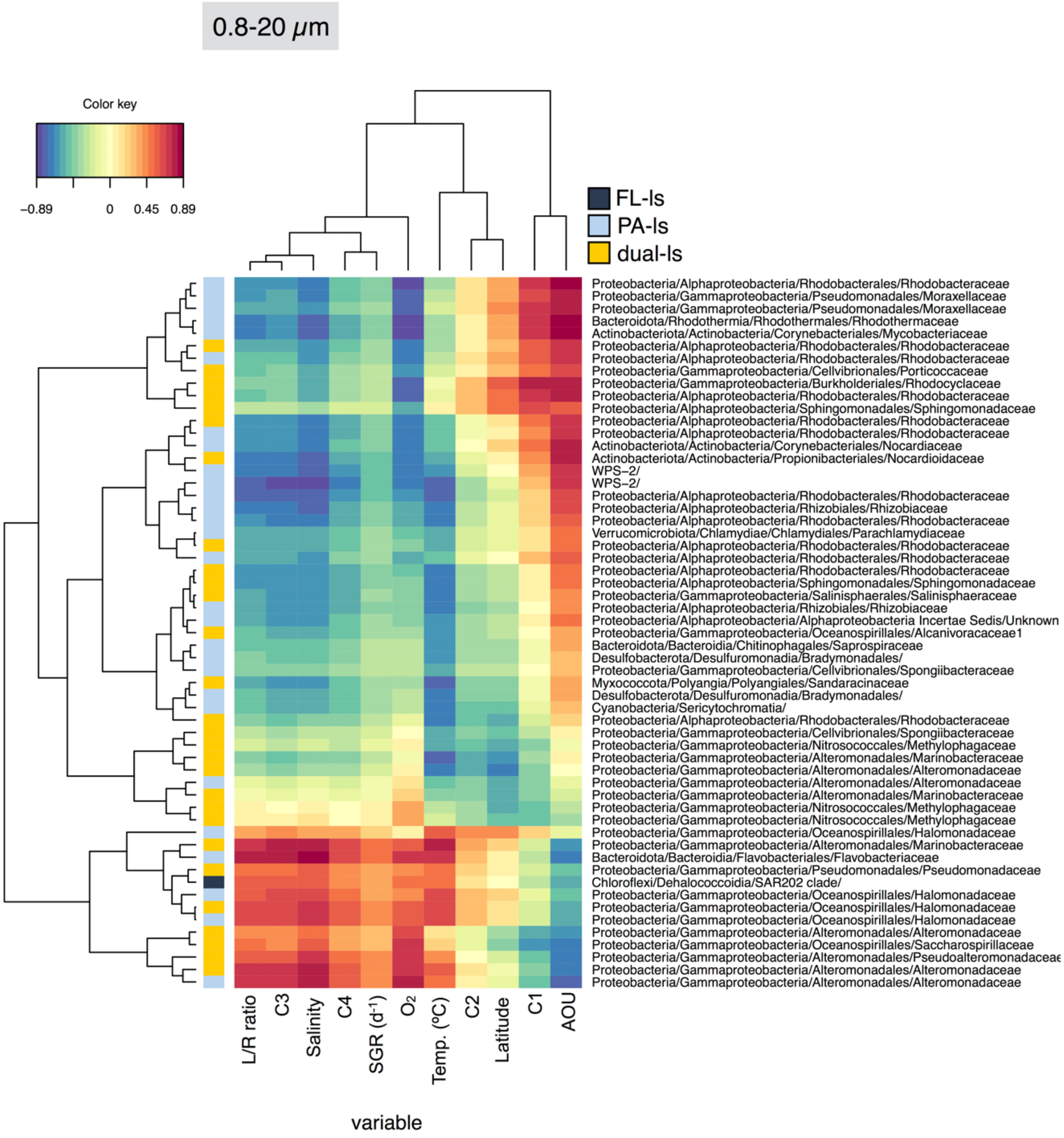


**Figure S10.** Pair-wise associations between the relative abundance of individual prokaryotic ASVs in the rRNA pool of the 0.8-20µm size fraction and the absolute fluorescence of the FDOM components, the labile to recalcitrant FDOM ratio and other abiotic and biotic variables. Only those ASVs that represented >1% in relative abundance in the rRNA were considered for the analysis. Left color bar indicates the lifestyle categorization of each of the ASVs (FL-ls: free-living taxa, PA-ls, taxa with a particle associated lifestyle, and dual lifestyle taxa). Only those ASVs that displayed correlations R>0.5 with any variable are shown.

**Table S1.** Sampling and sequencing information of the rDNA and rRNA samples for the complete (i.e. without sub-sampling) dataset.

| | | | | | | | rDNA | | rRNA | |
|---|---|---|---|---|---|---|---|---|---|---|
| station | filtersize (µm) | Longitud | Latitude | depth | date | Ocean basin | #Reads | #ASVs | #Reads | #ASVs |
| 10 | 0.8 | -26 | 14.52 | -4002 | 26/12/10 | North Atlantic | 31308 | 1053 | 153270 | 1349 |
| 10 | 0.2 | -26 | 14.52 | -4002 | 26/12/10 | North Atlantic | 14265 | 990 | 222054 | 1885 |
| 17 | 0.8 | -27.33 | -3.03 | -4002 | 2/1/11 | South Atlantic | 27697 | 1243 | 187695 | 1931 |
| 17 | 0.2 | -27.33 | -3.03 | -4002 | 2/1/11 | South Atlantic | 33865 | 1434 | 135500 | 2129 |
| 20 | 0.8 | -30.19 | -9.12 | -4001 | 5/1/11 | South Atlantic | 46782 | 1399 | 252513 | 1874 |
| 20 | 0.2 | -30.19 | -9.12 | -4001 | 5/1/11 | South Atlantic | 33591 | 1547 | 212076 | 1933 |
| 23 | 0.8 | -33.41 | -15.83 | -4003 | 8/1/11 | South Atlantic | 12406 | 1207 | | |
| 23 | 0.2 | -33.41 | -15.83 | -4003 | 8/1/11 | South Atlantic | 49030 | 1837 | 248213 | 2733 |
| 26 | 0.8 | -36.95 | -22.97 | -3907 | 11/1/11 | South Atlantic | 38489 | 1044 | 200124 | 2070 |
| 26 | 0.2 | -36.95 | -22.97 | -3907 | 11/1/11 | South Atlantic | 21748 | 1062 | 177358 | 1980 |
| 32 | 0.8 | -21.43 | -26.91 | -3199 | 24/1/11 | South Atlantic | 21673 | 942 | | |
| 32 | 0.2 | -21.43 | -26.91 | -3199 | 24/1/11 | South Atlantic | 26108 | 1380 | 172387 | 2351 |
| 35 | 0.8 | -11.8 | -28.62 | -3662 | 27/1/11 | South Atlantic | 36486 | 1373 | | |
| 35 | 0.2 | -11.8 | -28.62 | -3662 | 27/1/11 | South Atlantic | 33186 | 1646 | 282674 | 2953 |
| 41 | 0.8 | 6.84 | -31.81 | -4001 | 2/2/11 | South Atlantic | 39810 | 1459 | 174536 | 1951 |
| 41 | 0.2 | 6.84 | -31.81 | -4001 | 2/2/11 | South Atlantic | 36705 | 1552 | 215533 | 3099 |
| 43 | 0.8 | 12.7692 | -32.8128 | -3902 | 4/2/11 | South Atlantic | 32635 | 1360 | | |
| 43 | 0.2 | 12.7692 | -32.8128 | -3902 | 4/2/11 | South Atlantic | 42553 | 1331 | 221742 | 2057 |
| 50 | 0.8 | 39.89 | -33.55 | -4002 | 18/2/11 | Indian | 33044 | 961 | | |
| 50 | 0.2 | 39.89 | -33.55 | -4002 | 18/2/11 | Indian | 31117 | 1064 | 200868 | 1619 |
| 53 | 0.8 | 63.2478 | -27.9783 | -3500 | 25/2/11 | Indian | 34509 | 1284 | 229698 | 2394 |
| 53 | 0.2 | 63.2478 | -27.9783 | -3500 | 25/2/11 | Indian | 26225 | 1249 | 202008 | 2079 |
| 59 | 0.8 | 82.62 | -29.81 | -4000 | 3/3/11 | Indian | 27989 | 1537 | 226387 | 1548 |
| 59 | 0.2 | 82.62 | -29.81 | -4000 | 3/3/11 | Indian | 14613 | 1202 | 246533 | 2097 |
| 62 | 0.8 | 92.9852 | -29.6525 | -2400 | 6/3/11 | Indian | 12578 | 1096 | | |
| 62 | 0.2 | 92.9852 | -29.6525 | -2400 | 6/3/11 | Indian | 17406 | 927 | 176102 | 1272 |
| 65 | 0.8 | 103.3075 | -30.3327 | -4001 | 9/3/11 | Indian | 42244 | 1477 | | |
| 65 | 0.2 | 103.3075 | -30.3327 | -4001 | 9/3/11 | Indian | 19938 | 1233 | 222608 | 2132 |
| 67 | 0.8 | 110.18 | -31.16 | -4004 | 11/3/11 | Indian | 35743 | 1577 | | |
| 67 | 0.2 | 110.18 | -31.16 | -4004 | 11/3/11 | Indian | 49288 | 1798 | 249370 | 2467 |
| 74 | 0.8 | 135.19 | -39.23 | -3996 | 23/3/11 | Southern | 32099 | 1200 | 198692 | 1589 |
| 74 | 0.2 | 135.19 | -39.23 | -3996 | 23/3/11 | Southern | 34435 | 1381 | 246539 | 2030 |
| 77 | 0.8 | 150.41 | -38.64 | -4001 | 27/3/11 | South Pacific | 18570 | 1209 | 252081 | 2197 |
| 77 | 0.2 | 150.41 | -38.64 | -4001 | 27/3/11 | South Pacific | 17395 | 786 | 231329 | 1531 |
| 81 | 0.8 | 179.1413 | -28.4063 | -3501 | 18/4/11 | South Pacific | 37246 | 1673 | | |
| 81 | 0.2 | 179.1413 | -28.4063 | -3501 | 18/4/11 | South Pacific | 48211 | 1851 | 194688 | 2239 |
| 82 | 0.8 | -179.52 | -25.49 | -2150 | 19/4/11 | South Pacific | 54848 | 1534 | | |
| 82 | 0.2 | -179.52 | -25.49 | -2150 | 19/4/11 | South Pacific | 39898 | 1634 | 221590 | 2277 |
| 88 | 0.8 | -172.64 | -11.23 | -4001 | 25/4/11 | South Pacific | 12718 | 744 | 182654 | 1907 |
| 88 | 0.2 | -172.64 | -11.23 | -4001 | 25/4/11 | South Pacific | 44264 | 1694 | 210028 | 1925 |
| 91 | 0.8 | -170.7407 | -5.75 | -4018 | 28/4/11 | South Pacific | 13767 | 776 | | |
| 91 | 0.2 | -170.7407 | -5.75 | -4018 | 28/4/11 | South Pacific | 15063 | 1085 | | |
| 97 | 0.8 | -163.53 | 9.22 | -3818 | 4/5/11 | North Pacific | 45535 | 1270 | 239100 | 1966 |
| 97 | 0.2 | -163.53 | 9.22 | -3818 | 4/5/11 | North Pacific | 46761 | 1709 | 137889 | 1726 |
| 103 | 0.8 | -150.3192 | 21.0638 | -4013 | 16/5/11 | North Pacific | 13987 | 745 | | |
| 103 | 0.2 | -150.3192 | 21.0638 | -4013 | 16/5/11 | North Pacific | 39107 | 1517 | 178595 | 1774 |
| 109 | 0.8 | -133.26 | 18.04 | -4004 | 22/5/11 | North Pacific | 14343 | 1011 | 182226 | 1605 |
| 109 | 0.2 | -133.26 | 18.04 | -4004 | 22/5/11 | North Pacific | 16259 | 1324 | | |
| 112 | 0.8 | -124.4738 | 15.9087 | -4002 | 25/5/11 | North Pacific | 14548 | 1045 | | |
| 112 | 0.2 | -124.4738 | 15.9087 | -4002 | 25/5/11 | North Pacific | 14372 | 1152 | 211285 | 2557 |
| 118 | 0.8 | -108.06 | 12 | -3103 | 31/5/11 | North Pacific | 54444 | 1685 | 203195 | 2394 |
| 118 | 0.2 | -108.06 | 12 | -3103 | 31/5/11 | North Pacific | 15148 | 1073 | | |
| 121 | 0.8 | -99.2462 | 10.0927 | -3008 | 3/6/11 | North Pacific | 13296 | 411 | 241410 | 2382 |
| 121 | 0.2 | -99.2462 | 10.0927 | -3008 | 3/6/11 | North Pacific | 48251 | 1489 | 217633 | 2757 |
| 131 | 0.8 | -59.83 | 17.43 | -4003 | 25/6/11 | North Atlantic | 19036 | 807 | | |
| 131 | 0.2 | -59.83 | 17.43 | -4003 | 25/6/11 | North Atlantic | 17825 | 1230 | 260715 | 2275 |
| 134 | 0.8 | -52.6367 | 19.9897 | -4003 | 28/6/11 | North Atlantic | 14792 | 998 | | |
| 134 | 0.2 | -52.6367 | 19.9897 | -4003 | 28/6/11 | North Atlantic | 16489 | 1362 | 221714 | 2799 |
| 144 | 0.8 | -23.69 | 29.97 | -4003 | 8/7/11 | North Atlantic | 35701 | 1446 | | |
| 144 | 0.2 | -23.69 | 29.97 | -4003 | 8/7/11 | North Atlantic | 22929 | 1073 | 253300 | 1912 |

**Table S2.** Average FDOM properties of the different clusters of stations defined by the K-means algorithm

| **Cluster** | **Labile FDOM** (C3+C4) (R.U) | **Recalcitrant FDOM** (C1+C2) (R.U) | **L/R ratio** |
|---|---|---|---|
| Labile | 0.025 | 0.024 | 1.07 |
| Intermediate | 0.008 | 0.022 | 0.36 |
| Recalcitrant | 0.006 | 0.026 | 0.21 |

**Data S1. (separate file)**

List of genes (based on KOs annotation) that showed significant differential abundances in the 0.2-0.8 µm (labeled as Lifestyle= FL-ls) and the 0.8-20 µm (labeled as Lifestyle=PA-ls) metagenomes. The shared category includes genes that were found in both size fractions and thus represent either the genetic repertoire of dual-ls taxa or KOs shared by the FL-ls and PA-ls categories.
