## Supplementary DataS1 for "The quality of dissolved organic matter shapes the biogeography of the active bathypelagic microbiome"

| KO_id | Functional hierarchy 1 | Functional hierarchy 2 | Module_id | Module description | Name | Definition | Life_style |
|---|---|---|---|---|---|---|---|
| K00611 | Amino acid metabolism | Arginine and proline metabolism | M00029 | Urea cycle [PATH:map00220 map01230 map01100] | OTC, argF, argI | ornithine carbamoyltransferase [EC:2.1.3.3] | FL-ls |
| K00611 | Amino acid metabolism | Arginine and proline metabolism | M00844 | Arginine biosynthesis, ornithine => arginine [PATH:map0 | OTC, argF, argI | ornithine carbamoyltransferase [EC:2.1.3.3] | FL-ls |
| K00286 | Amino acid metabolism | Arginine and proline metabolism | M00015 | Proline biosynthesis, glutamate => proline [PATH:map00 | proC | pyrroline-5-carboxylate reductase [EC:1.5.1.2] | PA-ls |
| K00818 | Amino acid metabolism | Arginine and proline metabolism | M00028 | Ornithine biosynthesis, glutamate => ornithine [PATH:ma | E2.6.1.11, argD | acetylornithine aminotransferase [EC:2.6.1.11] | PA-ls |
| K00619 | Amino acid metabolism | Arginine and proline metabolism | M00028 | Ornithine biosynthesis, glutamate => ornithine [PATH:ma | argA | amino-acid N-acetyltransferase [EC:2.3.1.1] | PA-ls |
| K00620 | Amino acid metabolism | Arginine and proline metabolism | M00028 | Ornithine biosynthesis, glutamate => ornithine [PATH:ma | argJ | glutamate N-acetyltransferase / amino-acid N-acetyltran | PA-ls |
| K00930 | Amino acid metabolism | Arginine and proline metabolism | M00028 | Ornithine biosynthesis, glutamate => ornithine [PATH:ma | argB | acetylglutamate kinase [EC:2.7.2.8] | shared |
| K01438 | Amino acid metabolism | Arginine and proline metabolism | M00028 | Ornithine biosynthesis, glutamate => ornithine [PATH:ma | argE | acetylornithine deacetylase [EC:3.5.1.16] | shared |
| K14682 | Amino acid metabolism | Arginine and proline metabolism | M00028 | Ornithine biosynthesis, glutamate => ornithine [PATH:ma | argAB | amino-acid N-acetyltransferase [EC:2.3.1.1] | shared |
| K00145 | Amino acid metabolism | Arginine and proline metabolism | M00028 | Ornithine biosynthesis, glutamate => ornithine [PATH:ma | argC | N-acetyl-gamma-glutamyl-phosphate reductase [EC:1.2. | shared |
| K12659 | Amino acid metabolism | Arginine and proline metabolism | M00028 | Ornithine biosynthesis, glutamate => ornithine [PATH:ma | ARG56 | N-acetyl-gamma-glutamyl-phosphate reductase / acetylg | shared |
| K01438 | Amino acid metabolism | Arginine and proline metabolism | M00845 | Arginine biosynthesis, glutamate => acetylcitrulline => a | argE | acetylornithine deacetylase [EC:3.5.1.16] | shared |
| K00145 | Amino acid metabolism | Arginine and proline metabolism | M00845 | Arginine biosynthesis, glutamate => acetylcitrulline => a | argC | N-acetyl-gamma-glutamyl-phosphate reductase [EC:1.2. | shared |
| K09065 | Amino acid metabolism | Arginine and proline metabolism | M00845 | Arginine biosynthesis, glutamate => acetylcitrulline => a | argF | N-acetylornithine carbamoyltransferase [EC:2.1.3.9] | shared |
| K03785 | Amino acid metabolism | Aromatic amino acid metabolism | M00022 | Shikimate pathway, phosphoenolpyruvate + erythrose-4 | aroD | 3-dehydroquinate dehydratase I [EC:4.2.1.10] | FL-ls |
| K00800 | Amino acid metabolism | Aromatic amino acid metabolism | M00022 | Shikimate pathway, phosphoenolpyruvate + erythrose-4 | aroA | 3-phosphoshikimate 1-carboxyvinyltransferase [EC:2.5. | FL-ls |
| K01736 | Amino acid metabolism | Aromatic amino acid metabolism | M00022 | Shikimate pathway, phosphoenolpyruvate + erythrose-4 | aroC | chorismate synthase [EC:4.2.3.5] | FL-ls |
| K00014 | Amino acid metabolism | Aromatic amino acid metabolism | M00022 | Shikimate pathway, phosphoenolpyruvate + erythrose-4 | aroE | shikimate dehydrogenase [EC:1.1.1.25] | FL-ls |
| K00891 | Amino acid metabolism | Aromatic amino acid metabolism | M00022 | Shikimate pathway, phosphoenolpyruvate + erythrose-4 | E2.7.1.71, aroK, aro | shikimate kinase [EC:2.7.1.71] | FL-ls |
| K00766 | Amino acid metabolism | Aromatic amino acid metabolism | M00023 | Tryptophan biosynthesis, chorismate => tryptophan [PA | trpD | anthranilate phosphoribosyltransferase [EC:2.4.2.18] | FL-ls |
| K13503 | Amino acid metabolism | Aromatic amino acid metabolism | M00023 | Tryptophan biosynthesis, chorismate => tryptophan [PA | trpEG | anthranilate synthase [EC:4.1.3.27] | FL-ls |
| K01609 | Amino acid metabolism | Aromatic amino acid metabolism | M00023 | Tryptophan biosynthesis, chorismate => tryptophan [PA | trpC | indole-3-glycerol phosphate synthase [EC:4.1.1.48] | FL-ls |
| K04518 | Amino acid metabolism | Aromatic amino acid metabolism | M00024 | Phenylalanine biosynthesis, chorismate => phenylalanin | pheA2 | prephenate dehydratase [EC:4.2.1.51] | FL-ls |
| K00452 | Amino acid metabolism | Aromatic amino acid metabolism | M00038 | Tryptophan metabolism, tryptophan => kynurenine => 2- | HAAO | 3-hydroxyanthranilate 3,4-dioxygenase [EC:1.13.11.6] | FL-ls |
| K03392 | Amino acid metabolism | Aromatic amino acid metabolism | M00038 | Tryptophan metabolism, tryptophan => kynurenine => 2- | ACMSD | aminocarboxymuconate-semialdehyde decarboxylase [E | FL-ls |
| K01556 | Amino acid metabolism | Aromatic amino acid metabolism | M00038 | Tryptophan metabolism, tryptophan => kynurenine => 2- | KYNU, kynU | kynureninase [EC:3.7.1.3] | FL-ls |
| K00486 | Amino acid metabolism | Aromatic amino acid metabolism | M00038 | Tryptophan metabolism, tryptophan => kynurenine => 2- | KMO | kynurenine 3-monooxygenase [EC:1.14.13.9] | FL-ls |
| K05710 | Amino acid metabolism | Aromatic amino acid metabolism | M00545 | Trans-cinnamate degradation, trans-cinnamate => acety | hcaC | 3-phenylpropionate/trans-cinnamate dioxygenase ferred | FL-ls |
| K05708 | Amino acid metabolism | Aromatic amino acid metabolism | M00545 | Trans-cinnamate degradation, trans-cinnamate => acety | hcaE, hcaA1 | 3-phenylpropionate/trans-cinnamate dioxygenase subun | FL-ls |
| K05709 | Amino acid metabolism | Aromatic amino acid metabolism | M00545 | Trans-cinnamate degradation, trans-cinnamate => acety | hcaF, hcaA2 | 3-phenylpropionate/trans-cinnamate dioxygenase subun | FL-ls |
| K01735 | Amino acid metabolism | Aromatic amino acid metabolism | M00022 | Shikimate pathway, phosphoenolpyruvate + erythrose-4 | aroB | 3-dehydroquinate synthase [EC:4.2.3.4] | PA-ls |
| K01850 | Amino acid metabolism | Aromatic amino acid metabolism | M00024 | Phenylalanine biosynthesis, chorismate => phenylalanin | E5.4.99.5 | chorismate mutase [EC:5.4.99.5] | PA-ls |
| K04092 | Amino acid metabolism | Aromatic amino acid metabolism | M00024 | Phenylalanine biosynthesis, chorismate => phenylalanin | tyrA1 | chorismate mutase [EC:5.4.99.5] | PA-ls |
| K01850 | Amino acid metabolism | Aromatic amino acid metabolism | M00025 | Tyrosine biosynthesis, chorismate => tyrosine [PATH:m | E5.4.99.5 | chorismate mutase [EC:5.4.99.5] | PA-ls |
| K04092 | Amino acid metabolism | Aromatic amino acid metabolism | M00025 | Tyrosine biosynthesis, chorismate => tyrosine [PATH:m | tyrA1 | chorismate mutase [EC:5.4.99.5] | PA-ls |
| K01826 | Amino acid metabolism | Aromatic amino acid metabolism | M00533 | Homoprotocatechuate degradation, homoprotocatechuat | hpaF, hpcD | 5-carboxymethyl-2-hydroxymuconate isomerase [EC:5.3 | PA-ls |
| K05714 | Amino acid metabolism | Aromatic amino acid metabolism | M00545 | Trans-cinnamate degradation, trans-cinnamate => acety | mhpC | 2-hydroxy-6-oxonona-2,4-dienedioate hydrolase [EC:3.7 | PA-ls |
| K05712 | Amino acid metabolism | Aromatic amino acid metabolism | M00545 | Trans-cinnamate degradation, trans-cinnamate => acety | mhpA | 3-(3-hydroxy-phenyl)propionate hydroxylase [EC:1.14.1 | PA-ls |
| K13832 | Amino acid metabolism | Aromatic amino acid metabolism | M00022 | Shikimate pathway, phosphoenolpyruvate + erythrose-4 | aroDE, DHQ-SDH | 3-dehydroquinate dehydratase / shikimate dehydrogena | shared |
| K03786 | Amino acid metabolism | Aromatic amino acid metabolism | M00022 | Shikimate pathway, phosphoenolpyruvate + erythrose-4 | aroQ, qutE | 3-dehydroquinate dehydratase II [EC:4.2.1.10] | shared |
| K03856 | Amino acid metabolism | Aromatic amino acid metabolism | M00022 | Shikimate pathway, phosphoenolpyruvate + erythrose-4 | AROA2, aroA | 3-deoxy-7-phosphoheptulonate synthase [EC:2.5.1.54] | shared |
| K13829 | Amino acid metabolism | Aromatic amino acid metabolism | M00022 | Shikimate pathway, phosphoenolpyruvate + erythrose-4 | aroKB | shikimate kinase / 3-dehydroquinate synthase [EC:2.7.1 | shared |
| K13501 | Amino acid metabolism | Aromatic amino acid metabolism | M00023 | Tryptophan biosynthesis, chorismate => tryptophan [PA | TRP1 | anthranilate synthase / indole-3-glycerol phosphate syn | shared |
| K13497 | Amino acid metabolism | Aromatic amino acid metabolism | M00023 | Tryptophan biosynthesis, chorismate => tryptophan [PA | trpGD | anthranilate synthase/phosphoribosyltransferase [EC:4. | shared |
| K13498 | Amino acid metabolism | Aromatic amino acid metabolism | M00023 | Tryptophan biosynthesis, chorismate => tryptophan [PA | trpCF | indole-3-glycerol phosphate synthase / phosphoribosyla | shared |
| K01817 | Amino acid metabolism | Aromatic amino acid metabolism | M00023 | Tryptophan biosynthesis, chorismate => tryptophan [PA | trpF | phosphoribosylanthranilate isomerase [EC:5.3.1.24] | shared |
| K04093 | Amino acid metabolism | Aromatic amino acid metabolism | M00024 | Phenylalanine biosynthesis, chorismate => phenylalanin | pheA1 | chorismate mutase [EC:5.4.99.5] | shared |
| K04516 | Amino acid metabolism | Aromatic amino acid metabolism | M00024 | Phenylalanine biosynthesis, chorismate => phenylalanin | AROA1, aroA | chorismate mutase [EC:5.4.99.5] | shared |
| K06208 | Amino acid metabolism | Aromatic amino acid metabolism | M00024 | Phenylalanine biosynthesis, chorismate => phenylalanin | aroH | chorismate mutase [EC:5.4.99.5] | shared |
| K14170 | Amino acid metabolism | Aromatic amino acid metabolism | M00024 | Phenylalanine biosynthesis, chorismate => phenylalanin | pheA | chorismate mutase / prephenate dehydratase [EC:5.4.9 | shared |
| K01713 | Amino acid metabolism | Aromatic amino acid metabolism | M00024 | Phenylalanine biosynthesis, chorismate => phenylalanin | pheC | cyclohexadienyl dehydratase [EC:4.2.1.51 4.2.1.91] | shared |
| K04093 | Amino acid metabolism | Aromatic amino acid metabolism | M00025 | Tyrosine biosynthesis, chorismate => tyrosine [PATH:m | pheA1 | chorismate mutase [EC:5.4.99.5] | shared |
| K04516 | Amino acid metabolism | Aromatic amino acid metabolism | M00025 | Tyrosine biosynthesis, chorismate => tyrosine [PATH:m | AROA1, aroA | chorismate mutase [EC:5.4.99.5] | shared |
| K06208 | Amino acid metabolism | Aromatic amino acid metabolism | M00025 | Tyrosine biosynthesis, chorismate => tyrosine [PATH:m | aroH | chorismate mutase [EC:5.4.99.5] | shared |
| K14170 | Amino acid metabolism | Aromatic amino acid metabolism | M00025 | Tyrosine biosynthesis, chorismate => tyrosine [PATH:m | pheA | chorismate mutase / prephenate dehydratase [EC:5.4.9 | shared |
| K00543 | Amino acid metabolism | Aromatic amino acid metabolism | M00037 | Melatonin biosynthesis, tryptophan => serotonin => mela | ASMT | acetylserotonin O-methyltransferase [EC:2.1.1.4] | shared |
| K00453 | Amino acid metabolism | Aromatic amino acid metabolism | M00038 | Tryptophan metabolism, tryptophan => kynurenine => 2- | TDO2, kynA | tryptophan 2,3-dioxygenase [EC:1.13.11.11] | shared |

| | | | | | | | |
|---|---|---|---|---|---|---|---|
| K00455 | Amino acid metabolism | Aromatic amino acid metabolism | M00533 | Homoprotocatechuate degradation, homoprotocatechuat | hpaD, hpcB | 3,4-dihydroxyphenylacetate 2,3-dioxygenase [EC:1.13.1 | shared |
| K00151 | Amino acid metabolism | Aromatic amino acid metabolism | M00533 | Homoprotocatechuate degradation, homoprotocatechuat | hpaE, hpcC | 5-carboxymethyl-2-hydroxymuconic-semialdehyde dehyd | shared |
| K05921 | Amino acid metabolism | Aromatic amino acid metabolism | M00533 | Homoprotocatechuate degradation, homoprotocatechuat | hpaG | 5-oxopent-3-ene-1,2,5-tricarboxylate decarboxylase / 2- | shared |
| K05711 | Amino acid metabolism | Aromatic amino acid metabolism | M00545 | Trans-cinnamate degradation, trans-cinnamate => acety | hcaB | 2,3-dihydroxy-2,3-dihydrophenylpropionate dehydrogena | shared |
| K05713 | Amino acid metabolism | Aromatic amino acid metabolism | M00545 | Trans-cinnamate degradation, trans-cinnamate => acety | mhpB | 2,3-dihydroxyphenylpropionate 1,2-dioxygenase [EC:1.1 | shared |
| K01968 | Amino acid metabolism | Branched-chain amino acid metaboli | M00036 | Leucine degradation, leucine => acetoacetate + acetyl-C | E6.4.1.4A | 3-methylcrotonyl-CoA carboxylase alpha subunit [EC:6.4 | shared |
| K01969 | Amino acid metabolism | Branched-chain amino acid metaboli | M00036 | Leucine degradation, leucine => acetoacetate + acetyl-C | E6.4.1.4B | 3-methylcrotonyl-CoA carboxylase beta subunit [EC:6.4 | shared |
| K00253 | Amino acid metabolism | Branched-chain amino acid metaboli | M00036 | Leucine degradation, leucine => acetoacetate + acetyl-C | IVD, ivd | isovaleryl-CoA dehydrogenase [EC:1.3.8.4] | shared |
| K13766 | Amino acid metabolism | Branched-chain amino acid metaboli | M00036 | Leucine degradation, leucine => acetoacetate + acetyl-C | liuC | methylglutaconyl-CoA hydratase [EC:4.2.1.18] | shared |
| K00772 | Amino acid metabolism | Cysteine and methionine metabolism | M00034 | Methionine salvage pathway [PATH:map00270 map0110 | mtaP, MTAP | 5'-methylthioadenosine phosphorylase [EC:2.4.2.28] | FL-ls |
| K08963 | Amino acid metabolism | Cysteine and methionine metabolism | M00034 | Methionine salvage pathway [PATH:map00270 map0110 | mtnA | methylthioribose-1-phosphate isomerase [EC:5.3.1.23] | FL-ls |
| K00789 | Amino acid metabolism | Cysteine and methionine metabolism | M00034 | Methionine salvage pathway [PATH:map00270 map0110 | metK | S-adenosylmethionine synthetase [EC:2.5.1.6] | FL-ls |
| K00789 | Amino acid metabolism | Cysteine and methionine metabolism | M00035 | Methionine degradation [PATH:map00270 map01100] | metK | S-adenosylmethionine synthetase [EC:2.5.1.6] | FL-ls |
| K00789 | Amino acid metabolism | Cysteine and methionine metabolism | M00368 | Ethylene biosynthesis, methionine => ethylene [PATH:m | metK | S-adenosylmethionine synthetase [EC:2.5.1.6] | FL-ls |
| K00789 | Amino acid metabolism | Cysteine and methionine metabolism | M00609 | Cysteine biosynthesis, methionine => cysteine [PATH:m | metK | S-adenosylmethionine synthetase [EC:2.5.1.6] | FL-ls |
| K08967 | Amino acid metabolism | Cysteine and methionine metabolism | M00034 | Methionine salvage pathway [PATH:map00270 map0110 | mtnD, mtnZ, ADI1 | 1,2-dihydroxy-3-keto-5-methylthiopentene dioxygenase | shared |
| K08966 | Amino acid metabolism | Cysteine and methionine metabolism | M00034 | Methionine salvage pathway [PATH:map00270 map0110 | mtnX | 2-hydroxy-3-keto-5-methylthiopentenyl-1-phosphate pho | shared |
| K00899 | Amino acid metabolism | Cysteine and methionine metabolism | M00034 | Methionine salvage pathway [PATH:map00270 map0110 | mtnK | 5-methylthioribose kinase [EC:2.7.1.100] | shared |
| K01243 | Amino acid metabolism | Cysteine and methionine metabolism | M00034 | Methionine salvage pathway [PATH:map00270 map0110 | mtnN, mtn, pfs | adenosylhomocysteine nucleosidase [EC:3.2.2.9] | shared |
| K09880 | Amino acid metabolism | Cysteine and methionine metabolism | M00034 | Methionine salvage pathway [PATH:map00270 map0110 | mtnC, ENOPH1 | enolase-phosphatase E1 [EC:3.1.3.77] | shared |
| K08964 | Amino acid metabolism | Cysteine and methionine metabolism | M00034 | Methionine salvage pathway [PATH:map00270 map0110 | mtnB | methylthioribulose-1-phosphate dehydratase [EC:4.2.1.1 | shared |
| K01251 | Amino acid metabolism | Cysteine and methionine metabolism | M00035 | Methionine degradation [PATH:map00270 map01100] | E3.3.1.1, ahcY | adenosylhomocysteinase [EC:3.3.1.1] | shared |
| K01243 | Amino acid metabolism | Cysteine and methionine metabolism | M00609 | Cysteine biosynthesis, methionine => cysteine [PATH:m | mtnN, mtn, pfs | adenosylhomocysteine nucleosidase [EC:3.2.2.9] | shared |
| K01693 | Amino acid metabolism | Histidine metabolism | M00026 | Histidine biosynthesis, PRPP => histidine [PATH:map00 | hisB | imidazoleglycerol-phosphate dehydratase [EC:4.2.1.19] | FL-ls |
| K11755 | Amino acid metabolism | Histidine metabolism | M00026 | Histidine biosynthesis, PRPP => histidine [PATH:map00 | hisIE | phosphoribosyl-ATP pyrophosphohydrolase / phosphorib | FL-ls |
| K02502 | Amino acid metabolism | Histidine metabolism | M00026 | Histidine biosynthesis, PRPP => histidine [PATH:map00 | hisZ | ATP phosphoribosyltransferase regulatory subunit | PA-ls |
| K01523 | Amino acid metabolism | Histidine metabolism | M00026 | Histidine biosynthesis, PRPP => histidine [PATH:map00 | hisE | phosphoribosyl-ATP pyrophosphohydrolase [EC:3.6.1.31 | PA-ls |
| K01479 | Amino acid metabolism | Histidine metabolism | M00045 | Histidine degradation, histidine => N-formiminoglutamate | hutG | formiminoglutamase [EC:3.5.3.8] | PA-ls |
| K00765 | Amino acid metabolism | Histidine metabolism | M00026 | Histidine biosynthesis, PRPP => histidine [PATH:map00 | hisG | ATP phosphoribosyltransferase [EC:2.4.2.17] | shared |
| K00013 | Amino acid metabolism | Histidine metabolism | M00026 | Histidine biosynthesis, PRPP => histidine [PATH:map00 | hisD | histidinol dehydrogenase [EC:1.1.1.23] | shared |
| K04486 | Amino acid metabolism | Histidine metabolism | M00026 | Histidine biosynthesis, PRPP => histidine [PATH:map00 | E3.1.3.15B | histidinol-phosphatase (PHP family) [EC:3.1.3.15] | shared |
| K05602 | Amino acid metabolism | Histidine metabolism | M00026 | Histidine biosynthesis, PRPP => histidine [PATH:map00 | hisN | histidinol-phosphatase [EC:3.1.3.15] | shared |
| K02500 | Amino acid metabolism | Histidine metabolism | M00026 | Histidine biosynthesis, PRPP => histidine [PATH:map00 | hisF | imidazole glycerol-phosphate synthase subunit HisF [EC | shared |
| K02501 | Amino acid metabolism | Histidine metabolism | M00026 | Histidine biosynthesis, PRPP => histidine [PATH:map00 | hisH | imidazole glycerol-phosphate synthase subunit HisH [EC | shared |
| K01089 | Amino acid metabolism | Histidine metabolism | M00026 | Histidine biosynthesis, PRPP => histidine [PATH:map00 | hisB | imidazoleglycerol-phosphate dehydratase / histidinol-ph | shared |
| K01496 | Amino acid metabolism | Histidine metabolism | M00026 | Histidine biosynthesis, PRPP => histidine [PATH:map00 | hisI | phosphoribosyl-AMP cyclohydrolase [EC:3.5.4.19] | shared |
| K14152 | Amino acid metabolism | Histidine metabolism | M00026 | Histidine biosynthesis, PRPP => histidine [PATH:map00 | HIS4 | phosphoribosyl-ATP pyrophosphohydrolase / phosphorib | shared |
| K01814 | Amino acid metabolism | Histidine metabolism | M00026 | Histidine biosynthesis, PRPP => histidine [PATH:map00 | hisA | phosphoribosylformimino-5-aminoimidazole carboxamide | shared |
| K05603 | Amino acid metabolism | Histidine metabolism | M00045 | Histidine degradation, histidine => N-formiminoglutamate | hutF | formimidoylglutamate deiminase [EC:3.5.3.13] | shared |
| K01745 | Amino acid metabolism | Histidine metabolism | M00045 | Histidine degradation, histidine => N-formiminoglutamate | hutH, HAL | histidine ammonia-lyase [EC:4.3.1.3] | shared |
| K01468 | Amino acid metabolism | Histidine metabolism | M00045 | Histidine degradation, histidine => N-formiminoglutamate | hutI, AMDHD1 | imidazolonepropionase [EC:3.5.2.7] | shared |
| K01712 | Amino acid metabolism | Histidine metabolism | M00045 | Histidine degradation, histidine => N-formiminoglutamate | hutU, UROC1 | urocanate hydratase [EC:4.2.1.49] | shared |
| K05827 | Amino acid metabolism | Lysine metabolism | M00031 | Lysine biosynthesis, mediated by LysW, 2-aminoadipate | lysX | [lysine-biosynthesis-protein LysW]---L-2-aminoadipate li | FL-ls |
| K10206 | Amino acid metabolism | Lysine metabolism | M00527 | Lysine biosynthesis, DAP aminotransferase pathway, as | E2.6.1.83 | LL-diaminopimelate aminotransferase [EC:2.6.1.83] | FL-ls |
| K00674 | Amino acid metabolism | Lysine metabolism | M00016 | Lysine biosynthesis, succinyl-DAP pathway, aspartate = | dapD | 2,3,4,5-tetrahydropyridine-2,6-dicarboxylate N-succinylt | PA-ls |
| K01586 | Amino acid metabolism | Lysine metabolism | M00016 | Lysine biosynthesis, succinyl-DAP pathway, aspartate = | lysA | diaminopimelate decarboxylase [EC:4.1.1.20] | PA-ls |
| K01778 | Amino acid metabolism | Lysine metabolism | M00016 | Lysine biosynthesis, succinyl-DAP pathway, aspartate = | dapF | diaminopimelate epimerase [EC:5.1.1.7] | PA-ls |
| K14267 | Amino acid metabolism | Lysine metabolism | M00016 | Lysine biosynthesis, succinyl-DAP pathway, aspartate = | dapC | N-succinyldiaminopimelate aminotransferase [EC:2.6.1.1 | PA-ls |
| K01439 | Amino acid metabolism | Lysine metabolism | M00016 | Lysine biosynthesis, succinyl-DAP pathway, aspartate = | dapE | succinyl-diaminopimelate desuccinylase [EC:3.5.1.18] | PA-ls |
| K01586 | Amino acid metabolism | Lysine metabolism | M00525 | Lysine biosynthesis, acetyl-DAP pathway, aspartate => | lysA | diaminopimelate decarboxylase [EC:4.1.1.20] | PA-ls |
| K01778 | Amino acid metabolism | Lysine metabolism | M00525 | Lysine biosynthesis, acetyl-DAP pathway, aspartate => | dapF | diaminopimelate epimerase [EC:5.1.1.7] | PA-ls |
| K01586 | Amino acid metabolism | Lysine metabolism | M00526 | Lysine biosynthesis, DAP dehydrogenase pathway, asp | lysA | diaminopimelate decarboxylase [EC:4.1.1.20] | PA-ls |
| K01586 | Amino acid metabolism | Lysine metabolism | M00527 | Lysine biosynthesis, DAP aminotransferase pathway, as | lysA | diaminopimelate decarboxylase [EC:4.1.1.20] | PA-ls |
| K01778 | Amino acid metabolism | Lysine metabolism | M00527 | Lysine biosynthesis, DAP aminotransferase pathway, as | dapF | diaminopimelate epimerase [EC:5.1.1.7] | PA-ls |
| K01705 | Amino acid metabolism | Lysine metabolism | M00030 | Lysine biosynthesis, AAA pathway, 2-oxoglutarate => 2- | LYS4 | homoaconitate hydratase [EC:4.2.1.36] | shared |
| K14157 | Amino acid metabolism | Lysine metabolism | M00032 | Lysine degradation, lysine => saccharopine => acetoac | AASS | alpha-aminoadipic semialdehyde synthase [EC:1.5.1.8 1 | shared |
| K01705 | Amino acid metabolism | Lysine metabolism | M00433 | Lysine biosynthesis, 2-oxoglutarate => 2-oxoadipate [PA | LYS4 | homoaconitate hydratase [EC:4.2.1.36] | shared |
| K03340 | Amino acid metabolism | Lysine metabolism | M00526 | Lysine biosynthesis, DAP dehydrogenase pathway, asp | dapdh | diaminopimelate dehydrogenase [EC:1.4.1.16] | shared |
| K01480 | Amino acid metabolism | Polyamine biosynthesis | M00133 | Polyamine biosynthesis, arginine => agmatine => putres | speB | agmatinase [EC:3.5.3.11] | FL-ls |
| K02626 | Amino acid metabolism | Polyamine biosynthesis | M00133 | Polyamine biosynthesis, arginine => agmatine => putres | pdaD | arginine decarboxylase [EC:4.1.1.19] | FL-ls |

| | | | | | | | |
|---|---|---|---|---|---|---|---|
| K09472 | Amino acid metabolism | Polyamine biosynthesis | M00136 | GABA biosynthesis, prokaryotes, putrescine => GABA [ | puuC, aldH | 4-(gamma-glutamylamino)butanal dehydrogenase [EC:1. | PA-ls |
| K01584 | Amino acid metabolism | Polyamine biosynthesis | M00133 | Polyamine biosynthesis, arginine => agmatine => putres | adiA | arginine decarboxylase [EC:4.1.1.19] | shared |
| K01585 | Amino acid metabolism | Polyamine biosynthesis | M00133 | Polyamine biosynthesis, arginine => agmatine => putres | speA | arginine decarboxylase [EC:4.1.1.19] | shared |
| K00657 | Amino acid metabolism | Polyamine biosynthesis | M00135 | GABA biosynthesis, eukaryotes, putrescine => GABA [ | speG, SAT | diamine N-acetyltransferase [EC:2.3.1.57] | shared |
| K09471 | Amino acid metabolism | Polyamine biosynthesis | M00136 | GABA biosynthesis, prokaryotes, putrescine => GABA [ | puuB, ordL | gamma-glutamylputrescine oxidase [EC:1.4.3.-] | shared |
| K09470 | Amino acid metabolism | Polyamine biosynthesis | M00136 | GABA biosynthesis, prokaryotes, putrescine => GABA [ | puuA | gamma-glutamylputrescine synthase [EC:6.3.1.11] | shared |
| K00872 | Amino acid metabolism | Serine and threonine metabolism | M00018 | Threonine biosynthesis, aspartate => homoserine => thr | thrB1 | homoserine kinase [EC:2.7.1.39] | FL-ls |
| K02204 | Amino acid metabolism | Serine and threonine metabolism | M00018 | Threonine biosynthesis, aspartate => homoserine => thr | thrB2 | homoserine kinase type II [EC:2.7.1.39] | PA-ls |
| K00836 | Amino acid metabolism | Serine and threonine metabolism | M00033 | Ectoine biosynthesis, aspartate => ectoine [PATH:map0 | ectB, dat | diaminobutyrate-2-oxoglutarate transaminase [EC:2.6.1. | PA-ls |
| K06718 | Amino acid metabolism | Serine and threonine metabolism | M00033 | Ectoine biosynthesis, aspartate => ectoine [PATH:map0 | ectA | L-2,4-diaminobutyric acid acetyltransferase [EC:2.3.1.1 | PA-ls |
| K06720 | Amino acid metabolism | Serine and threonine metabolism | M00033 | Ectoine biosynthesis, aspartate => ectoine [PATH:map0 | ectC | L-ectoine synthase [EC:4.2.1.108] | PA-ls |
| K00130 | Amino acid metabolism | Serine and threonine metabolism | M00555 | Betaine biosynthesis, choline => betaine [PATH:map002 | betB, gbsA | betaine-aldehyde dehydrogenase [EC:1.2.1.8] | PA-ls |
| K00108 | Amino acid metabolism | Serine and threonine metabolism | M00555 | Betaine biosynthesis, choline => betaine [PATH:map002 | betA, CHDH | choline dehydrogenase [EC:1.1.99.1] | shared |
| K04127 | Biosynthesis of other sec | beta-Lactam biosynthesis | M00673 | Cephamycin C biosynthesis, aminoadipate + cycteine + | cefD | isopenicillin-N epimerase [EC:5.1.1.17] | FL-ls |
| K05375 | Biosynthesis of other sec | beta-Lactam biosynthesis | M00736 | Nocardicin A biosynthesis, L-pHPG + arginine + serine = | mbtH, nocI | MbtH protein | PA-ls |
| K10852 | Biosynthesis of other sec | beta-Lactam biosynthesis | M00672 | Penicillin biosynthesis, aminoadipate + cycteine + valine | PENDE | isopenicillin-N N-acyltransferase [EC:2.3.1.164] | shared |
| K04126 | Biosynthesis of other sec | beta-Lactam biosynthesis | M00672 | Penicillin biosynthesis, aminoadipate + cycteine + valine | PCBC | isopenicillin-N synthase [EC:1.21.3.1] | shared |
| K04126 | Biosynthesis of other sec | beta-Lactam biosynthesis | M00673 | Cephamycin C biosynthesis, aminoadipate + cycteine + | PCBC | isopenicillin-N synthase [EC:1.21.3.1] | shared |
| K13063 | Biosynthesis of other sec | Biosynthesis of other secondary me | M00835 | Pyocyanine biosynthesis, chorismate => pyocyanine [P. | phzE | 2-amino-4-deoxychorismate synthase [EC:2.6.1.86] | PA-ls |
| K06998 | Biosynthesis of other sec | Biosynthesis of other secondary me | M00835 | Pyocyanine biosynthesis, chorismate => pyocyanine [P. | phzF | trans-2,3-dihydro-3-hydroxyanthranilate isomerase [EC: | PA-ls |
| K14266 | Biosynthesis of other sec | Biosynthesis of other secondary me | M00789 | Rebeccamycin biosynthesis, tryptophan => rebeccamyc | prnA, rebH, ktzQ | tryptophan 7-halogenase [EC:1.14.19.9] | shared |
| K14266 | Biosynthesis of other sec | Biosynthesis of other secondary me | M00790 | Pyrrolnitrin biosynthesis, tryptophan => pyrrolnitrin [PAT | prnA, rebH, ktzQ | tryptophan 7-halogenase [EC:1.14.19.9] | shared |
| K02291 | Biosynthesis of terpenoid | Other terpenoid biosynthesis | M00097 | beta-Carotene biosynthesis, GGAP => beta-carotene [P | crtB | 15-cis-phytoene synthase [EC:2.5.1.32] | shared |
| K06443 | Biosynthesis of terpenoid | Other terpenoid biosynthesis | M00097 | beta-Carotene biosynthesis, GGAP => beta-carotene [P | lcyB, crtL1, crtY | lycopene beta-cyclase [EC:5.5.1.19] | shared |
| K15746 | Biosynthesis of terpenoid | Other terpenoid biosynthesis | M00372 | Abscisic acid biosynthesis, beta-carotene => abscisic a | crtZ | beta-carotene 3-hydroxylase [EC:1.14.15.24] | shared |
| K01823 | Biosynthesis of terpenoid | Terpenoid backbone biosynthesis | M00095 | C5 isoprenoid biosynthesis, mevalonate pathway [PATH | idi, IDI | isopentenyl-diphosphate Delta-isomerase [EC:5.3.3.2] | FL-ls |
| K01823 | Biosynthesis of terpenoid | Terpenoid backbone biosynthesis | M00096 | C5 isoprenoid biosynthesis, non-mevalonate pathway [P | idi, IDI | isopentenyl-diphosphate Delta-isomerase [EC:5.3.3.2] | FL-ls |
| K13787 | Biosynthesis of terpenoid | Terpenoid backbone biosynthesis | M00364 | C10-C20 isoprenoid biosynthesis, bacteria [PATH:map0 | idsA | geranylgeranyl diphosphate synthase, type I [EC:2.5.1. | FL-ls |
| K13789 | Biosynthesis of terpenoid | Terpenoid backbone biosynthesis | M00364 | C10-C20 isoprenoid biosynthesis, bacteria [PATH:map0 | GGPS | geranylgeranyl diphosphate synthase, type II [EC:2.5.1 | FL-ls |
| K01823 | Biosynthesis of terpenoid | Terpenoid backbone biosynthesis | M00364 | C10-C20 isoprenoid biosynthesis, bacteria [PATH:map0 | idi, IDI | isopentenyl-diphosphate Delta-isomerase [EC:5.3.3.2] | FL-ls |
| K13787 | Biosynthesis of terpenoid | Terpenoid backbone biosynthesis | M00365 | C10-C20 isoprenoid biosynthesis, archaea [PATH:map0 | idsA | geranylgeranyl diphosphate synthase, type I [EC:2.5.1. | FL-ls |
| K01823 | Biosynthesis of terpenoid | Terpenoid backbone biosynthesis | M00365 | C10-C20 isoprenoid biosynthesis, archaea [PATH:map0 | idi, IDI | isopentenyl-diphosphate Delta-isomerase [EC:5.3.3.2] | FL-ls |
| K13789 | Biosynthesis of terpenoid | Terpenoid backbone biosynthesis | M00366 | C10-C20 isoprenoid biosynthesis, plants [PATH:map009 | GGPS | geranylgeranyl diphosphate synthase, type II [EC:2.5.1 | FL-ls |
| K01823 | Biosynthesis of terpenoid | Terpenoid backbone biosynthesis | M00366 | C10-C20 isoprenoid biosynthesis, plants [PATH:map009 | idi, IDI | isopentenyl-diphosphate Delta-isomerase [EC:5.3.3.2] | FL-ls |
| K01823 | Biosynthesis of terpenoid | Terpenoid backbone biosynthesis | M00367 | C10-C20 isoprenoid biosynthesis, non-plant eukaryotes | idi, IDI | isopentenyl-diphosphate Delta-isomerase [EC:5.3.3.2] | FL-ls |
| K00054 | Biosynthesis of terpenoid | Terpenoid backbone biosynthesis | M00849 | C5 isoprenoid biosynthesis, mevalonate pathway, archa | mvaA | hydroxymethylglutaryl-CoA reductase [EC:1.1.1.88] | FL-ls |
| K06981 | Biosynthesis of terpenoid | Terpenoid backbone biosynthesis | M00849 | C5 isoprenoid biosynthesis, mevalonate pathway, archa | ipk | isopentenyl phosphate kinase [EC:2.7.4.26] | FL-ls |
| K01823 | Biosynthesis of terpenoid | Terpenoid backbone biosynthesis | M00849 | C5 isoprenoid biosynthesis, mevalonate pathway, archa | idi, IDI | isopentenyl-diphosphate Delta-isomerase [EC:5.3.3.2] | FL-ls |
| K03526 | Biosynthesis of terpenoid | Terpenoid backbone biosynthesis | M00096 | C5 isoprenoid biosynthesis, non-mevalonate pathway [P | gcpE, ispG | (E)-4-hydroxy-3-methylbut-2-enyl-diphosphate synthase | PA-ls |
| K12506 | Biosynthesis of terpenoid | Terpenoid backbone biosynthesis | M00096 | C5 isoprenoid biosynthesis, non-mevalonate pathway [P | ispDF | 2-C-methyl-D-erythritol 4-phosphate cytidylyltransferase | PA-ls |
| K00919 | Biosynthesis of terpenoid | Terpenoid backbone biosynthesis | M00096 | C5 isoprenoid biosynthesis, non-mevalonate pathway [P | ispE | 4-diphosphocytidyl-2-C-methyl-D-erythritol kinase [EC:2. | PA-ls |
| K03527 | Biosynthesis of terpenoid | Terpenoid backbone biosynthesis | M00096 | C5 isoprenoid biosynthesis, non-mevalonate pathway [P | ispH, lytB | 4-hydroxy-3-methylbut-2-en-1-yl diphosphate reductase | PA-ls |
| K00795 | Biosynthesis of terpenoid | Terpenoid backbone biosynthesis | M00364 | C10-C20 isoprenoid biosynthesis, bacteria [PATH:map0 | ispA | farnesyl diphosphate synthase [EC:2.5.1.1 2.5.1.10] | PA-ls |
| K01597 | Biosynthesis of terpenoid | Terpenoid backbone biosynthesis | M00095 | C5 isoprenoid biosynthesis, mevalonate pathway [PATH | MVD, mvaD | diphosphomevalonate decarboxylase [EC:4.1.1.33] | shared |
| K00938 | Biosynthesis of terpenoid | Terpenoid backbone biosynthesis | M00095 | C5 isoprenoid biosynthesis, mevalonate pathway [PATH | E2.7.4.2, mvaK2 | phosphomevalonate kinase [EC:2.7.4.2] | shared |
| K00099 | Biosynthesis of terpenoid | Terpenoid backbone biosynthesis | M00096 | C5 isoprenoid biosynthesis, non-mevalonate pathway [P | dxr | 1-deoxy-D-xylulose-5-phosphate reductoisomerase [EC: | shared |
| K01770 | Biosynthesis of terpenoid | Terpenoid backbone biosynthesis | M00096 | C5 isoprenoid biosynthesis, non-mevalonate pathway [P | ispF | 2-C-methyl-D-erythritol 2,4-cyclodiphosphate synthase [ | shared |
| K00991 | Biosynthesis of terpenoid | Terpenoid backbone biosynthesis | M00096 | C5 isoprenoid biosynthesis, non-mevalonate pathway [P | ispD | 2-C-methyl-D-erythritol 4-phosphate cytidylyltransferase | shared |
| K05554 | Biosynthesis of terpenoid | Type II polyketide biosynthesis | M00778 | Type II polyketide backbone biosynthesis, acyl-CoA + n | actVII | aromatase [EC:4.2.1.-] | PA-ls |
| K05555 | Biosynthesis of terpenoid | Type II polyketide biosynthesis | M00778 | Type II polyketide backbone biosynthesis, acyl-CoA + n | actIV | cyclase [EC:4.-.-.-] | shared |
| K14630 | Biosynthesis of terpenoid | Type II polyketide biosynthesis | M00779 | Dihydrokalafungin biosynthesis, octaketide => dihydroka | actVA5 | two-component flavin-dependent monooxygenase [EC:1 | shared |
| K07404 | Carbohydrate metabolism | Central carbohydrate metabolism | M00004 | Pentose phosphate pathway (Pentose phosphate cycle) | pgl | 6-phosphogluconolactonase [EC:3.1.1.31] | FL-ls |
| K07404 | Carbohydrate metabolism | Central carbohydrate metabolism | M00006 | Pentose phosphate pathway, oxidative phase, glucose | pgl | 6-phosphogluconolactonase [EC:3.1.1.31] | FL-ls |
| K07404 | Carbohydrate metabolism | Central carbohydrate metabolism | M00008 | Entner-Doudoroff pathway, glucose-6P => glyceraldehy | pgl | 6-phosphogluconolactonase [EC:3.1.1.31] | FL-ls |
| K03738 | Carbohydrate metabolism | Central carbohydrate metabolism | M00309 | Non-phosphorylative Entner-Doudoroff pathway, glucona | aor | aldehyde:ferredoxin oxidoreductase [EC:1.2.7.5] | FL-ls |
| K01057 | Carbohydrate metabolism | Central carbohydrate metabolism | M00004 | Pentose phosphate pathway (Pentose phosphate cycle) | PGLS, pgl, devB | 6-phosphogluconolactonase [EC:3.1.1.31] | PA-ls |
| K01057 | Carbohydrate metabolism | Central carbohydrate metabolism | M00006 | Pentose phosphate pathway, oxidative phase, glucose | PGLS, pgl, devB | 6-phosphogluconolactonase [EC:3.1.1.31] | PA-ls |
| K01057 | Carbohydrate metabolism | Central carbohydrate metabolism | M00008 | Entner-Doudoroff pathway, glucose-6P => glyceraldehy | PGLS, pgl, devB | 6-phosphogluconolactonase [EC:3.1.1.31] | PA-ls |
| K01690 | Carbohydrate metabolism | Central carbohydrate metabolism | M00008 | Entner-Doudoroff pathway, glucose-6P => glyceraldehy | edd | phosphogluconate dehydratase [EC:4.2.1.12] | PA-ls |
| K00874 | Carbohydrate metabolism | Central carbohydrate metabolism | M00308 | Semi-phosphorylative Entner-Doudoroff pathway, glucon | kdgK | 2-dehydro-3-deoxygluconokinase [EC:2.7.1.45] | PA-ls |

| | | | | | | | |
|---|---|---|---|---|---|---|---|
| K13937 | Carbohydrate metabolism | Central carbohydrate metabolism | M00004 | Pentose phosphate pathway (Pentose phosphate cycle) | H6PD | hexose-6-phosphate dehydrogenase [EC:1.1.1.47 3.1.1 | shared |
| K00616 | Carbohydrate metabolism | Central carbohydrate metabolism | M00004 | Pentose phosphate pathway (Pentose phosphate cycle) | E2.2.1.2, talA, talB | transaldolase [EC:2.2.1.2] | shared |
| K13937 | Carbohydrate metabolism | Central carbohydrate metabolism | M00006 | Pentose phosphate pathway, oxidative phase, glucose | H6PD | hexose-6-phosphate dehydrogenase [EC:1.1.1.47 3.1.1 | shared |
| K00616 | Carbohydrate metabolism | Central carbohydrate metabolism | M00007 | Pentose phosphate pathway, non-oxidative phase, fruct | E2.2.1.2, talA, talB | transaldolase [EC:2.2.1.2] | shared |
| K00030 | Carbohydrate metabolism | Central carbohydrate metabolism | M00009 | Citrate cycle (TCA cycle, Krebs cycle) [PATH:map0002 | IDH3 | isocitrate dehydrogenase (NAD+) [EC:1.1.1.41] | shared |
| K00030 | Carbohydrate metabolism | Central carbohydrate metabolism | M00010 | Citrate cycle, first carbon oxidation, oxaloacetate => 2- | IDH3 | isocitrate dehydrogenase (NAD+) [EC:1.1.1.41] | shared |
| K01637 | Carbohydrate metabolism | Other carbohydrate metabolism | M00012 | Glyoxylate cycle [PATH:map00630 map01200 map0110 | E4.1.3.1, aceA | isocitrate lyase [EC:4.1.3.1] | PA-ls |
| K03331 | Carbohydrate metabolism | Other carbohydrate metabolism | M00014 | Glucuronate pathway (uronate pathway) [PATH:map0004 | DCXR | L-xylulose reductase [EC:1.1.1.10] | PA-ls |
| K00854 | Carbohydrate metabolism | Other carbohydrate metabolism | M00014 | Glucuronate pathway (uronate pathway) [PATH:map0004 | xylB, XYLB | xylulokinase [EC:2.7.1.17] | PA-ls |
| K00874 | Carbohydrate metabolism | Other carbohydrate metabolism | M00061 | D-Glucuronate degradation, D-glucuronate => pyruvate | kdgK | 2-dehydro-3-deoxygluconokinase [EC:2.7.1.45] | PA-ls |
| K00040 | Carbohydrate metabolism | Other carbohydrate metabolism | M00061 | D-Glucuronate degradation, D-glucuronate => pyruvate | uxuB | fructuronate reductase [EC:1.1.1.57] | PA-ls |
| K01812 | Carbohydrate metabolism | Other carbohydrate metabolism | M00061 | D-Glucuronate degradation, D-glucuronate => pyruvate | uxaC | glucuronate isomerase [EC:5.3.1.12] | PA-ls |
| K01686 | Carbohydrate metabolism | Other carbohydrate metabolism | M00061 | D-Glucuronate degradation, D-glucuronate => pyruvate | uxuA | mannonate dehydratase [EC:4.2.1.8] | PA-ls |
| K08323 | Carbohydrate metabolism | Other carbohydrate metabolism | M00061 | D-Glucuronate degradation, D-glucuronate => pyruvate | rspA, manD | mannonate dehydratase [EC:4.2.1.8] | PA-ls |
| K00103 | Carbohydrate metabolism | Other carbohydrate metabolism | M00129 | Ascorbate biosynthesis, animals, glucose-1P => ascorb | GULO | L-gulonolactone oxidase [EC:1.1.3.8] | PA-ls |
| K14448 | Carbohydrate metabolism | Other carbohydrate metabolism | M00373 | Ethylmalonyl pathway [PATH:map00630 map01200 map0 | mcd | (2S)-methylsuccinyl-CoA dehydrogenase [EC:1.3.8.12] | PA-ls |
| K00883 | Carbohydrate metabolism | Other carbohydrate metabolism | M00552 | D-galactonate degradation, De Ley-Doudoroff pathway, | dgoK | 2-dehydro-3-deoxygalactonokinase [EC:2.7.1.58] | PA-ls |
| K06044 | Carbohydrate metabolism | Other carbohydrate metabolism | M00565 | Trehalose biosynthesis, D-glucose 1P => trehalose [PA | treY, glgY | (1->4)-alpha-D-glucan 1-alpha-D-glucosylmutase [EC:5.4 | PA-ls |
| K00700 | Carbohydrate metabolism | Other carbohydrate metabolism | M00565 | Trehalose biosynthesis, D-glucose 1P => trehalose [PA | GBE1, glgB | 1,4-alpha-glucan branching enzyme [EC:2.4.1.18] | PA-ls |
| K01236 | Carbohydrate metabolism | Other carbohydrate metabolism | M00565 | Trehalose biosynthesis, D-glucose 1P => trehalose [PA | treZ, glgZ | maltooligosyltrehalose trehalohydrolase [EC:3.2.1.141] | PA-ls |
| K00874 | Carbohydrate metabolism | Other carbohydrate metabolism | M00631 | D-Galacturonate degradation (bacteria), D-galacturonate | kdgK | 2-dehydro-3-deoxygluconokinase [EC:2.7.1.45] | PA-ls |
| K01812 | Carbohydrate metabolism | Other carbohydrate metabolism | M00631 | D-Galacturonate degradation (bacteria), D-galacturonate | uxaC | glucuronate isomerase [EC:5.3.1.12] | PA-ls |
| K00700 | Carbohydrate metabolism | Other carbohydrate metabolism | M00854 | Glycogen biosynthesis, glucose-1P => glycogen/starch | GBE1, glgB | 1,4-alpha-glucan branching enzyme [EC:2.4.1.18] | PA-ls |
| K00705 | Carbohydrate metabolism | Other carbohydrate metabolism | M00855 | Glycogen degradation, glycogen => glucose-6P [PATH: | malQ | 4-alpha-glucanotransferase [EC:2.4.1.25] | PA-ls |
| K02438 | Carbohydrate metabolism | Other carbohydrate metabolism | M00855 | Glycogen degradation, glycogen => glucose-6P [PATH: | glgX | glycogen debranching enzyme [EC:3.2.1.196] | PA-ls |
| K05351 | Carbohydrate metabolism | Other carbohydrate metabolism | M00014 | Glucuronate pathway (uronate pathway) [PATH:map0004 | E1.1.1.9 | D-xylulose reductase [EC:1.1.1.9] | shared |
| K14451 | Carbohydrate metabolism | Other carbohydrate metabolism | M00373 | Ethylmalonyl pathway [PATH:map00630 map01200 map0 | mcl2 | (3S)-malyl-CoA thioesterase [EC:3.1.2.30] | shared |
| K14446 | Carbohydrate metabolism | Other carbohydrate metabolism | M00373 | Ethylmalonyl pathway [PATH:map00630 map01200 map0 | ccr | crotonyl-CoA carboxylase/reductase [EC:1.3.1.85] | shared |
| K14447 | Carbohydrate metabolism | Other carbohydrate metabolism | M00373 | Ethylmalonyl pathway [PATH:map00630 map01200 map0 | ecm | ethylmalonyl-CoA mutase [EC:5.4.99.63] | shared |
| K01631 | Carbohydrate metabolism | Other carbohydrate metabolism | M00552 | D-galactonate degradation, De Ley-Doudoroff pathway, | dgoA | 2-dehydro-3-deoxyphosphogalactonate aldolase [EC:4.1 | shared |
| K01684 | Carbohydrate metabolism | Other carbohydrate metabolism | M00552 | D-galactonate degradation, De Ley-Doudoroff pathway, | dgoD | galactonate dehydratase [EC:4.2.1.6] | shared |
| K16149 | Carbohydrate metabolism | Other carbohydrate metabolism | M00565 | Trehalose biosynthesis, D-glucose 1P => trehalose [PA | K16149 | 1,4-alpha-glucan branching enzyme [EC:2.4.1.18] | shared |
| K00703 | Carbohydrate metabolism | Other carbohydrate metabolism | M00565 | Trehalose biosynthesis, D-glucose 1P => trehalose [PA | glgA | starch synthase [EC:2.4.1.21] | shared |
| K01685 | Carbohydrate metabolism | Other carbohydrate metabolism | M00631 | D-Galacturonate degradation (bacteria), D-galacturonate | uxaA | altronate hydrolase [EC:4.2.1.7] | shared |
| K00041 | Carbohydrate metabolism | Other carbohydrate metabolism | M00631 | D-Galacturonate degradation (bacteria), D-galacturonate | uxaB | tagaturonate reductase [EC:1.1.1.58] | shared |
| K16149 | Carbohydrate metabolism | Other carbohydrate metabolism | M00854 | Glycogen biosynthesis, glucose-1P => glycogen/starch | K16149 | 1,4-alpha-glucan branching enzyme [EC:2.4.1.18] | shared |
| K16150 | Carbohydrate metabolism | Other carbohydrate metabolism | M00854 | Glycogen biosynthesis, glucose-1P => glycogen/starch | K16150 | glycogen synthase [EC:2.4.1.11] | shared |
| K00703 | Carbohydrate metabolism | Other carbohydrate metabolism | M00854 | Glycogen biosynthesis, glucose-1P => glycogen/starch | glgA | starch synthase [EC:2.4.1.21] | shared |
| K00330 | Energy metabolism | ATP synthesis | M00144 | NADH:quinone oxidoreductase, prokaryotes [PATH:map | nuoA | NADH-quinone oxidoreductase subunit A [EC:7.1.1.2] | FL-ls |
| K00331 | Energy metabolism | ATP synthesis | M00144 | NADH:quinone oxidoreductase, prokaryotes [PATH:map | nuoB | NADH-quinone oxidoreductase subunit B [EC:7.1.1.2] | FL-ls |
| K00332 | Energy metabolism | ATP synthesis | M00144 | NADH:quinone oxidoreductase, prokaryotes [PATH:map | nuoC | NADH-quinone oxidoreductase subunit C [EC:7.1.1.2] | FL-ls |
| K00333 | Energy metabolism | ATP synthesis | M00144 | NADH:quinone oxidoreductase, prokaryotes [PATH:map | nuoD | NADH-quinone oxidoreductase subunit D [EC:7.1.1.2] | FL-ls |
| K00337 | Energy metabolism | ATP synthesis | M00144 | NADH:quinone oxidoreductase, prokaryotes [PATH:map | nuoH | NADH-quinone oxidoreductase subunit H [EC:7.1.1.2] | FL-ls |
| K00338 | Energy metabolism | ATP synthesis | M00144 | NADH:quinone oxidoreductase, prokaryotes [PATH:map | nuoI | NADH-quinone oxidoreductase subunit I [EC:7.1.1.2] | FL-ls |
| K00340 | Energy metabolism | ATP synthesis | M00144 | NADH:quinone oxidoreductase, prokaryotes [PATH:map | nuoK | NADH-quinone oxidoreductase subunit K [EC:7.1.1.2] | FL-ls |
| K00341 | Energy metabolism | ATP synthesis | M00144 | NADH:quinone oxidoreductase, prokaryotes [PATH:map | nuoL | NADH-quinone oxidoreductase subunit L [EC:7.1.1.2] | FL-ls |
| K00342 | Energy metabolism | ATP synthesis | M00144 | NADH:quinone oxidoreductase, prokaryotes [PATH:map | nuoM | NADH-quinone oxidoreductase subunit M [EC:7.1.1.2] | FL-ls |
| K00343 | Energy metabolism | ATP synthesis | M00144 | NADH:quinone oxidoreductase, prokaryotes [PATH:map | nuoN | NADH-quinone oxidoreductase subunit N [EC:7.1.1.2] | FL-ls |
| K05573 | Energy metabolism | ATP synthesis | M00145 | NAD(P)H:quinone oxidoreductase, chloroplasts and cya | ndhB | NAD(P)H-quinone oxidoreductase subunit 2 [EC:7.1.1.2] | FL-ls |
| K05578 | Energy metabolism | ATP synthesis | M00145 | NAD(P)H:quinone oxidoreductase, chloroplasts and cya | ndhG | NAD(P)H-quinone oxidoreductase subunit 6 [EC:7.1.1.2] | FL-ls |
| K02277 | Energy metabolism | ATP synthesis | M00155 | Cytochrome c oxidase, prokaryotes [PATH:map00190] | coxD, ctaF | cytochrome c oxidase subunit IV [EC:1.9.3.1] | FL-ls |
| K02117 | Energy metabolism | ATP synthesis | M00159 | V-type ATPase, prokaryotes [PATH:map00190] | ATPVA, ntpA, atpA | V/A-type H+/Na+-transporting ATPase subunit A [EC:7.1 | FL-ls |
| K02118 | Energy metabolism | ATP synthesis | M00159 | V-type ATPase, prokaryotes [PATH:map00190] | ATPVB, ntpB, atpB | V/A-type H+/Na+-transporting ATPase subunit B | FL-ls |
| K02119 | Energy metabolism | ATP synthesis | M00159 | V-type ATPase, prokaryotes [PATH:map00190] | ATPVC, ntpC, atpC | V/A-type H+/Na+-transporting ATPase subunit C | FL-ls |
| K02120 | Energy metabolism | ATP synthesis | M00159 | V-type ATPase, prokaryotes [PATH:map00190] | ATPVD, ntpD, atpD | V/A-type H+/Na+-transporting ATPase subunit D | FL-ls |
| K02121 | Energy metabolism | ATP synthesis | M00159 | V-type ATPase, prokaryotes [PATH:map00190] | ATPVE, ntpE, atpE | V/A-type H+/Na+-transporting ATPase subunit E | FL-ls |
| K02122 | Energy metabolism | ATP synthesis | M00159 | V-type ATPase, prokaryotes [PATH:map00190] | ATPVF, ntpF, atpF | V/A-type H+/Na+-transporting ATPase subunit F | FL-ls |
| K02123 | Energy metabolism | ATP synthesis | M00159 | V-type ATPase, prokaryotes [PATH:map00190] | ATPVI, ntpI, atpI | V/A-type H+/Na+-transporting ATPase subunit I | FL-ls |
| K02124 | Energy metabolism | ATP synthesis | M00159 | V-type ATPase, prokaryotes [PATH:map00190] | ATPVK, ntpK, atpK | V/A-type H+/Na+-transporting ATPase subunit K | FL-ls |
| K02636 | Energy metabolism | ATP synthesis | M00162 | Cytochrome b6f complex [PATH:map00195] | petC | cytochrome b6-f complex iron-sulfur subunit [EC:7.1.1.6 | FL-ls |

| | | | | | | | |
|---|---|---|---|---|---|---|---|
| K00334 | Energy metabolism | ATP synthesis | M00144 | NADH:quinone oxidoreductase, prokaryotes [PATH:map( | nuoE | NADH-quinone oxidoreductase subunit E [EC:7.1.1.2] | PA-ls |
| K00336 | Energy metabolism | ATP synthesis | M00144 | NADH:quinone oxidoreductase, prokaryotes [PATH:map( | nuoG | NADH-quinone oxidoreductase subunit G [EC:7.1.1.2] | PA-ls |
| K03891 | Energy metabolism | ATP synthesis | M00151 | Cytochrome bc1 complex respiratory unit [PATH:map001 | qcrB | ubiquinol-cytochrome c reductase cytochrome b subunit | PA-ls |
| K03889 | Energy metabolism | ATP synthesis | M00151 | Cytochrome bc1 complex respiratory unit [PATH:map001 | qcrC | ubiquinol-cytochrome c reductase cytochrome c subunit | PA-ls |
| K03890 | Energy metabolism | ATP synthesis | M00151 | Cytochrome bc1 complex respiratory unit [PATH:map001 | qcrA | ubiquinol-cytochrome c reductase iron-sulfur subunit | PA-ls |
| K02276 | Energy metabolism | ATP synthesis | M00155 | Cytochrome c oxidase, prokaryotes [PATH:map00190] | coxC, ctaE | cytochrome c oxidase subunit III [EC:1.9.3.1] | PA-ls |
| K02297 | Energy metabolism | ATP synthesis | M00417 | Cytochrome o ubiquinol oxidase [PATH:map00190] | cyoA | cytochrome o ubiquinol oxidase subunit II [EC:7.1.1.3] | PA-ls |
| K13378 | Energy metabolism | ATP synthesis | M00144 | NADH:quinone oxidoreductase, prokaryotes [PATH:map( | nuoCD | NADH-quinone oxidoreductase subunit C/D [EC:7.1.1.2] | shared |
| K00335 | Energy metabolism | ATP synthesis | M00144 | NADH:quinone oxidoreductase, prokaryotes [PATH:map( | nuoF | NADH-quinone oxidoreductase subunit F [EC:7.1.1.2] | shared |
| K00339 | Energy metabolism | ATP synthesis | M00144 | NADH:quinone oxidoreductase, prokaryotes [PATH:map( | nuoJ | NADH-quinone oxidoreductase subunit J [EC:7.1.1.2] | shared |
| K05572 | Energy metabolism | ATP synthesis | M00145 | NAD(P)H:quinone oxidoreductase, chloroplasts and cya | ndhA | NAD(P)H-quinone oxidoreductase subunit 1 [EC:7.1.1.2] | shared |
| K05574 | Energy metabolism | ATP synthesis | M00145 | NAD(P)H:quinone oxidoreductase, chloroplasts and cya | ndhC | NAD(P)H-quinone oxidoreductase subunit 3 [EC:7.1.1.2] | shared |
| K05575 | Energy metabolism | ATP synthesis | M00145 | NAD(P)H:quinone oxidoreductase, chloroplasts and cya | ndhD | NAD(P)H-quinone oxidoreductase subunit 4 [EC:7.1.1.2] | shared |
| K05576 | Energy metabolism | ATP synthesis | M00145 | NAD(P)H:quinone oxidoreductase, chloroplasts and cya | ndhE | NAD(P)H-quinone oxidoreductase subunit 4L [EC:7.1.1.2 | shared |
| K05577 | Energy metabolism | ATP synthesis | M00145 | NAD(P)H:quinone oxidoreductase, chloroplasts and cya | ndhF | NAD(P)H-quinone oxidoreductase subunit 5 [EC:7.1.1.2] | shared |
| K05579 | Energy metabolism | ATP synthesis | M00145 | NAD(P)H:quinone oxidoreductase, chloroplasts and cya | ndhH | NAD(P)H-quinone oxidoreductase subunit H [EC:7.1.1.2] | shared |
| K05580 | Energy metabolism | ATP synthesis | M00145 | NAD(P)H:quinone oxidoreductase, chloroplasts and cya | ndhI | NAD(P)H-quinone oxidoreductase subunit I [EC:7.1.1.2] | shared |
| K05581 | Energy metabolism | ATP synthesis | M00145 | NAD(P)H:quinone oxidoreductase, chloroplasts and cya | ndhJ | NAD(P)H-quinone oxidoreductase subunit J [EC:7.1.1.2] | shared |
| K03886 | Energy metabolism | ATP synthesis | M00151 | Cytochrome bc1 complex respiratory unit [PATH:map001 | MQCRA, qcrA, bfcA | menaquinol-cytochrome c reductase iron-sulfur subunit | shared |
| K02258 | Energy metabolism | ATP synthesis | M00154 | Cytochrome c oxidase [PATH:map00190] | COX11, ctaG | cytochrome c oxidase assembly protein subunit 11 | shared |
| K02274 | Energy metabolism | ATP synthesis | M00155 | Cytochrome c oxidase, prokaryotes [PATH:map00190] | coxA, ctaD | cytochrome c oxidase subunit I [EC:1.9.3.1] | shared |
| K15408 | Energy metabolism | ATP synthesis | M00155 | Cytochrome c oxidase, prokaryotes [PATH:map00190] | coxAC | cytochrome c oxidase subunit I+III [EC:1.9.3.1] | shared |
| K02275 | Energy metabolism | ATP synthesis | M00155 | Cytochrome c oxidase, prokaryotes [PATH:map00190] | coxB, ctaC | cytochrome c oxidase subunit II [EC:1.9.3.1] | shared |
| K02827 | Energy metabolism | ATP synthesis | M00416 | Cytochrome aa3-600 menaquinol oxidase [PATH:map001 | qoxB | cytochrome aa3-600 menaquinol oxidase subunit I [EC:7 | shared |
| K02826 | Energy metabolism | ATP synthesis | M00416 | Cytochrome aa3-600 menaquinol oxidase [PATH:map001 | qoxA | cytochrome aa3-600 menaquinol oxidase subunit II [EC: | shared |
| K02298 | Energy metabolism | ATP synthesis | M00417 | Cytochrome o ubiquinol oxidase [PATH:map00190] | cyoB | cytochrome o ubiquinol oxidase subunit I [EC:7.1.1.3] | shared |
| K02299 | Energy metabolism | ATP synthesis | M00417 | Cytochrome o ubiquinol oxidase [PATH:map00190] | cyoC | cytochrome o ubiquinol oxidase subunit III | shared |
| K02300 | Energy metabolism | ATP synthesis | M00417 | Cytochrome o ubiquinol oxidase [PATH:map00190] | cyoD | cytochrome o ubiquinol oxidase subunit IV | shared |
| K15016 | Energy metabolism | Carbon fixation | M00374 | Dicarboxylate-hydroxybutyrate cycle [PATH:map00720 r | K15016 | enoyl-CoA hydratase / 3-hydroxyacyl-CoA dehydrogena | FL-ls |
| K15019 | Energy metabolism | Carbon fixation | M00375 | Hydroxypropionate-hydroxybutylate cycle [PATH:map00 | K15019 | 3-hydroxypropionyl-coenzyme A dehydratase [EC:4.2.1. | FL-ls |
| K15016 | Energy metabolism | Carbon fixation | M00375 | Hydroxypropionate-hydroxybutylate cycle [PATH:map00 | K15016 | enoyl-CoA hydratase / 3-hydroxyacyl-CoA dehydrogena | FL-ls |
| K00855 | Energy metabolism | Carbon fixation | M00165 | Reductive pentose phosphate cycle (Calvin cycle) [PAT | PRK, prkB | phosphoribulokinase [EC:2.7.1.19] | shared |
| K00855 | Energy metabolism | Carbon fixation | M00166 | Reductive pentose phosphate cycle, ribulose-5P => glyc | PRK, prkB | phosphoribulokinase [EC:2.7.1.19] | shared |
| K15020 | Energy metabolism | Carbon fixation | M00375 | Hydroxypropionate-hydroxybutylate cycle [PATH:map00 | K15020 | acryloyl-coenzyme A reductase [EC:1.3.1.84] | shared |
| K14470 | Energy metabolism | Carbon fixation | M00376 | 3-Hydroxypropionate bi-cycle [PATH:map00720 map012 | mct | 2-methylfumaryl-CoA isomerase [EC:5.4.1.3] | shared |
| K09709 | Energy metabolism | Carbon fixation | M00376 | 3-Hydroxypropionate bi-cycle [PATH:map00720 map012 | meh | 3-methylfumaryl-CoA hydratase [EC:4.2.1.153] | shared |
| K03389 | Energy metabolism | Methane metabolism | M00356 | Methanogenesis, methanol => methane [PATH:map0068 | hdrB2 | heterodisulfide reductase subunit B2 [EC:1.8.7.3 1.8.98 | FL-ls |
| K03389 | Energy metabolism | Methane metabolism | M00357 | Methanogenesis, acetate => methane [PATH:map00680 | hdrB2 | heterodisulfide reductase subunit B2 [EC:1.8.7.3 1.8.98 | FL-ls |
| K05884 | Energy metabolism | Methane metabolism | M00358 | Coenzyme M biosynthesis [PATH:map00680 map01120] | comC | L-2-hydroxycarboxylate dehydrogenase (NAD+) [EC:1.1 | FL-ls |
| K06034 | Energy metabolism | Methane metabolism | M00358 | Coenzyme M biosynthesis [PATH:map00680 map01120] | comD | sulfopyruvate decarboxylase subunit alpha [EC:4.1.1.79 | FL-ls |
| K13039 | Energy metabolism | Methane metabolism | M00358 | Coenzyme M biosynthesis [PATH:map00680 map01120] | comE | sulfopyruvate decarboxylase subunit beta [EC:4.1.1.79] | FL-ls |
| K14941 | Energy metabolism | Methane metabolism | M00378 | F420 biosynthesis [PATH:map00680 map01120] | cofC | 2-phospho-L-lactate guanylyltransferase [EC:2.7.7.68] | FL-ls |
| K11781 | Energy metabolism | Methane metabolism | M00378 | F420 biosynthesis [PATH:map00680 map01120] | cofH | 5-amino-6-(D-ribitylamino)uracil---L-tyrosine 4-hydroxyph | FL-ls |
| K11780 | Energy metabolism | Methane metabolism | M00378 | F420 biosynthesis [PATH:map00680 map01120] | cofG | 7,8-didemethyl-8-hydroxy-5-deazariboflavin synthase [E | FL-ls |
| K12234 | Energy metabolism | Methane metabolism | M00378 | F420 biosynthesis [PATH:map00680 map01120] | cofE, fbiB | coenzyme F420-0:L-glutamate ligase / coenzyme F420-1 | FL-ls |
| K11212 | Energy metabolism | Methane metabolism | M00378 | F420 biosynthesis [PATH:map00680 map01120] | cofD | LPPG:FO 2-phospho-L-lactate transferase [EC:2.7.8.28] | FL-ls |
| K03389 | Energy metabolism | Methane metabolism | M00563 | Methanogenesis, methylamine/dimethylamine/trimethylar | hdrB2 | heterodisulfide reductase subunit B2 [EC:1.8.7.3 1.8.98 | FL-ls |
| K14083 | Energy metabolism | Methane metabolism | M00563 | Methanogenesis, methylamine/dimethylamine/trimethylar | mttB | trimethylamine---corrinoid protein Co-methyltransferase [ | FL-ls |
| K03389 | Energy metabolism | Methane metabolism | M00567 | Methanogenesis, CO2 => methane [PATH:map00680 ma | hdrB2 | heterodisulfide reductase subunit B2 [EC:1.8.7.3 1.8.98 | FL-ls |
| K03388 | Energy metabolism | Methane metabolism | M00356 | Methanogenesis, methanol => methane [PATH:map0068 | hdrA2 | heterodisulfide reductase subunit A2 [EC:1.8.7.3 1.8.98 | shared |
| K03388 | Energy metabolism | Methane metabolism | M00357 | Methanogenesis, acetate => methane [PATH:map00680 | hdrA2 | heterodisulfide reductase subunit A2 [EC:1.8.7.3 1.8.98 | shared |
| K05979 | Energy metabolism | Methane metabolism | M00358 | Coenzyme M biosynthesis [PATH:map00680 map01120] | comB | 2-phosphosulfolactate phosphatase [EC:3.1.3.71] | shared |
| K08097 | Energy metabolism | Methane metabolism | M00358 | Coenzyme M biosynthesis [PATH:map00680 map01120] | comA | phosphosulfolactate synthase [EC:4.4.1.19] | shared |
| K11779 | Energy metabolism | Methane metabolism | M00378 | F420 biosynthesis [PATH:map00680 map01120] | fbiC | FO synthase [EC:2.5.1.147 4.3.1.32] | shared |
| K03388 | Energy metabolism | Methane metabolism | M00563 | Methanogenesis, methylamine/dimethylamine/trimethylar | hdrA2 | heterodisulfide reductase subunit A2 [EC:1.8.7.3 1.8.98 | shared |
| K00205 | Energy metabolism | Methane metabolism | M00567 | Methanogenesis, CO2 => methane [PATH:map00680 ma | fwdF, fmdF | 4Fe-4S ferredoxin | shared |
| K00320 | Energy metabolism | Methane metabolism | M00567 | Methanogenesis, CO2 => methane [PATH:map00680 ma | mer | 5,10-methylenetetrahydromethanopterin reductase [EC:1 | shared |
| K00200 | Energy metabolism | Methane metabolism | M00567 | Methanogenesis, CO2 => methane [PATH:map00680 ma | fwdA, fmdA | formylmethanofuran dehydrogenase subunit A [EC:1.2.7 | shared |
| K00201 | Energy metabolism | Methane metabolism | M00567 | Methanogenesis, CO2 => methane [PATH:map00680 ma | fwdB, fmdB | formylmethanofuran dehydrogenase subunit B [EC:1.2.7 | shared |
| K00202 | Energy metabolism | Methane metabolism | M00567 | Methanogenesis, CO2 => methane [PATH:map00680 ma | fwdC, fmdC | formylmethanofuran dehydrogenase subunit C [EC:1.2.7 | shared |

| | | | | | | | |
|---|---|---|---|---|---|---|---|
| K00672 | Energy metabolism | Methane metabolism | M00567 | Methanogenesis, CO2 => methane [PATH:map00680 ma | ftr | formylmethanofuran--tetrahydromethanopterin N-formyltr | shared |
| K03388 | Energy metabolism | Methane metabolism | M00567 | Methanogenesis, CO2 => methane [PATH:map00680 ma | hdrA2 | heterodisulfide reductase subunit A2 [EC:1.8.7.3 1.8.98 | shared |
| K01499 | Energy metabolism | Methane metabolism | M00567 | Methanogenesis, CO2 => methane [PATH:map00680 ma | mch | methenyltetrahydromethanopterin cyclohydrolase [EC:3. | shared |
| K00368 | Energy metabolism | Nitrogen metabolism | M00529 | Denitrification, nitrate => nitrogen [PATH:map00910 map | nirK | nitrite reductase (NO-forming) [EC:1.7.2.1] | FL-ls |
| K00362 | Energy metabolism | Nitrogen metabolism | M00530 | Dissimilatory nitrate reduction, nitrate => ammonia [PAT | nirB | nitrite reductase (NADH) large subunit [EC:1.7.1.15] | PA-ls |
| K00363 | Energy metabolism | Nitrogen metabolism | M00530 | Dissimilatory nitrate reduction, nitrate => ammonia [PAT | nirD | nitrite reductase (NADH) small subunit [EC:1.7.1.15] | PA-ls |
| K00372 | Energy metabolism | Nitrogen metabolism | M00531 | Assimilatory nitrate reduction, nitrate => ammonia [PATH | nasA | assimilatory nitrate reductase catalytic subunit [EC:1.7. | PA-ls |
| K00366 | Energy metabolism | Nitrogen metabolism | M00531 | Assimilatory nitrate reduction, nitrate => ammonia [PATH | nirA | ferredoxin-nitrite reductase [EC:1.7.7.1] | PA-ls |
| K02568 | Energy metabolism | Nitrogen metabolism | M00529 | Denitrification, nitrate => nitrogen [PATH:map00910 map | napB | cytochrome c-type protein NapB | shared |
| K04561 | Energy metabolism | Nitrogen metabolism | M00529 | Denitrification, nitrate => nitrogen [PATH:map00910 map | norB | nitric oxide reductase subunit B [EC:1.7.2.5] | shared |
| K02305 | Energy metabolism | Nitrogen metabolism | M00529 | Denitrification, nitrate => nitrogen [PATH:map00910 map | norC | nitric oxide reductase subunit C | shared |
| K00376 | Energy metabolism | Nitrogen metabolism | M00529 | Denitrification, nitrate => nitrogen [PATH:map00910 map | nosZ | nitrous-oxide reductase [EC:1.7.2.4] | shared |
| K02567 | Energy metabolism | Nitrogen metabolism | M00529 | Denitrification, nitrate => nitrogen [PATH:map00910 map | napA | periplasmic nitrate reductase NapA [EC:1.7.99.-] | shared |
| K02568 | Energy metabolism | Nitrogen metabolism | M00530 | Dissimilatory nitrate reduction, nitrate => ammonia [PAT | napB | cytochrome c-type protein NapB | shared |
| K02567 | Energy metabolism | Nitrogen metabolism | M00530 | Dissimilatory nitrate reduction, nitrate => ammonia [PAT | napA | periplasmic nitrate reductase NapA [EC:1.7.99.-] | shared |
| K00360 | Energy metabolism | Nitrogen metabolism | M00531 | Assimilatory nitrate reduction, nitrate => ammonia [PATH | nasB | assimilatory nitrate reductase electron transfer subunit | shared |
| K00367 | Energy metabolism | Nitrogen metabolism | M00531 | Assimilatory nitrate reduction, nitrate => ammonia [PATH | narB | ferredoxin-nitrate reductase [EC:1.7.7.2] | shared |
| K08928 | Energy metabolism | Photosynthesis | M00597 | Anoxygenic photosystem II [BR:ko00194] | pufL | photosynthetic reaction center L subunit | shared |
| K08929 | Energy metabolism | Photosynthesis | M00597 | Anoxygenic photosystem II [BR:ko00194] | pufM | photosynthetic reaction center M subunit | shared |
| K00381 | Energy metabolism | Sulfur metabolism | M00176 | Assimilatory sulfate reduction, sulfate => H2S [PATH:ma | cysI | sulfite reductase (NADPH) hemoprotein beta-component | FL-ls |
| K00380 | Energy metabolism | Sulfur metabolism | M00176 | Assimilatory sulfate reduction, sulfate => H2S [PATH:ma | cysJ | sulfite reductase (NADPH) flavoprotein alpha-component | PA-ls |
| K00390 | Energy metabolism | Sulfur metabolism | M00176 | Assimilatory sulfate reduction, sulfate => H2S [PATH:ma | cysH | phosphoadenosine phosphosulfate reductase [EC:1.8.4 | shared |
| K00392 | Energy metabolism | Sulfur metabolism | M00176 | Assimilatory sulfate reduction, sulfate => H2S [PATH:ma | sir | sulfite reductase (ferredoxin) [EC:1.8.7.1] | shared |
| K00394 | Energy metabolism | Sulfur metabolism | M00596 | Dissimilatory sulfate reduction, sulfate => H2S [PATH:m | aprA | adenylylsulfate reductase, subunit A [EC:1.8.99.2] | shared |
| K00395 | Energy metabolism | Sulfur metabolism | M00596 | Dissimilatory sulfate reduction, sulfate => H2S [PATH:m | aprB | adenylylsulfate reductase, subunit B [EC:1.8.99.2] | shared |
| K09810 | Environmental informatio | ABC-2 type and other transport syst | M00255 | Lipoprotein-releasing system [PATH:map02010] [BR:ko0 | lolD | lipoprotein-releasing system ATP-binding protein [EC:3.6 | FL-ls |
| K09808 | Environmental informatio | ABC-2 type and other transport syst | M00255 | Lipoprotein-releasing system [PATH:map02010] [BR:ko0 | lolC_E | lipoprotein-releasing system permease protein | FL-ls |
| K09689 | Environmental informatio | ABC-2 type and other transport syst | M00249 | Capsular polysaccharide transport system [PATH:map02 | ABC-2.CPSE.A | capsular polysaccharide transport system ATP-binding | PA-ls |
| K09688 | Environmental informatio | ABC-2 type and other transport syst | M00249 | Capsular polysaccharide transport system [PATH:map02 | ABC-2.CPSE.P | capsular polysaccharide transport system permease pro | PA-ls |
| K10107 | Environmental informatio | ABC-2 type and other transport syst | M00249 | Capsular polysaccharide transport system [PATH:map02 | ABC-2.CPSE.P1 | capsular polysaccharide transport system permease pro | PA-ls |
| K09691 | Environmental informatio | ABC-2 type and other transport syst | M00250 | Lipopolysaccharide transport system [PATH:map02010] | ABC-2.LPSE.A | lipopolysaccharide transport system ATP-binding protein | PA-ls |
| K09690 | Environmental informatio | ABC-2 type and other transport syst | M00250 | Lipopolysaccharide transport system [PATH:map02010] | ABC-2.LPSE.P | lipopolysaccharide transport system permease protein | PA-ls |
| K09812 | Environmental informatio | ABC-2 type and other transport syst | M00256 | Cell division transport system [PATH:map02010] [BR:ko | ftsE | cell division transport system ATP-binding protein | PA-ls |
| K09811 | Environmental informatio | ABC-2 type and other transport syst | M00256 | Cell division transport system [PATH:map02010] [BR:ko | ftsX | cell division transport system permease protein | PA-ls |
| K09695 | Environmental informatio | ABC-2 type and other transport syst | M00252 | Lipooligosaccharide transport system [PATH:map02010] | nodI | lipooligosaccharide transport system ATP-binding protei | shared |
| K09694 | Environmental informatio | ABC-2 type and other transport syst | M00252 | Lipooligosaccharide transport system [PATH:map02010] | nodJ | lipooligosaccharide transport system permease protein | shared |
| K02193 | Environmental informatio | ABC-2 type and other transport syst | M00259 | Heme transport system [PATH:map02010] [BR:ko02000] | ccmA | heme exporter protein A [EC:3.6.3.41] | shared |
| K02194 | Environmental informatio | ABC-2 type and other transport syst | M00259 | Heme transport system [PATH:map02010] [BR:ko02000] | ccmB | heme exporter protein B | shared |
| K02195 | Environmental informatio | ABC-2 type and other transport syst | M00259 | Heme transport system [PATH:map02010] [BR:ko02000] | ccmC | heme exporter protein C | shared |
| K06861 | Environmental informatio | ABC-2 type and other transport syst | M00320 | Lipopolysaccharide export system [PATH:map02010] [B | lptB | lipopolysaccharide export system ATP-binding protein [E | shared |
| K07091 | Environmental informatio | ABC-2 type and other transport syst | M00320 | Lipopolysaccharide export system [PATH:map02010] [B | lptF | lipopolysaccharide export system permease protein | shared |
| K11720 | Environmental informatio | ABC-2 type and other transport syst | M00320 | Lipopolysaccharide export system [PATH:map02010] [B | lptG | lipopolysaccharide export system permease protein | shared |
| K01493 | Environmental informatio | Bacterial secretion system | M00429 | Competence-related DNA transformation transporter [BR | comEB | dCMP deaminase [EC:3.5.4.12] | FL-ls |
| K02453 | Environmental informatio | Bacterial secretion system | M00331 | Type II general secretion system [PATH:map03070] [BR | gspD | general secretion pathway protein D | PA-ls |
| K02454 | Environmental informatio | Bacterial secretion system | M00331 | Type II general secretion system [PATH:map03070] [BR | gspE | general secretion pathway protein E | PA-ls |
| K02455 | Environmental informatio | Bacterial secretion system | M00331 | Type II general secretion system [PATH:map03070] [BR | gspF | general secretion pathway protein F | PA-ls |
| K02458 | Environmental informatio | Bacterial secretion system | M00331 | Type II general secretion system [PATH:map03070] [BR | gspI | general secretion pathway protein I | PA-ls |
| K02459 | Environmental informatio | Bacterial secretion system | M00331 | Type II general secretion system [PATH:map03070] [BR | gspJ | general secretion pathway protein J | PA-ls |
| K02460 | Environmental informatio | Bacterial secretion system | M00331 | Type II general secretion system [PATH:map03070] [BR | gspK | general secretion pathway protein K | PA-ls |
| K03200 | Environmental informatio | Bacterial secretion system | M00333 | Type IV secretion system [PATH:map03070] [BR:ko020 | virB5, lvhB5 | type IV secretion system protein VirB5 | PA-ls |
| K12536 | Environmental informatio | Bacterial secretion system | M00328 | Hemophore/metalloprotease transport system [BR:ko02 | hasD, prtD, aprD | ATP-binding cassette, subfamily C, bacterial exporter fo | shared |
| K12539 | Environmental informatio | Bacterial secretion system | M00329 | Multiple protein transport system [BR:ko02044] | prsD | ATP-binding cassette, subfamily C, bacterial PrsD | shared |
| K12541 | Environmental informatio | Bacterial secretion system | M00330 | Adhesin protein transport system [BR:ko02044] | lapB | ATP-binding cassette, subfamily C, bacterial LapB | shared |
| K02452 | Environmental informatio | Bacterial secretion system | M00331 | Type II general secretion system [PATH:map03070] [BR | gspC | general secretion pathway protein C | shared |
| K02456 | Environmental informatio | Bacterial secretion system | M00331 | Type II general secretion system [PATH:map03070] [BR | gspG | general secretion pathway protein G | shared |
| K02457 | Environmental informatio | Bacterial secretion system | M00331 | Type II general secretion system [PATH:map03070] [BR | gspH | general secretion pathway protein H | shared |
| K02461 | Environmental informatio | Bacterial secretion system | M00331 | Type II general secretion system [PATH:map03070] [BR | gspL | general secretion pathway protein L | shared |
| K02462 | Environmental informatio | Bacterial secretion system | M00331 | Type II general secretion system [PATH:map03070] [BR | gspM | general secretion pathway protein M | shared |
| K02464 | Environmental informatio | Bacterial secretion system | M00331 | Type II general secretion system [PATH:map03070] [BR | gspO | general secretion pathway protein O [EC:3.4.23.43 2.1.1 | shared |
| K03224 | Environmental informatio | Bacterial secretion system | M00332 | Type III secretion system [PATH:map03070] [BR:ko020 | yscN, sctN, hrcN, s | ATP synthase in type III secretion protein N [EC:3.6.3.5 | shared |

| | | | | | | | |
|---|---|---|---|---|---|---|---|
| K03219 | Environmental informatio | Bacterial secretion system | M00332 | Type III secretion system [PATH:map03070] [BR:ko020 | yscC, sctC, ssaC | type III secretion protein C | shared |
| K03222 | Environmental informatio | Bacterial secretion system | M00332 | Type III secretion system [PATH:map03070] [BR:ko020 | yscJ, sctJ, hrcJ, s | type III secretion protein J | shared |
| K03223 | Environmental informatio | Bacterial secretion system | M00332 | Type III secretion system [PATH:map03070] [BR:ko020 | yscL, sctL | type III secretion protein L | shared |
| K03226 | Environmental informatio | Bacterial secretion system | M00332 | Type III secretion system [PATH:map03070] [BR:ko020 | yscR, sctR, hrcR, s | type III secretion protein R | shared |
| K03228 | Environmental informatio | Bacterial secretion system | M00332 | Type III secretion system [PATH:map03070] [BR:ko020 | yscT, sctT, hrcT, s | type III secretion protein T | shared |
| K03229 | Environmental informatio | Bacterial secretion system | M00332 | Type III secretion system [PATH:map03070] [BR:ko020 | yscU, sctU, hrcU, s | type III secretion protein U | shared |
| K03230 | Environmental informatio | Bacterial secretion system | M00332 | Type III secretion system [PATH:map03070] [BR:ko020 | yscV, sctV, hrcV, s | type III secretion protein V | shared |
| K03195 | Environmental informatio | Bacterial secretion system | M00333 | Type IV secretion system [PATH:map03070] [BR:ko020 | virB10, lvhB10 | type IV secretion system protein VirB10 | shared |
| K03198 | Environmental informatio | Bacterial secretion system | M00333 | Type IV secretion system [PATH:map03070] [BR:ko020 | virB3, lvhB3 | type IV secretion system protein VirB3 | shared |
| K03199 | Environmental informatio | Bacterial secretion system | M00333 | Type IV secretion system [PATH:map03070] [BR:ko020 | virB4, lvhB4 | type IV secretion system protein VirB4 | shared |
| K03201 | Environmental informatio | Bacterial secretion system | M00333 | Type IV secretion system [PATH:map03070] [BR:ko020 | virB6, lvhB6 | type IV secretion system protein VirB6 | shared |
| K03203 | Environmental informatio | Bacterial secretion system | M00333 | Type IV secretion system [PATH:map03070] [BR:ko020 | virB8, lvhB8 | type IV secretion system protein VirB8 | shared |
| K03204 | Environmental informatio | Bacterial secretion system | M00333 | Type IV secretion system [PATH:map03070] [BR:ko020 | virB9, lvhB9 | type IV secretion system protein VirB9 | shared |
| K11892 | Environmental informatio | Bacterial secretion system | M00334 | Type VI secretion system [PATH:map03070] [BR:ko020 | impK, ompA, vasF, | type VI secretion system protein ImpK | shared |
| K11891 | Environmental informatio | Bacterial secretion system | M00334 | Type VI secretion system [PATH:map03070] [BR:ko020 | impL, vasK, icmF | type VI secretion system protein ImpL | shared |
| K11906 | Environmental informatio | Bacterial secretion system | M00334 | Type VI secretion system [PATH:map03070] [BR:ko020 | vasD, lip | type VI secretion system protein VasD | shared |
| K11907 | Environmental informatio | Bacterial secretion system | M00334 | Type VI secretion system [PATH:map03070] [BR:ko020 | vasG, clpV | type VI secretion system protein VasG | shared |
| K11903 | Environmental informatio | Bacterial secretion system | M00334 | Type VI secretion system [PATH:map03070] [BR:ko020 | hcp | type VI secretion system secreted protein Hcp | shared |
| K11904 | Environmental informatio | Bacterial secretion system | M00334 | Type VI secretion system [PATH:map03070] [BR:ko020 | vgrG | type VI secretion system secreted protein VgrG | shared |
| K13408 | Environmental informatio | Bacterial secretion system | M00339 | RaxAB-RaxC type I secretion system [PATH:map04626] | raxA | membrane fusion protein | shared |
| K16299 | Environmental informatio | Bacterial secretion system | M00571 | AlgE-type Mannuronan C-5-Epimerase transport system | eexD | ATP-binding cassette, subfamily C, bacterial EexD | shared |
| K07664 | Environmental informatio | Drug efflux transporter/pump | M00645 | Multidrug resistance, efflux pump SmeABC [PATH:map0 | baeR, smeR | two-component system, OmpR family, response regulat | shared |
| K07642 | Environmental informatio | Drug efflux transporter/pump | M00645 | Multidrug resistance, efflux pump SmeABC [PATH:map0 | baeS, smeS | two-component system, OmpR family, sensor histidine k | shared |
| K07664 | Environmental informatio | Drug efflux transporter/pump | M00646 | Multidrug resistance, efflux pump AcrAD-TolC [PATH:ma | baeR, smeR | two-component system, OmpR family, response regulat | shared |
| K07642 | Environmental informatio | Drug efflux transporter/pump | M00646 | Multidrug resistance, efflux pump AcrAD-TolC [PATH:ma | baeS, smeS | two-component system, OmpR family, sensor histidine k | shared |
| K07799 | Environmental informatio | Drug efflux transporter/pump | M00648 | Multidrug resistance, efflux pump MdtABC [PATH:map02 | mdtA | membrane fusion protein, multidrug efflux system | shared |
| K07788 | Environmental informatio | Drug efflux transporter/pump | M00648 | Multidrug resistance, efflux pump MdtABC [PATH:map02 | mdtB | multidrug efflux pump | shared |
| K07789 | Environmental informatio | Drug efflux transporter/pump | M00648 | Multidrug resistance, efflux pump MdtABC [PATH:map02 | mdtC | multidrug efflux pump | shared |
| K07664 | Environmental informatio | Drug efflux transporter/pump | M00648 | Multidrug resistance, efflux pump MdtABC [PATH:map02 | baeR, smeR | two-component system, OmpR family, response regulat | shared |
| K07642 | Environmental informatio | Drug efflux transporter/pump | M00648 | Multidrug resistance, efflux pump MdtABC [PATH:map02 | baeS, smeS | two-component system, OmpR family, sensor histidine k | shared |
| K05685 | Environmental informatio | Drug efflux transporter/pump | M00709 | Macrolide resistance, MacAB-TolC transporter | macB | macrolide transport system ATP-binding/permease prot | shared |
| K11618 | Environmental informatio | Drug resistance | M00754 | Nisin resistance, phage shock protein homolog LiaH | liaR | two-component system, NarL family, response regulator | FL-ls |
| K08218 | Environmental informatio | Drug resistance | M00628 | beta-Lactam resistance, AmpC system [PATH:map0150 | ampG | MFS transporter, PAT family, beta-lactamase induction | PA-ls |
| K03760 | Environmental informatio | Drug resistance | M00722 | Cationic antimicrobial peptide (CAMP) resistance, phosp | eptA, pmrC | lipid A ethanolaminephosphotransferase [EC:2.7.8.43] | PA-ls |
| K01448 | Environmental informatio | Drug resistance | M00727 | Cationic antimicrobial peptide (CAMP) resistance, N-ace | amiABC | N-acetylmuramoyl-L-alanine amidase [EC:3.5.1.28] | PA-ls |
| K03553 | Environmental informatio | Drug resistance | M00729 | Fluoroquinolone resistance, gyrase-protecting protein Q | recA | recombination protein RecA | PA-ls |
| K07638 | Environmental informatio | Drug resistance | M00742 | Aminoglycoside resistance, protease FtsH | envZ | two-component system, OmpR family, osmolarity senso | PA-ls |
| K07659 | Environmental informatio | Drug resistance | M00742 | Aminoglycoside resistance, protease FtsH | ompR | two-component system, OmpR family, phosphate regulo | PA-ls |
| K07638 | Environmental informatio | Drug resistance | M00743 | Aminoglycoside resistance, protease HtpX | envZ | two-component system, OmpR family, osmolarity senso | PA-ls |
| K07659 | Environmental informatio | Drug resistance | M00743 | Aminoglycoside resistance, protease HtpX | ompR | two-component system, OmpR family, phosphate regulo | PA-ls |
| K03673 | Environmental informatio | Drug resistance | M00728 | Cationic antimicrobial peptide (CAMP) resistance, envel | dsbA | thiol:disulfide interchange protein DsbA | shared |
| K11617 | Environmental informatio | Drug resistance | M00754 | Nisin resistance, phage shock protein homolog LiaH | liaS | two-component system, NarL family, sensor histidine ki | shared |
| K09817 | Environmental informatio | Metallic cation, iron-siderophore and | M00242 | Zinc transport system [PATH:map02010] [BR:ko02000] | znuC | zinc transport system ATP-binding protein [EC:3.6.3.-] | FL-ls |
| K09816 | Environmental informatio | Metallic cation, iron-siderophore and | M00242 | Zinc transport system [PATH:map02010] [BR:ko02000] | znuB | zinc transport system permease protein | FL-ls |
| K02013 | Environmental informatio | Metallic cation, iron-siderophore and | M00240 | Iron complex transport system [PATH:map02010] [BR:k | ABC.FEV.A | iron complex transport system ATP-binding protein [EC: | PA-ls |
| K02015 | Environmental informatio | Metallic cation, iron-siderophore and | M00240 | Iron complex transport system [PATH:map02010] [BR:k | ABC.FEV.P | iron complex transport system permease protein | PA-ls |
| K02016 | Environmental informatio | Metallic cation, iron-siderophore and | M00240 | Iron complex transport system [PATH:map02010] [BR:k | ABC.FEV.S | iron complex transport system substrate-binding protein | PA-ls |
| K02006 | Environmental informatio | Metallic cation, iron-siderophore and | M00245 | Cobalt/nickel transport system [PATH:map02010] [BR:k | cbiO | cobalt/nickel transport system ATP-binding protein | PA-ls |
| K02008 | Environmental informatio | Metallic cation, iron-siderophore and | M00245 | Cobalt/nickel transport system [PATH:map02010] [BR:k | cbiQ | cobalt/nickel transport system permease protein | PA-ls |
| K02007 | Environmental informatio | Metallic cation, iron-siderophore and | M00245 | Cobalt/nickel transport system [PATH:map02010] [BR:k | cbiM | cobalt/nickel transport system permease protein | PA-ls |
| K02006 | Environmental informatio | Metallic cation, iron-siderophore and | M00246 | Nickel transport system [PATH:map02010] [BR:ko02000 | cbiO | cobalt/nickel transport system ATP-binding protein | PA-ls |
| K02008 | Environmental informatio | Metallic cation, iron-siderophore and | M00246 | Nickel transport system [PATH:map02010] [BR:ko02000 | cbiQ | cobalt/nickel transport system permease protein | PA-ls |
| K02007 | Environmental informatio | Metallic cation, iron-siderophore and | M00246 | Nickel transport system [PATH:map02010] [BR:ko02000 | cbiM | cobalt/nickel transport system permease protein | PA-ls |
| K11605 | Environmental informatio | Metallic cation, iron-siderophore and | M00317 | Manganese/iron transport system [PATH:map02010] [B | sitC | manganese/iron transport system permease protein | PA-ls |
| K11606 | Environmental informatio | Metallic cation, iron-siderophore and | M00317 | Manganese/iron transport system [PATH:map02010] [B | sitD | manganese/iron transport system permease protein | PA-ls |
| K03523 | Environmental informatio | Metallic cation, iron-siderophore and | M00581 | Biotin transport system [PATH:map02010] [BR:ko02000] | bioY | biotin transport system substrate-specific component | PA-ls |
| K03523 | Environmental informatio | Metallic cation, iron-siderophore and | M00582 | Energy-coupling factor transport system [PATH:map020 | bioY | biotin transport system substrate-specific component | PA-ls |
| K06074 | Environmental informatio | Metallic cation, iron-siderophore and | M00241 | Vitamin B12 transport system [PATH:map02010] [BR:ko | ABC.VB12.A, btuD | vitamin B12 transport system ATP-binding protein [EC:3 | shared |
| K06073 | Environmental informatio | Metallic cation, iron-siderophore and | M00241 | Vitamin B12 transport system [PATH:map02010] [BR:ko | ABC.VB12.P, btuC | vitamin B12 transport system permease protein | shared |
| K06858 | Environmental informatio | Metallic cation, iron-siderophore and | M00241 | Vitamin B12 transport system [PATH:map02010] [BR:ko | ABC.VB12.S1, btuF | vitamin B12 transport system substrate-binding protein | shared |

| | | | | | | | |
|---|---|---|---|---|---|---|---|
| K09815 | Environmental informatio | Metallic cation, iron-siderophore and | M00242 | Zinc transport system [PATH:map02010] [BR:ko02000] | znuA | zinc transport system substrate-binding protein | shared |
| K10094 | Environmental informatio | Metallic cation, iron-siderophore and | M00246 | Nickel transport system [PATH:map02010] [BR:ko02000 | cbiK | nickel transport protein | shared |
| K11607 | Environmental informatio | Metallic cation, iron-siderophore and | M00317 | Manganese/iron transport system [PATH:map02010] [BI | sitB | manganese/iron transport system ATP-binding protein | shared |
| K11604 | Environmental informatio | Metallic cation, iron-siderophore and | M00317 | Manganese/iron transport system [PATH:map02010] [BI | sitA | manganese/iron transport system substrate-binding prot | shared |
| K11710 | Environmental informatio | Metallic cation, iron-siderophore and | M00319 | Manganese/zinc/iron transport system [PATH:map0201 | troB, mntB, znuC | manganese/zinc/iron transport system ATP- binding prot | shared |
| K11708 | Environmental informatio | Metallic cation, iron-siderophore and | M00319 | Manganese/zinc/iron transport system [PATH:map0201 | troC, mntC, znuB | manganese/zinc/iron transport system permease protei | shared |
| K11709 | Environmental informatio | Metallic cation, iron-siderophore and | M00319 | Manganese/zinc/iron transport system [PATH:map0201 | troD, mntD, znuB | manganese/zinc/iron transport system permease protei | shared |
| K11707 | Environmental informatio | Metallic cation, iron-siderophore and | M00319 | Manganese/zinc/iron transport system [PATH:map0201 | troA, mntA, znuA | manganese/zinc/iron transport system substrate-binding | shared |
| K16785 | Environmental informatio | Metallic cation, iron-siderophore and | M00582 | Energy-coupling factor transport system [PATH:map020 | ecfT | energy-coupling factor transport system permease prote | shared |
| K02017 | Environmental informatio | Mineral and organic ion transport sy | M00189 | Molybdate transport system [PATH:map02010] [BR:ko0 | modC | molybdate transport system ATP-binding protein [EC:3.6 | PA-ls |
| K02018 | Environmental informatio | Mineral and organic ion transport sy | M00189 | Molybdate transport system [PATH:map02010] [BR:ko0 | modB | molybdate transport system permease protein | PA-ls |
| K02020 | Environmental informatio | Mineral and organic ion transport sy | M00189 | Molybdate transport system [PATH:map02010] [BR:ko0 | modA | molybdate transport system substrate-binding protein | PA-ls |
| K02012 | Environmental informatio | Mineral and organic ion transport sy | M00190 | Iron(III) transport system [PATH:map02010] [BR:ko020 | afuA, fbpA | iron(III) transport system substrate-binding protein | PA-ls |
| K02052 | Environmental informatio | Mineral and organic ion transport sy | M00193 | Putative spermidine/putrescine transport system [BR:k | ABC.SP.A | putative spermidine/putrescine transport system ATP-bi | PA-ls |
| K02053 | Environmental informatio | Mineral and organic ion transport sy | M00193 | Putative spermidine/putrescine transport system [BR:k | ABC.SP.P | putative spermidine/putrescine transport system permea | PA-ls |
| K02054 | Environmental informatio | Mineral and organic ion transport sy | M00193 | Putative spermidine/putrescine transport system [BR:k | ABC.SP.P1 | putative spermidine/putrescine transport system permea | PA-ls |
| K02055 | Environmental informatio | Mineral and organic ion transport sy | M00193 | Putative spermidine/putrescine transport system [BR:k | ABC.SP.S | putative spermidine/putrescine transport system substra | PA-ls |
| K02000 | Environmental informatio | Mineral and organic ion transport sy | M00208 | Glycine betaine/proline transport system [PATH:map020 | proV | glycine betaine/proline transport system ATP-binding pr | PA-ls |
| K02001 | Environmental informatio | Mineral and organic ion transport sy | M00208 | Glycine betaine/proline transport system [PATH:map020 | proW | glycine betaine/proline transport system permease prote | PA-ls |
| K05847 | Environmental informatio | Mineral and organic ion transport sy | M00209 | Osmoprotectant transport system [PATH:map02010] [BI | opuA | osmoprotectant transport system ATP-binding protein | PA-ls |
| K05846 | Environmental informatio | Mineral and organic ion transport sy | M00209 | Osmoprotectant transport system [PATH:map02010] [BI | opuBD | osmoprotectant transport system permease protein | PA-ls |
| K05845 | Environmental informatio | Mineral and organic ion transport sy | M00209 | Osmoprotectant transport system [PATH:map02010] [BI | opuC | osmoprotectant transport system substrate-binding prot | PA-ls |
| K06857 | Environmental informatio | Mineral and organic ion transport sy | M00186 | Tungstate transport system [PATH:map02010] [BR:ko02 | tupC, vupC | tungstate transport system ATP-binding protein [EC:7.3 | shared |
| K05773 | Environmental informatio | Mineral and organic ion transport sy | M00186 | Tungstate transport system [PATH:map02010] [BR:ko02 | tupB, vupB | tungstate transport system permease protein | shared |
| K05772 | Environmental informatio | Mineral and organic ion transport sy | M00186 | Tungstate transport system [PATH:map02010] [BR:ko02 | tupA, vupA | tungstate transport system substrate-binding protein | shared |
| K05776 | Environmental informatio | Mineral and organic ion transport sy | M00189 | Molybdate transport system [PATH:map02010] [BR:ko0 | modF | molybdate transport system ATP-binding protein | shared |
| K02010 | Environmental informatio | Mineral and organic ion transport sy | M00190 | Iron(III) transport system [PATH:map02010] [BR:ko020 | afuC, fbpC | iron(III) transport system ATP-binding protein [EC:3.6.3. | shared |
| K02011 | Environmental informatio | Mineral and organic ion transport sy | M00190 | Iron(III) transport system [PATH:map02010] [BR:ko020 | afuB, fbpB | iron(III) transport system permease protein | shared |
| K02062 | Environmental informatio | Mineral and organic ion transport sy | M00191 | Thiamine transport system [PATH:map02010] [BR:ko02 | thiQ | thiamine transport system ATP-binding protein | shared |
| K02063 | Environmental informatio | Mineral and organic ion transport sy | M00191 | Thiamine transport system [PATH:map02010] [BR:ko02 | thiP | thiamine transport system permease protein | shared |
| K02064 | Environmental informatio | Mineral and organic ion transport sy | M00191 | Thiamine transport system [PATH:map02010] [BR:ko02 | thiB, tbpA | thiamine transport system substrate-binding protein | shared |
| K02002 | Environmental informatio | Mineral and organic ion transport sy | M00208 | Glycine betaine/proline transport system [PATH:map020 | proX | glycine betaine/proline transport system substrate-bindi | shared |
| K11072 | Environmental informatio | Mineral and organic ion transport sy | M00299 | Spermidine/putrescine transport system [PATH:map0201 | potA | spermidine/putrescine transport system ATP-binding pro | shared |
| K11070 | Environmental informatio | Mineral and organic ion transport sy | M00299 | Spermidine/putrescine transport system [PATH:map0201 | potC | spermidine/putrescine transport system permease prote | shared |
| K11071 | Environmental informatio | Mineral and organic ion transport sy | M00299 | Spermidine/putrescine transport system [PATH:map0201 | potB | spermidine/putrescine transport system permease prote | shared |
| K11069 | Environmental informatio | Mineral and organic ion transport sy | M00299 | Spermidine/putrescine transport system [PATH:map0201 | potD | spermidine/putrescine transport system substrate-bindir | shared |
| K11076 | Environmental informatio | Mineral and organic ion transport sy | M00300 | Putrescine transport system [PATH:map02010] [BR:ko0 | potG | putrescine transport system ATP-binding protein | shared |
| K11074 | Environmental informatio | Mineral and organic ion transport sy | M00300 | Putrescine transport system [PATH:map02010] [BR:ko0 | potI | putrescine transport system permease protein | shared |
| K11075 | Environmental informatio | Mineral and organic ion transport sy | M00300 | Putrescine transport system [PATH:map02010] [BR:ko0 | potH | putrescine transport system permease protein | shared |
| K11073 | Environmental informatio | Mineral and organic ion transport sy | M00300 | Putrescine transport system [PATH:map02010] [BR:ko0 | potF | putrescine transport system substrate-binding protein | shared |
| K11079 | Environmental informatio | Mineral and organic ion transport sy | M00301 | Mannopine transport system [PATH:map02010] [BR:ko0 | attA2 | mannopine transport system permease protein | shared |
| K11084 | Environmental informatio | Mineral and organic ion transport sy | M00302 | 2-Aminoethylphosphonate transport system [PATH:map | phnT | 2-aminoethylphosphonate transport system ATP-binding | shared |
| K11083 | Environmental informatio | Mineral and organic ion transport sy | M00302 | 2-Aminoethylphosphonate transport system [PATH:map | phnU | 2-aminoethylphosphonate transport system permease p | shared |
| K11952 | Environmental informatio | Mineral and organic ion transport sy | M00321 | Bicarbonate transport system [PATH:map02010] [BR:ko | cmpC | bicarbonate transport system ATP-binding protein [EC:3 | shared |
| K15599 | Environmental informatio | Mineral and organic ion transport sy | M00442 | Putative hydroxymethylpyrimidine transport system [PA | thiX | putative hydroxymethylpyrimidine transport system pern | shared |
| K02031 | Environmental informatio | Peptide and nickel transport system | M00239 | Peptides/nickel transport system [PATH:map02010] [BR | ABC.PE.A | peptide/nickel transport system ATP-binding protein | FL-ls |
| K02032 | Environmental informatio | Peptide and nickel transport system | M00239 | Peptides/nickel transport system [PATH:map02010] [BR | ABC.PE.A1 | peptide/nickel transport system ATP-binding protein | shared |
| K02033 | Environmental informatio | Peptide and nickel transport system | M00239 | Peptides/nickel transport system [PATH:map02010] [BR | ABC.PE.P | peptide/nickel transport system permease protein | shared |
| K02034 | Environmental informatio | Peptide and nickel transport system | M00239 | Peptides/nickel transport system [PATH:map02010] [BR | ABC.PE.P1 | peptide/nickel transport system permease protein | shared |
| K02035 | Environmental informatio | Peptide and nickel transport system | M00239 | Peptides/nickel transport system [PATH:map02010] [BR | ABC.PE.S | peptide/nickel transport system substrate-binding protei | shared |
| K12371 | Environmental informatio | Peptide and nickel transport system | M00324 | Dipeptide transport system [PATH:map02010] [BR:ko02 | dppD | dipeptide transport system ATP-binding protein | shared |
| K12372 | Environmental informatio | Peptide and nickel transport system | M00324 | Dipeptide transport system [PATH:map02010] [BR:ko02 | dppF | dipeptide transport system ATP-binding protein | shared |
| K12369 | Environmental informatio | Peptide and nickel transport system | M00324 | Dipeptide transport system [PATH:map02010] [BR:ko02 | dppB | dipeptide transport system permease protein | shared |
| K12370 | Environmental informatio | Peptide and nickel transport system | M00324 | Dipeptide transport system [PATH:map02010] [BR:ko02 | dppC | dipeptide transport system permease protein | shared |
| K13892 | Environmental informatio | Peptide and nickel transport system | M00348 | Glutathione transport system [PATH:map02010] [BR:ko | gsiA | glutathione transport system ATP-binding protein | shared |
| K13890 | Environmental informatio | Peptide and nickel transport system | M00348 | Glutathione transport system [PATH:map02010] [BR:ko | gsiC | glutathione transport system permease protein | shared |
| K13891 | Environmental informatio | Peptide and nickel transport system | M00348 | Glutathione transport system [PATH:map02010] [BR:ko | gsiD | glutathione transport system permease protein | shared |
| K13896 | Environmental informatio | Peptide and nickel transport system | M00349 | Microcin C transport system [PATH:map02010] [BR:ko0 | yejF | microcin C transport system ATP-binding protein | shared |
| K13894 | Environmental informatio | Peptide and nickel transport system | M00349 | Microcin C transport system [PATH:map02010] [BR:ko0 | yejB | microcin C transport system permease protein | shared |
| K13895 | Environmental informatio | Peptide and nickel transport system | M00349 | Microcin C transport system [PATH:map02010] [BR:ko0 | yejE | microcin C transport system permease protein | shared |

| | | | | | | | |
|---|---|---|---|---|---|---|---|
| K13893 | Environmental informatio | Peptide and nickel transport system | M00349 | Microcin C transport system [PATH:map02010] [BR:ko0 | yejA | microcin C transport system substrate-binding protein | shared |
| K02041 | Environmental informatio | Phosphate and amino acid transport | M00223 | Phosphonate transport system [PATH:map02010] [BR:k | phnC | phosphonate transport system ATP-binding protein [EC: | FL-ls |
| K02042 | Environmental informatio | Phosphate and amino acid transport | M00223 | Phosphonate transport system [PATH:map02010] [BR:k | phnE | phosphonate transport system permease protein | FL-ls |
| K02044 | Environmental informatio | Phosphate and amino acid transport | M00223 | Phosphonate transport system [PATH:map02010] [BR:k | phnD | phosphonate transport system substrate-binding protein | FL-ls |
| K10024 | Environmental informatio | Phosphate and amino acid transport | M00235 | Arginine/ornithine transport system [PATH:map02010] [E | aotQ | arginine/ornithine transport system permease protein | FL-ls |
| K02036 | Environmental informatio | Phosphate and amino acid transport | M00222 | Phosphate transport system [PATH:map02010] [BR:ko0 | pstB | phosphate transport system ATP-binding protein [EC:7.3 | PA-ls |
| K02037 | Environmental informatio | Phosphate and amino acid transport | M00222 | Phosphate transport system [PATH:map02010] [BR:ko0 | pstC | phosphate transport system permease protein | PA-ls |
| K02038 | Environmental informatio | Phosphate and amino acid transport | M00222 | Phosphate transport system [PATH:map02010] [BR:ko0 | pstA | phosphate transport system permease protein | PA-ls |
| K10036 | Environmental informatio | Phosphate and amino acid transport | M00227 | Glutamine transport system [PATH:map02010] [BR:ko02 | glnH | glutamine transport system substrate-binding protein | PA-ls |
| K10019 | Environmental informatio | Phosphate and amino acid transport | M00231 | Octopine/nopaline transport system [PATH:map02010] [ | occM, nocM | octopine/nopaline transport system permease protein | PA-ls |
| K10020 | Environmental informatio | Phosphate and amino acid transport | M00231 | Octopine/nopaline transport system [PATH:map02010] [ | occQ, nocQ | octopine/nopaline transport system permease protein | PA-ls |
| K10018 | Environmental informatio | Phosphate and amino acid transport | M00231 | Octopine/nopaline transport system [PATH:map02010] [ | occT, nocT | octopine/nopaline transport system substrate-binding pr | PA-ls |
| K09970 | Environmental informatio | Phosphate and amino acid transport | M00232 | General L-amino acid transport system [PATH:map0201 | aapQ, bztB | general L-amino acid transport system permease protein | PA-ls |
| K09971 | Environmental informatio | Phosphate and amino acid transport | M00232 | General L-amino acid transport system [PATH:map0201 | aapM, bztC | general L-amino acid transport system permease protein | PA-ls |
| K09969 | Environmental informatio | Phosphate and amino acid transport | M00232 | General L-amino acid transport system [PATH:map0201 | aapJ, bztA | general L-amino acid transport system substrate-binding | PA-ls |
| K10008 | Environmental informatio | Phosphate and amino acid transport | M00233 | Glutamate transport system [PATH:map02010] [BR:ko02 | gluA | glutamate transport system ATP-binding protein [EC:7.4 | PA-ls |
| K10006 | Environmental informatio | Phosphate and amino acid transport | M00233 | Glutamate transport system [PATH:map02010] [BR:ko02 | gluC | glutamate transport system permease protein | PA-ls |
| K10007 | Environmental informatio | Phosphate and amino acid transport | M00233 | Glutamate transport system [PATH:map02010] [BR:ko02 | gluD | glutamate transport system permease protein | PA-ls |
| K10005 | Environmental informatio | Phosphate and amino acid transport | M00233 | Glutamate transport system [PATH:map02010] [BR:ko02 | gluB | glutamate transport system substrate-binding protein | PA-ls |
| K02071 | Environmental informatio | Phosphate and amino acid transport | M00238 | D-Methionine transport system [PATH:map02010] [BR:k | metN | D-methionine transport system ATP-binding protein | PA-ls |
| K02072 | Environmental informatio | Phosphate and amino acid transport | M00238 | D-Methionine transport system [PATH:map02010] [BR:k | metI | D-methionine transport system permease protein | PA-ls |
| K02073 | Environmental informatio | Phosphate and amino acid transport | M00238 | D-Methionine transport system [PATH:map02010] [BR:k | metQ | D-methionine transport system substrate-binding protein | PA-ls |
| K10017 | Environmental informatio | Phosphate and amino acid transport | M00225 | Lysine/arginine/ornithine transport system [PATH:map02 | hisP | histidine transport system ATP-binding protein [EC:7.4.2 | shared |
| K10015 | Environmental informatio | Phosphate and amino acid transport | M00225 | Lysine/arginine/ornithine transport system [PATH:map02 | hisM | histidine transport system permease protein | shared |
| K10016 | Environmental informatio | Phosphate and amino acid transport | M00225 | Lysine/arginine/ornithine transport system [PATH:map02 | hisQ | histidine transport system permease protein | shared |
| K10013 | Environmental informatio | Phosphate and amino acid transport | M00225 | Lysine/arginine/ornithine transport system [PATH:map02 | argT | lysine/arginine/ornithine transport system substrate-bind | shared |
| K10017 | Environmental informatio | Phosphate and amino acid transport | M00226 | Histidine transport system [PATH:map02010] [BR:ko020 | hisP | histidine transport system ATP-binding protein [EC:7.4.2 | shared |
| K10015 | Environmental informatio | Phosphate and amino acid transport | M00226 | Histidine transport system [PATH:map02010] [BR:ko020 | hisM | histidine transport system permease protein | shared |
| K10016 | Environmental informatio | Phosphate and amino acid transport | M00226 | Histidine transport system [PATH:map02010] [BR:ko020 | hisQ | histidine transport system permease protein | shared |
| K10038 | Environmental informatio | Phosphate and amino acid transport | M00227 | Glutamine transport system [PATH:map02010] [BR:ko02 | glnQ | glutamine transport system ATP-binding protein [EC:7.4 | shared |
| K10037 | Environmental informatio | Phosphate and amino acid transport | M00227 | Glutamine transport system [PATH:map02010] [BR:ko02 | glnP | glutamine transport system permease protein | shared |
| K10041 | Environmental informatio | Phosphate and amino acid transport | M00228 | Aspartate/glutamate/glutamine transport system [PATH: | peb1C, glnQ | aspartate/glutamate/glutamine transport system ATP-bin | shared |
| K10040 | Environmental informatio | Phosphate and amino acid transport | M00228 | Aspartate/glutamate/glutamine transport system [PATH: | peb1B, glnP, glnM | aspartate/glutamate/glutamine transport system permea | shared |
| K10039 | Environmental informatio | Phosphate and amino acid transport | M00228 | Aspartate/glutamate/glutamine transport system [PATH: | peb1A, glnH | aspartate/glutamate/glutamine transport system substra | shared |
| K10021 | Environmental informatio | Phosphate and amino acid transport | M00231 | Octopine/nopaline transport system [PATH:map02010] [ | occP, nocP | octopine/nopaline transport system ATP-binding protein | shared |
| K09972 | Environmental informatio | Phosphate and amino acid transport | M00232 | General L-amino acid transport system [PATH:map0201 | aapP, bztD | general L-amino acid transport system ATP-binding prot | shared |
| K10010 | Environmental informatio | Phosphate and amino acid transport | M00234 | Cystine transport system [PATH:map02010] [BR:ko0200 | tcyC, yecC | L-cystine transport system ATP-binding protein [EC:7.4. | shared |
| K10009 | Environmental informatio | Phosphate and amino acid transport | M00234 | Cystine transport system [PATH:map02010] [BR:ko0200 | tcyB, yecS | L-cystine transport system permease protein | shared |
| K10025 | Environmental informatio | Phosphate and amino acid transport | M00235 | Arginine/ornithine transport system [PATH:map02010] [E | aotP | arginine/ornithine transport system ATP-binding protein [ | shared |
| K10023 | Environmental informatio | Phosphate and amino acid transport | M00235 | Arginine/ornithine transport system [PATH:map02010] [E | aotM | arginine/ornithine transport system permease protein | shared |
| K10022 | Environmental informatio | Phosphate and amino acid transport | M00235 | Arginine/ornithine transport system [PATH:map02010] [E | aotJ | arginine/ornithine transport system substrate-binding pro | shared |
| K11957 | Environmental informatio | Phosphate and amino acid transport | M00322 | Neutral amino acid transport system [PATH:map02010] [ | natA | neutral amino acid transport system ATP-binding protein | shared |
| K11962 | Environmental informatio | Phosphate and amino acid transport | M00323 | Urea transport system [PATH:map02010] [BR:ko02000] | urtD | urea transport system ATP-binding protein | shared |
| K11963 | Environmental informatio | Phosphate and amino acid transport | M00323 | Urea transport system [PATH:map02010] [BR:ko02000] | urtE | urea transport system ATP-binding protein | shared |
| K11960 | Environmental informatio | Phosphate and amino acid transport | M00323 | Urea transport system [PATH:map02010] [BR:ko02000] | urtB | urea transport system permease protein | shared |
| K11961 | Environmental informatio | Phosphate and amino acid transport | M00323 | Urea transport system [PATH:map02010] [BR:ko02000] | urtC | urea transport system permease protein | shared |
| K11959 | Environmental informatio | Phosphate and amino acid transport | M00323 | Urea transport system [PATH:map02010] [BR:ko02000] | urtA | urea transport system substrate-binding protein | shared |
| K02755 | Environmental informatio | Phosphotransferase system (PTS) | M00271 | PTS system, beta-glucosides-specific II component [PA | PTS-Bgl-EIIA, bglF, | PTS system, beta-glucoside-specific IIA component [EC | shared |
| K02757 | Environmental informatio | Phosphotransferase system (PTS) | M00271 | PTS system, beta-glucosides-specific II component [PA | PTS-Bgl-EIIC, bglF, | PTS system, beta-glucoside-specific IIC component | shared |
| K02761 | Environmental informatio | Phosphotransferase system (PTS) | M00275 | PTS system, cellobiose-specific II component [PATH:ma | PTS-Cel-EIIC, celB, | PTS system, cellobiose-specific IIC component | shared |
| K10109 | Environmental informatio | Saccharide, polyol, and lipid transpo | M00194 | Maltose/maltodextrin transport system [PATH:map0201 | malF | maltose/maltodextrin transport system permease protein | PA-ls |
| K10110 | Environmental informatio | Saccharide, polyol, and lipid transpo | M00194 | Maltose/maltodextrin transport system [PATH:map0201 | malG | maltose/maltodextrin transport system permease protein | PA-ls |
| K05816 | Environmental informatio | Saccharide, polyol, and lipid transpo | M00198 | Putative sn-glycerol-phosphate transport system [PATH | ugpC | sn-glycerol 3-phosphate transport system ATP-binding p | PA-ls |
| K05814 | Environmental informatio | Saccharide, polyol, and lipid transpo | M00198 | Putative sn-glycerol-phosphate transport system [PATH | ugpA | sn-glycerol 3-phosphate transport system permease pro | PA-ls |
| K05813 | Environmental informatio | Saccharide, polyol, and lipid transpo | M00198 | Putative sn-glycerol-phosphate transport system [PATH | ugpB | sn-glycerol 3-phosphate transport system substrate-bin | PA-ls |
| K10228 | Environmental informatio | Saccharide, polyol, and lipid transpo | M00200 | Putative sorbitol/mannitol transport system [PATH:map0 | smoF, mtlF | sorbitol/mannitol transport system permease protein | PA-ls |
| K10229 | Environmental informatio | Saccharide, polyol, and lipid transpo | M00200 | Putative sorbitol/mannitol transport system [PATH:map0 | smoG, mtlG | sorbitol/mannitol transport system permease protein | PA-ls |
| K10227 | Environmental informatio | Saccharide, polyol, and lipid transpo | M00200 | Putative sorbitol/mannitol transport system [PATH:map0 | smoE, mtlE | sorbitol/mannitol transport system substrate-binding pro | PA-ls |
| K10237 | Environmental informatio | Saccharide, polyol, and lipid transpo | M00204 | Trehalose/maltose transport system [PATH:map02010] [ | thuF, sugA | trehalose/maltose transport system permease protein | PA-ls |
| K10238 | Environmental informatio | Saccharide, polyol, and lipid transpo | M00204 | Trehalose/maltose transport system [PATH:map02010] [ | thuG, sugB | trehalose/maltose transport system permease protein | PA-ls |

| | | | | | | | |
|---|---|---|---|---|---|---|---|
| K10236 | Environmental informatio | Saccharide, polyol, and lipid transpo | M00204 | Trehalose/maltose transport system [PATH:map02010] [ | thuE | trehalose/maltose transport system substrate-binding p | PA-ls |
| K02065 | Environmental informatio | Saccharide, polyol, and lipid transpo | M00210 | Phospholipid transport system [PATH:map02010] [BR:k | mlaF, linL, mkl | phospholipid/cholesterol/gamma-HCH transport system | PA-ls |
| K02066 | Environmental informatio | Saccharide, polyol, and lipid transpo | M00210 | Phospholipid transport system [PATH:map02010] [BR:k | mlaE, linK | phospholipid/cholesterol/gamma-HCH transport system | PA-ls |
| K02067 | Environmental informatio | Saccharide, polyol, and lipid transpo | M00210 | Phospholipid transport system [PATH:map02010] [BR:k | mlaD, linM | phospholipid/cholesterol/gamma-HCH transport system | PA-ls |
| K10441 | Environmental informatio | Saccharide, polyol, and lipid transpo | M00212 | Ribose transport system [PATH:map02010] [BR:ko0200 | rbsA | ribose transport system ATP-binding protein [EC:3.6.3.1 | PA-ls |
| K10440 | Environmental informatio | Saccharide, polyol, and lipid transpo | M00212 | Ribose transport system [PATH:map02010] [BR:ko0200 | rbsC | ribose transport system permease protein | PA-ls |
| K10547 | Environmental informatio | Saccharide, polyol, and lipid transpo | M00216 | Multiple sugar transport system [PATH:map02010] [BR: | ABC.GGU.P, gguB | putative multiple sugar transport system permease prot | PA-ls |
| K10546 | Environmental informatio | Saccharide, polyol, and lipid transpo | M00216 | Multiple sugar transport system [PATH:map02010] [BR: | ABC.GGU.S, chvE | putative multiple sugar transport system substrate-bindi | PA-ls |
| K10554 | Environmental informatio | Saccharide, polyol, and lipid transpo | M00218 | Fructose transport system [PATH:map02010] [BR:ko02 | frcA | fructose transport system ATP-binding protein | PA-ls |
| K10553 | Environmental informatio | Saccharide, polyol, and lipid transpo | M00218 | Fructose transport system [PATH:map02010] [BR:ko02 | frcC | fructose transport system permease protein | PA-ls |
| K10552 | Environmental informatio | Saccharide, polyol, and lipid transpo | M00218 | Fructose transport system [PATH:map02010] [BR:ko02 | frcB | fructose transport system substrate-binding protein | PA-ls |
| K02065 | Environmental informatio | Saccharide, polyol, and lipid transpo | M00669 | gamma-Hexachlorocyclohexane transport system [BR:k | mlaF, linL, mkl | phospholipid/cholesterol/gamma-HCH transport system | PA-ls |
| K02066 | Environmental informatio | Saccharide, polyol, and lipid transpo | M00669 | gamma-Hexachlorocyclohexane transport system [BR:k | mlaE, linK | phospholipid/cholesterol/gamma-HCH transport system | PA-ls |
| K02067 | Environmental informatio | Saccharide, polyol, and lipid transpo | M00669 | gamma-Hexachlorocyclohexane transport system [BR:k | mlaD, linM | phospholipid/cholesterol/gamma-HCH transport system | PA-ls |
| K02065 | Environmental informatio | Saccharide, polyol, and lipid transpo | M00670 | Mce transport system [BR:ko02000] | mlaF, linL, mkl | phospholipid/cholesterol/gamma-HCH transport system | PA-ls |
| K02066 | Environmental informatio | Saccharide, polyol, and lipid transpo | M00670 | Mce transport system [BR:ko02000] | mlaE, linK | phospholipid/cholesterol/gamma-HCH transport system | PA-ls |
| K02067 | Environmental informatio | Saccharide, polyol, and lipid transpo | M00670 | Mce transport system [BR:ko02000] | mlaD, linM | phospholipid/cholesterol/gamma-HCH transport system | PA-ls |
| K10112 | Environmental informatio | Saccharide, polyol, and lipid transpo | M00194 | Maltose/maltodextrin transport system [PATH:map0201 | msmX, msmK, malK | multiple sugar transport system ATP-binding protein | shared |
| K10111 | Environmental informatio | Saccharide, polyol, and lipid transpo | M00194 | Maltose/maltodextrin transport system [PATH:map0201 | malK, mtlK, thuK | multiple sugar transport system ATP-binding protein [E | shared |
| K10112 | Environmental informatio | Saccharide, polyol, and lipid transpo | M00196 | Multiple sugar transport system [PATH:map02010] [BR: | msmX, msmK, malK | multiple sugar transport system ATP-binding protein | shared |
| K10118 | Environmental informatio | Saccharide, polyol, and lipid transpo | M00196 | Multiple sugar transport system [PATH:map02010] [BR: | msmF | raffinose/stachyose/melibiose transport system permea | shared |
| K10119 | Environmental informatio | Saccharide, polyol, and lipid transpo | M00196 | Multiple sugar transport system [PATH:map02010] [BR: | msmG | raffinose/stachyose/melibiose transport system permea | shared |
| K10117 | Environmental informatio | Saccharide, polyol, and lipid transpo | M00196 | Multiple sugar transport system [PATH:map02010] [BR: | msmE | raffinose/stachyose/melibiose transport system substra | shared |
| K10112 | Environmental informatio | Saccharide, polyol, and lipid transpo | M00197 | Putative fructooligosaccharide transport system [BR:ko | msmX, msmK, malK | multiple sugar transport system ATP-binding protein | shared |
| K05815 | Environmental informatio | Saccharide, polyol, and lipid transpo | M00198 | Putative sn-glycerol-phosphate transport system [PATH | ugpE | sn-glycerol 3-phosphate transport system permease pro | shared |
| K10189 | Environmental informatio | Saccharide, polyol, and lipid transpo | M00199 | L-Arabinose/lactose transport system [PATH:map02010] | lacF, araP | lactose/L-arabinose transport system permease protein | shared |
| K10190 | Environmental informatio | Saccharide, polyol, and lipid transpo | M00199 | L-Arabinose/lactose transport system [PATH:map02010] | lacG, araQ | lactose/L-arabinose transport system permease protein | shared |
| K10112 | Environmental informatio | Saccharide, polyol, and lipid transpo | M00200 | Putative sorbitol/mannitol transport system [PATH:map0 | msmX, msmK, malK | multiple sugar transport system ATP-binding protein | shared |
| K10111 | Environmental informatio | Saccharide, polyol, and lipid transpo | M00200 | Putative sorbitol/mannitol transport system [PATH:map0 | malK, mtlK, thuK | multiple sugar transport system ATP-binding protein [E | shared |
| K10235 | Environmental informatio | Saccharide, polyol, and lipid transpo | M00201 | alpha-Glucoside transport system [PATH:map02010] [B | aglK | alpha-glucoside transport system ATP-binding protein | shared |
| K10233 | Environmental informatio | Saccharide, polyol, and lipid transpo | M00201 | alpha-Glucoside transport system [PATH:map02010] [B | aglF, ggtC | alpha-glucoside transport system permease protein | shared |
| K10234 | Environmental informatio | Saccharide, polyol, and lipid transpo | M00201 | alpha-Glucoside transport system [PATH:map02010] [B | aglG, ggtD | alpha-glucoside transport system permease protein | shared |
| K10232 | Environmental informatio | Saccharide, polyol, and lipid transpo | M00201 | alpha-Glucoside transport system [PATH:map02010] [B | aglE, ggtB | alpha-glucoside transport system substrate-binding prot | shared |
| K10112 | Environmental informatio | Saccharide, polyol, and lipid transpo | M00201 | alpha-Glucoside transport system [PATH:map02010] [B | msmX, msmK, malK | multiple sugar transport system ATP-binding protein | shared |
| K10195 | Environmental informatio | Saccharide, polyol, and lipid transpo | M00202 | Oligogalacturonide transport system [PATH:map02010] | togA | oligogalacturonide transport system ATP-binding protein | shared |
| K10192 | Environmental informatio | Saccharide, polyol, and lipid transpo | M00202 | Oligogalacturonide transport system [PATH:map02010] | togB | oligogalacturonide transport system substrate-binding p | shared |
| K10199 | Environmental informatio | Saccharide, polyol, and lipid transpo | M00203 | Glucose/arabinose transport system [PATH:map02010] | ABC.GLC.A | glucose/arabinose transport system ATP-binding protein | shared |
| K10111 | Environmental informatio | Saccharide, polyol, and lipid transpo | M00204 | Trehalose/maltose transport system [PATH:map02010] [ | malK, mtlK, thuK | multiple sugar transport system ATP-binding protein [E | shared |
| K10112 | Environmental informatio | Saccharide, polyol, and lipid transpo | M00206 | Cellobiose transport system [PATH:map02010] [BR:ko0 | msmX, msmK, malK | multiple sugar transport system ATP-binding protein | shared |
| K10112 | Environmental informatio | Saccharide, polyol, and lipid transpo | M00207 | Putative multiple sugar transport system [BR:ko02000] | msmX, msmK, malK | multiple sugar transport system ATP-binding protein | shared |
| K10111 | Environmental informatio | Saccharide, polyol, and lipid transpo | M00207 | Putative multiple sugar transport system [BR:ko02000] | malK, mtlK, thuK | multiple sugar transport system ATP-binding protein [E | shared |
| K07323 | Environmental informatio | Saccharide, polyol, and lipid transpo | M00210 | Phospholipid transport system [PATH:map02010] [BR:k | mlaC | phospholipid transport system substrate-binding protein | shared |
| K07122 | Environmental informatio | Saccharide, polyol, and lipid transpo | M00210 | Phospholipid transport system [PATH:map02010] [BR:k | mlaB | phospholipid transport system transporter-binding protei | shared |
| K10539 | Environmental informatio | Saccharide, polyol, and lipid transpo | M00213 | L-Arabinose transport system [PATH:map02010] [BR:ko | araG | L-arabinose transport system ATP-binding protein [EC:3 | shared |
| K10545 | Environmental informatio | Saccharide, polyol, and lipid transpo | M00215 | D-Xylose transport system [PATH:map02010] [BR:ko02 | xylG | D-xylose transport system ATP-binding protein [EC:3.6. | shared |
| K10544 | Environmental informatio | Saccharide, polyol, and lipid transpo | M00215 | D-Xylose transport system [PATH:map02010] [BR:ko02 | xylH | D-xylose transport system permease protein | shared |
| K10543 | Environmental informatio | Saccharide, polyol, and lipid transpo | M00215 | D-Xylose transport system [PATH:map02010] [BR:ko02 | xylF | D-xylose transport system substrate-binding protein | shared |
| K10548 | Environmental informatio | Saccharide, polyol, and lipid transpo | M00216 | Multiple sugar transport system [PATH:map02010] [BR: | ABC.GGU.A, gguA | putative multiple sugar transport system ATP-binding pr | shared |
| K10549 | Environmental informatio | Saccharide, polyol, and lipid transpo | M00217 | D-Allose transport system [PATH:map02010] [BR:ko020 | alsB | D-allose transport system substrate-binding protein | shared |
| K10556 | Environmental informatio | Saccharide, polyol, and lipid transpo | M00219 | AI-2 transport system [PATH:map02010] [BR:ko02000] | lsrC | AI-2 transport system permease protein | shared |
| K10562 | Environmental informatio | Saccharide, polyol, and lipid transpo | M00220 | Rhamnose transport system [PATH:map02010] [BR:ko0 | rhaT | rhamnose transport system ATP-binding protein [EC:3.6 | shared |
| K10560 | Environmental informatio | Saccharide, polyol, and lipid transpo | M00220 | Rhamnose transport system [PATH:map02010] [BR:ko0 | rhaP | rhamnose transport system permease protein | shared |
| K10561 | Environmental informatio | Saccharide, polyol, and lipid transpo | M00220 | Rhamnose transport system [PATH:map02010] [BR:ko0 | rhaQ | rhamnose transport system permease protein | shared |
| K10559 | Environmental informatio | Saccharide, polyol, and lipid transpo | M00220 | Rhamnose transport system [PATH:map02010] [BR:ko0 | rhaS | rhamnose transport system substrate-binding protein | shared |
| K10112 | Environmental informatio | Saccharide, polyol, and lipid transpo | M00491 | arabinogalactan oligomer/maltooligosaccharide transport | msmX, msmK, malK | multiple sugar transport system ATP-binding protein | shared |
| K10111 | Environmental informatio | Saccharide, polyol, and lipid transpo | M00491 | arabinogalactan oligomer/maltooligosaccharide transport | malK, mtlK, thuK | multiple sugar transport system ATP-binding protein [E | shared |
| K10112 | Environmental informatio | Saccharide, polyol, and lipid transpo | M00602 | Arabinosaccharide transport system [PATH:map02010] | msmX, msmK, malK | multiple sugar transport system ATP-binding protein | shared |
| K10112 | Environmental informatio | Saccharide, polyol, and lipid transpo | M00605 | Glucose/mannose transport system [PATH:map02010] [ | msmX, msmK, malK | multiple sugar transport system ATP-binding protein | shared |
| K10112 | Environmental informatio | Saccharide, polyol, and lipid transpo | M00606 | N,N'-Diacetylchitobiose transport system [PATH:map020 | msmX, msmK, malK | multiple sugar transport system ATP-binding protein | shared |
| K11618 | Environmental informatio | Two-component regulatory system | M00481 | LiaS-LiaR (cell wall stress response) two-component reg | liaR | two-component system, NarL family, response regulator | FL-ls |

| | | | | | | | |
|---|---|---|---|---|---|---|---|
| K07703 | Environmental informatior | Two-component regulatory system | M00488 | DcuS-DcuR (C4-dicarboxylate metabolism) two-compone | dcuR | two-component system, CitB family, response regulator | FL-ls |
| K07657 | Environmental informatior | Two-component regulatory system | M00434 | PhoR-PhoB (phosphate starvation response) two-compo | phoB | two-component system, OmpR family, phosphate regulo | PA-ls |
| K07636 | Environmental informatior | Two-component regulatory system | M00434 | PhoR-PhoB (phosphate starvation response) two-compo | phoR | two-component system, OmpR family, phosphate regulo | PA-ls |
| K07776 | Environmental informatior | Two-component regulatory system | M00443 | SenX3-RegX3 (phosphate starvation response) two-com | regX3 | two-component system, OmpR family, response regulato | PA-ls |
| K07768 | Environmental informatior | Two-component regulatory system | M00443 | SenX3-RegX3 (phosphate starvation response) two-com | senX3 | two-component system, OmpR family, sensor histidine k | PA-ls |
| K07638 | Environmental informatior | Two-component regulatory system | M00445 | EnvZ-OmpR (osmotic stress response) two-component r | envZ | two-component system, OmpR family, osmolarity senso | PA-ls |
| K07659 | Environmental informatior | Two-component regulatory system | M00445 | EnvZ-OmpR (osmotic stress response) two-component r | ompR | two-component system, OmpR family, phosphate regulo | PA-ls |
| K07665 | Environmental informatior | Two-component regulatory system | M00452 | CusS-CusR (copper tolerance) two-component regulator | cusR, copR, silR | two-component system, OmpR family, copper resistance | PA-ls |
| K07644 | Environmental informatior | Two-component regulatory system | M00452 | CusS-CusR (copper tolerance) two-component regulator | cusS, copS, silS | two-component system, OmpR family, heavy metal sens | PA-ls |
| K07646 | Environmental informatior | Two-component regulatory system | M00454 | KdpD-KdpE (potassium transport) two-component regula | kdpD | two-component system, OmpR family, sensor histidine k | PA-ls |
| K07774 | Environmental informatior | Two-component regulatory system | M00457 | TctE-TctD (tricarboxylic acid transport) two-component r | tctD | two-component system, OmpR family, response regulato | PA-ls |
| K07649 | Environmental informatior | Two-component regulatory system | M00457 | TctE-TctD (tricarboxylic acid transport) two-component r | tctE | two-component system, OmpR family, sensor histidine k | PA-ls |
| K07775 | Environmental informatior | Two-component regulatory system | M00458 | ResE-ResD (aerobic and anaerobic respiration) two-com | resD | two-component system, OmpR family, response regulato | PA-ls |
| K07669 | Environmental informatior | Two-component regulatory system | M00460 | MprB-MprA (maintenance of persistent infection) two-co | mprA | two-component system, OmpR family, response regulato | PA-ls |
| K07653 | Environmental informatior | Two-component regulatory system | M00460 | MprB-MprA (maintenance of persistent infection) two-co | mprB | two-component system, OmpR family, sensor histidine k | PA-ls |
| K07670 | Environmental informatior | Two-component regulatory system | M00461 | MtrB-MtrA (osmotic stress response) two-component re | mtrA | two-component system, OmpR family, response regulato | PA-ls |
| K07654 | Environmental informatior | Two-component regulatory system | M00461 | MtrB-MtrA (osmotic stress response) two-component re | mtrB | two-component system, OmpR family, sensor histidine k | PA-ls |
| K07655 | Environmental informatior | Two-component regulatory system | M00462 | PrrB-PrrA (intracellular multiplication) two-component re | prrB | two-component system, OmpR family, sensor histidine k | PA-ls |
| K07684 | Environmental informatior | Two-component regulatory system | M00471 | NarX-NarL (nitrate respiration) two-component regulatory | narL | two-component system, NarL family, nitrate/nitrite respo | PA-ls |
| K07678 | Environmental informatior | Two-component regulatory system | M00475 | BarA-UvrY (central carbon metabolism) two-component r | barA, gacS, varS | two-component system, NarL family, sensor histidine kir | PA-ls |
| K07693 | Environmental informatior | Two-component regulatory system | M00479 | DesK-DesR (membrane lipid fluidity regulation) two-comp | desR | two-component system, NarL family, response regulator | PA-ls |
| K07778 | Environmental informatior | Two-component regulatory system | M00479 | DesK-DesR (membrane lipid fluidity regulation) two-comp | desK | two-component system, NarL family, sensor histidine kir | PA-ls |
| K11638 | Environmental informatior | Two-component regulatory system | M00487 | CitS-CitT (magnesium-citrate transport) two-component r | K11638, citT | two-component system, CitB family, response regulator | PA-ls |
| K13599 | Environmental informatior | Two-component regulatory system | M00498 | NtrY-NtrX (nitrogen regulation) two-component regulatory | ntrX | two-component system, NtrC family, nitrogen regulation | PA-ls |
| K02667 | Environmental informatior | Two-component regulatory system | M00501 | PilS-PilR (type 4 fimbriae synthesis) two-component reg | pilR, pehR | two-component system, NtrC family, response regulator | PA-ls |
| K10126 | Environmental informatior | Two-component regulatory system | M00504 | DctB-DctD (C4-dicarboxylate transport) two-component r | dctD | two-component system, NtrC family, C4-dicarboxylate tr | PA-ls |
| K10125 | Environmental informatior | Two-component regulatory system | M00504 | DctB-DctD (C4-dicarboxylate transport) two-component r | dctB | two-component system, NtrC family, C4-dicarboxylate tr | PA-ls |
| K13490 | Environmental informatior | Two-component regulatory system | M00509 | WspE-WspRF (chemosensory) two-component regulator | wspE | two-component system, chemotaxis family, sensor histi | PA-ls |
| K07658 | Environmental informatior | Two-component regulatory system | M00434 | PhoR-PhoB (phosphate starvation response) two-compo | phoB1, phoP | two-component system, OmpR family, alkaline phosphat | shared |
| K07661 | Environmental informatior | Two-component regulatory system | M00446 | RstB-RstA two-component regulatory system [PATH:ma | rstA | two-component system, OmpR family, response regulato | shared |
| K07639 | Environmental informatior | Two-component regulatory system | M00446 | RstB-RstA two-component regulatory system [PATH:ma | rstB | two-component system, OmpR family, sensor histidine k | shared |
| K07663 | Environmental informatior | Two-component regulatory system | M00449 | CreC-CreB (phosphate regulation) two-component regula | creB | two-component system, OmpR family, catabolic regulati | shared |
| K07641 | Environmental informatior | Two-component regulatory system | M00449 | CreC-CreB (phosphate regulation) two-component regula | creC | two-component system, OmpR family, sensor histidine k | shared |
| K07664 | Environmental informatior | Two-component regulatory system | M00450 | BaeS-BaeR (envelope stress response) two-component | baeR, smeR | two-component system, OmpR family, response regulato | shared |
| K07642 | Environmental informatior | Two-component regulatory system | M00450 | BaeS-BaeR (envelope stress response) two-component | baeS, smeS | two-component system, OmpR family, sensor histidine k | shared |
| K07647 | Environmental informatior | Two-component regulatory system | M00455 | TorS-TorR (TMAO respiration) two-component regulatory | torS | two-component system, OmpR family, sensor histidine k | shared |
| K07772 | Environmental informatior | Two-component regulatory system | M00455 | TorS-TorR (TMAO respiration) two-component regulatory | torR | two-component system, OmpR family, torCAD operon re | shared |
| K07773 | Environmental informatior | Two-component regulatory system | M00456 | ArcB-ArcA (anoxic redox control) two-component regulat | arcA | two-component system, OmpR family, aerobic respiratio | shared |
| K07648 | Environmental informatior | Two-component regulatory system | M00456 | ArcB-ArcA (anoxic redox control) two-component regulat | arcB | two-component system, OmpR family, aerobic respiratio | shared |
| K07651 | Environmental informatior | Two-component regulatory system | M00458 | ResE-ResD (aerobic and anaerobic respiration) two-com | resE | two-component system, OmpR family, sensor histidine k | shared |
| K07668 | Environmental informatior | Two-component regulatory system | M00459 | VicK-VicR (cell wall metabolism) two-component regulato | vicR | two-component system, OmpR family, response regulato | shared |
| K07652 | Environmental informatior | Two-component regulatory system | M00459 | VicK-VicR (cell wall metabolism) two-component regulato | vicK | two-component system, OmpR family, sensor histidine k | shared |
| K07671 | Environmental informatior | Two-component regulatory system | M00462 | PrrB-PrrA (intracellular multiplication) two-component re | prrA | two-component system, OmpR family, response regulato | shared |
| K07656 | Environmental informatior | Two-component regulatory system | M00463 | TrcS-TrcR two-component regulatory system [PATH:map | trcS | two-component system, OmpR family, sensor histidine k | shared |
| K11521 | Environmental informatior | Two-component regulatory system | M00465 | ManS-ManR (manganese homeostasis) two-component r | K11521, manR | two-component system, OmpR family, manganese sensi | shared |
| K11520 | Environmental informatior | Two-component regulatory system | M00465 | ManS-ManR (manganese homeostasis) two-component r | manS | two-component system, OmpR family, manganese sensi | shared |
| K07769 | Environmental informatior | Two-component regulatory system | M00466 | NblS-NblR (photosynthesis) two-component regulatory s | nblS | two-component system, OmpR family, sensor histidine k | shared |
| K10697 | Environmental informatior | Two-component regulatory system | M00467 | SasA-RpaAB (circadian timing mediating) two-componen | rpaA | two-component system, OmpR family, response regulato | shared |
| K11329 | Environmental informatior | Two-component regulatory system | M00467 | SasA-RpaAB (circadian timing mediating) two-componen | rpaB | two-component system, OmpR family, response regulato | shared |
| K07673 | Environmental informatior | Two-component regulatory system | M00471 | NarX-NarL (nitrate respiration) two-component regulatory | narX | two-component system, NarL family, nitrate/nitrite senso | shared |
| K07685 | Environmental informatior | Two-component regulatory system | M00472 | NarQ-NarP (nitrate respiration) two-component regulator | narP | two-component system, NarL family, nitrate/nitrite respo | shared |
| K07675 | Environmental informatior | Two-component regulatory system | M00473 | UhpB-UhpA (hexose phosphates uptake) two-componen | uhpB | two-component system, NarL family, sensor histidine kir | shared |
| K07686 | Environmental informatior | Two-component regulatory system | M00473 | UhpB-UhpA (hexose phosphates uptake) two-componen | uhpA | two-component system, NarL family, uhpT operon respo | shared |
| K07677 | Environmental informatior | Two-component regulatory system | M00474 | RcsC-RcsD-RcsB (capsule synthesis) two-component re | rcsC | two-component system, NarL family, capsular synthesis | shared |
| K07687 | Environmental informatior | Two-component regulatory system | M00474 | RcsC-RcsD-RcsB (capsule synthesis) two-component re | rcsB | two-component system, NarL family, captular synthesis | shared |
| K07689 | Environmental informatior | Two-component regulatory system | M00475 | BarA-UvrY (central carbon metabolism) two-component r | uvrY, gacA, varA | two-component system, NarL family, invasion response | shared |
| K07777 | Environmental informatior | Two-component regulatory system | M00478 | DegS-DegU (multicellular behavior control) two-compone | degS | two-component system, NarL family, sensor histidine kir | shared |
| K11617 | Environmental informatior | Two-component regulatory system | M00481 | LiaS-LiaR (cell wall stress response) two-component reg | liaS | two-component system, NarL family, sensor histidine kir | shared |
| K07695 | Environmental informatior | Two-component regulatory system | M00482 | DevS-DevR (redox response) two-component regulatory | devR | two-component system, NarL family, response regulator | shared |
| K07682 | Environmental informatior | Two-component regulatory system | M00482 | DevS-DevR (redox response) two-component regulatory | devS | two-component system, NarL family, sensor histidine kir | shared |

| | | | | | | | |
|---|---|---|---|---|---|---|---|
| K07696 | Environmental informatior | Two-component regulatory system | M00483 | NreB-NreC (dissimilatory nitrate/nitrite reduction) two-co | nreC | two-component system, NarL family, response regulator | shared |
| K07683 | Environmental informatior | Two-component regulatory system | M00483 | NreB-NreC (dissimilatory nitrate/nitrite reduction) two-co | nreB | two-component system, NarL family, sensor histidine kir | shared |
| K11624 | Environmental informatior | Two-component regulatory system | M00484 | YdfH-YdfI two-component regulatory system [PATH:map | ydfI | two-component system, NarL family, response regulator | shared |
| K07697 | Environmental informatior | Two-component regulatory system | M00485 | KinABCDE-Spo0FA (sporulation control) two-component | kinB | two-component system, sporulation sensor kinase B [E( | shared |
| K07698 | Environmental informatior | Two-component regulatory system | M00485 | KinABCDE-Spo0FA (sporulation control) two-component | kinC | two-component system, sporulation sensor kinase C [E( | shared |
| K13533 | Environmental informatior | Two-component regulatory system | M00485 | KinABCDE-Spo0FA (sporulation control) two-component | kinE | two-component system, sporulation sensor kinase E [E( | shared |
| K11637 | Environmental informatior | Two-component regulatory system | M00487 | CitS-CitT (magnesium-citrate transport) two-component r | citS | two-component system, CitB family, sensor histidine kin | shared |
| K07701 | Environmental informatior | Two-component regulatory system | M00488 | DcuS-DcuR (C4-dicarboxylate metabolism) two-compone | dcuS | two-component system, CitB family, sensor histidine kin | shared |
| K11615 | Environmental informatior | Two-component regulatory system | M00490 | MalK-MalR (malate transport) two-component regulatory | malR | two-component system, CitB family, response regulator | shared |
| K07705 | Environmental informatior | Two-component regulatory system | M00492 | LytS-LytR two-component regulatory system [PATH:map | lytT, lytR | two-component system, LytTR family, response regulatc | shared |
| K07704 | Environmental informatior | Two-component regulatory system | M00492 | LytS-LytR two-component regulatory system [PATH:map | lytS | two-component system, LytTR family, sensor histidine k | shared |
| K08083 | Environmental informatior | Two-component regulatory system | M00493 | AlgZ-AlgR (alginate production) two-component regulator | algR | two-component system, LytTR family, response regulatc | shared |
| K08082 | Environmental informatior | Two-component regulatory system | M00493 | AlgZ-AlgR (alginate production) two-component regulator | algZ | two-component system, LytTR family, sensor histidine k | shared |
| K07712 | Environmental informatior | Two-component regulatory system | M00497 | GlnL-GlnG (nitrogen regulation) two-component regulato | glnG, ntrC | two-component system, NtrC family, nitrogen regulation | shared |
| K07708 | Environmental informatior | Two-component regulatory system | M00497 | GlnL-GlnG (nitrogen regulation) two-component regulato | glnL, ntrB | two-component system, NtrC family, nitrogen regulation | shared |
| K13598 | Environmental informatior | Two-component regulatory system | M00498 | NtrY-NtrX (nitrogen regulation) two-component regulatory | ntrY | two-component system, NtrC family, nitrogen regulation | shared |
| K07713 | Environmental informatior | Two-component regulatory system | M00499 | HydH-HydG (metal tolerance) two-component regulatory | zraR, hydG | two-component system, NtrC family, response regulator | shared |
| K07709 | Environmental informatior | Two-component regulatory system | M00499 | HydH-HydG (metal tolerance) two-component regulatory | zraS, hydH | two-component system, NtrC family, sensor histidine kir | shared |
| K07714 | Environmental informatior | Two-component regulatory system | M00500 | AtoS-AtoC (cPHB biosynthesis) two-component regulato | atoC | two-component system, NtrC family, response regulator | shared |
| K07710 | Environmental informatior | Two-component regulatory system | M00500 | AtoS-AtoC (cPHB biosynthesis) two-component regulato | atoS | two-component system, NtrC family, sensor histidine kir | shared |
| K02668 | Environmental informatior | Two-component regulatory system | M00501 | PilS-PilR (type 4 fimbriae synthesis) two-component reg | pilS, pehS | two-component system, NtrC family, sensor histidine kir | shared |
| K08475 | Environmental informatior | Two-component regulatory system | M00503 | PgtB-PgtA (phosphoglycerate transport) two-component | pgtB | two-component system, NtrC family, phosphoglycerate t | shared |
| K11384 | Environmental informatior | Two-component regulatory system | M00505 | KinB-AlgB (alginate production) two-component regulato | algB | two-component system, NtrC family, response regulator | shared |
| K11383 | Environmental informatior | Two-component regulatory system | M00505 | KinB-AlgB (alginate production) two-component regulato | kinB | two-component system, NtrC family, sensor histidine kir | shared |
| K06596 | Environmental informatior | Two-component regulatory system | M00507 | ChpA-ChpB/PilGH (chemosensory) two-component regul | chpA | chemosensory pili system protein ChpA (sensor histidin | shared |
| K06597 | Environmental informatior | Two-component regulatory system | M00507 | ChpA-ChpB/PilGH (chemosensory) two-component regul | chpB | chemosensory pili system protein ChpB (putative protei | shared |
| K02657 | Environmental informatior | Two-component regulatory system | M00507 | ChpA-ChpB/PilGH (chemosensory) two-component regul | pilG | twitching motility two-component system response regul | shared |
| K02658 | Environmental informatior | Two-component regulatory system | M00507 | ChpA-ChpB/PilGH (chemosensory) two-component regul | pilH | twitching motility two-component system response regul | shared |
| K02487 | Environmental informatior | Two-component regulatory system | M00507 | ChpA-ChpB/PilGH (chemosensory) two-component regul | pilL | type IV pili sensor histidine kinase and response regulat | shared |
| K11444 | Environmental informatior | Two-component regulatory system | M00509 | WspE-WspRF (chemosensory) two-component regulatory | wspR | two-component system, chemotaxis family, response re | shared |
| K11354 | Environmental informatior | Two-component regulatory system | M00510 | Cph1-Rcp1 (light response) two-component regulatory s | cph1 | two-component system, chemotaxis family, sensor kinas | shared |
| K13040 | Environmental informatior | Two-component regulatory system | M00514 | TtrS-TtrR (tetrathionate respiration) two-component regu | ttrS | two-component system, LuxR family, sensor histidine ki | shared |
| K07720 | Environmental informatior | Two-component regulatory system | M00519 | YesM-YesN two-component regulatory system [PATH:ma | yesN | two-component system, response regulator YesN | shared |
| K07718 | Environmental informatior | Two-component regulatory system | M00519 | YesM-YesN two-component regulatory system [PATH:ma | yesM | two-component system, sensor histidine kinase YesM [E | shared |
| K14981 | Environmental informatior | Two-component regulatory system | M00520 | ChvG-ChvI (acidity sensing) two-component regulatory s | chvI | two-component system, OmpR family, response regulatc | shared |
| K14980 | Environmental informatior | Two-component regulatory system | M00520 | ChvG-ChvI (acidity sensing) two-component regulatory s | chvG | two-component system, OmpR family, sensor histidine k | shared |
| K15012 | Environmental informatior | Two-component regulatory system | M00523 | RegB-RegA (redox response) two-component regulatory | regA, regR, actR | two-component system, response regulator RegA | shared |
| K15011 | Environmental informatior | Two-component regulatory system | M00523 | RegB-RegA (redox response) two-component regulatory | regB, regS, actS | two-component system, sensor histidine kinase RegB [E | shared |
| K14987 | Environmental informatior | Two-component regulatory system | M00524 | FixL-FixJ (nitrogen fixation) two-component regulatory sy | fixJ | two-component system, LuxR family, response regulato | shared |
| K14986 | Environmental informatior | Two-component regulatory system | M00524 | FixL-FixJ (nitrogen fixation) two-component regulatory sy | fixL | two-component system, LuxR family, sensor kinase FixL | shared |
| K02489 | Environmental informatior | Two-component regulatory system | M00662 | Hk1-Rrp1 (glycerol uptake and utilization) two-componer | hk1 | two-component system, glycerol uptake and utilization s | shared |
| K07665 | Gene set | Drug resistance | M00745 | Imipenem resistance, repression of porin OprD | cusR, copR, silR | two-component system, OmpR family, copper resistance | PA-ls |
| K07644 | Gene set | Drug resistance | M00745 | Imipenem resistance, repression of porin OprD | cusS, copS, silS | two-component system, OmpR family, heavy metal sens | PA-ls |
| K03224 | Gene set | Pathogenicity | M00542 | EHEC/EPEC pathogenicity signature, T3SS and effector | yscN, sctN, hrcN, s | ATP synthase in type III secretion protein N [EC:3.6.3.5 | shared |
| K03219 | Gene set | Pathogenicity | M00542 | EHEC/EPEC pathogenicity signature, T3SS and effector | yscC, sctC, ssaC | type III secretion protein C | shared |
| K03222 | Gene set | Pathogenicity | M00542 | EHEC/EPEC pathogenicity signature, T3SS and effector | yscJ, sctJ, hrcJ, ss | type III secretion protein J | shared |
| K03226 | Gene set | Pathogenicity | M00542 | EHEC/EPEC pathogenicity signature, T3SS and effector | yscR, sctR, hrcR, s | type III secretion protein R | shared |
| K03228 | Gene set | Pathogenicity | M00542 | EHEC/EPEC pathogenicity signature, T3SS and effector | yscT, sctT, hrcT, ss | type III secretion protein T | shared |
| K03229 | Gene set | Pathogenicity | M00542 | EHEC/EPEC pathogenicity signature, T3SS and effector | yscU, sctU, hrcU, s | type III secretion protein U | shared |
| K03230 | Gene set | Pathogenicity | M00542 | EHEC/EPEC pathogenicity signature, T3SS and effector | yscV, sctV, hrcV, s | type III secretion protein V | shared |
| K10954 | Gene set | Pathogenicity | M00850 | Vibrio cholerae pathogenicity signature, cholera toxins [ | zot | zona occludens toxin | shared |
| K03224 | Gene set | Plant pathogenicity | M00660 | Xanthomonas spp. pathogenicity signature, T3SS and e | yscN, sctN, hrcN, s | ATP synthase in type III secretion protein N [EC:3.6.3.5 | shared |
| K03222 | Gene set | Plant pathogenicity | M00660 | Xanthomonas spp. pathogenicity signature, T3SS and e | yscJ, sctJ, hrcJ, ss | type III secretion protein J | shared |
| K03223 | Gene set | Plant pathogenicity | M00660 | Xanthomonas spp. pathogenicity signature, T3SS and e | yscL, sctL | type III secretion protein L | shared |
| K03226 | Gene set | Plant pathogenicity | M00660 | Xanthomonas spp. pathogenicity signature, T3SS and e | yscR, sctR, hrcR, s | type III secretion protein R | shared |
| K03228 | Gene set | Plant pathogenicity | M00660 | Xanthomonas spp. pathogenicity signature, T3SS and e | yscT, sctT, hrcT, ss | type III secretion protein T | shared |
| K03229 | Gene set | Plant pathogenicity | M00660 | Xanthomonas spp. pathogenicity signature, T3SS and e | yscU, sctU, hrcU, s | type III secretion protein U | shared |
| K03230 | Gene set | Plant pathogenicity | M00660 | Xanthomonas spp. pathogenicity signature, T3SS and e | yscV, sctV, hrcV, s | type III secretion protein V | shared |
| K03432 | Genetic information proc | Proteasome | M00342 | Bacterial proteasome [PATH:map03050] [BR:ko03051] | psmA, prcA | proteasome alpha subunit [EC:3.4.25.1] | FL-ls |
| K03433 | Genetic information proc | Proteasome | M00342 | Bacterial proteasome [PATH:map03050] [BR:ko03051] | psmB, prcB | proteasome beta subunit [EC:3.4.25.1] | FL-ls |

| | | | | | | | |
|---|---|---|---|---|---|---|---|
| K03432 | Genetic information proc | Proteasome | M00343 | Archaeal proteasome [PATH:map03050] [BR:ko03051] | psmA, prcA | proteasome alpha subunit [EC:3.4.25.1] | FL-ls |
| K03433 | Genetic information proc | Proteasome | M00343 | Archaeal proteasome [PATH:map03050] [BR:ko03051] | psmB, prcB | proteasome beta subunit [EC:3.4.25.1] | FL-ls |
| K03420 | Genetic information proc | Proteasome | M00343 | Archaeal proteasome [PATH:map03050] [BR:ko03051] | psmR | proteasome regulatory subunit | FL-ls |
| K13527 | Genetic information proc | Proteasome | M00342 | Bacterial proteasome [PATH:map03050] [BR:ko03051] | mpa | proteasome-associated ATPase | PA-ls |
| K02729 | Genetic information proc | Proteasome | M00337 | Immunoproteasome [PATH:map03050] [BR:ko03051] | PSMA5 | 20S proteasome subunit alpha 5 [EC:3.4.25.1] | shared |
| K02729 | Genetic information proc | Proteasome | M00340 | Proteasome, 20S core particle [PATH:map03050] [BR:ko | PSMA5 | 20S proteasome subunit alpha 5 [EC:3.4.25.1] | shared |
| K14007 | Genetic information proc | Protein processing | M00404 | COPII complex [PATH:map04141] | SEC24 | protein transport protein SEC24 | shared |
| K10896 | Genetic information proc | Repair system | M00413 | FA core complex [PATH:map03460] [BR:ko03400] | FANCM | fanconi anemia group M protein | FL-ls |
| K10771 | Genetic information proc | Repair system | M00296 | BER complex [PATH:map03410] [BR:ko03400] | APEX1 | AP endonuclease 1 [EC:4.2.99.18] | shared |
| K10884 | Genetic information proc | Repair system | M00297 | DNA-PK complex [PATH:map03450 map04110] [BR:ko03 | XRCC6, KU70, G22I | ATP-dependent DNA helicase 2 subunit 1 | shared |
| K02866 | Genetic information proc | Ribosome | M00177 | Ribosome, eukaryotes [PATH:map03010] [BR:ko03011] | RP-L10e, RPL10 | large subunit ribosomal protein L10e | FL-ls |
| K02877 | Genetic information proc | Ribosome | M00177 | Ribosome, eukaryotes [PATH:map03010] [BR:ko03011] | RP-L15e, RPL15 | large subunit ribosomal protein L15e | FL-ls |
| K02883 | Genetic information proc | Ribosome | M00177 | Ribosome, eukaryotes [PATH:map03010] [BR:ko03011] | RP-L18e, RPL18 | large subunit ribosomal protein L18e | FL-ls |
| K02885 | Genetic information proc | Ribosome | M00177 | Ribosome, eukaryotes [PATH:map03010] [BR:ko03011] | RP-L19e, RPL19 | large subunit ribosomal protein L19e | FL-ls |
| K02889 | Genetic information proc | Ribosome | M00177 | Ribosome, eukaryotes [PATH:map03010] [BR:ko03011] | RP-L21e, RPL21 | large subunit ribosomal protein L21e | FL-ls |
| K02896 | Genetic information proc | Ribosome | M00177 | Ribosome, eukaryotes [PATH:map03010] [BR:ko03011] | RP-L24e, RPL24 | large subunit ribosomal protein L24e | FL-ls |
| K02910 | Genetic information proc | Ribosome | M00177 | Ribosome, eukaryotes [PATH:map03010] [BR:ko03011] | RP-L31e, RPL31 | large subunit ribosomal protein L31e | FL-ls |
| K02912 | Genetic information proc | Ribosome | M00177 | Ribosome, eukaryotes [PATH:map03010] [BR:ko03011] | RP-L32e, RPL32 | large subunit ribosomal protein L32e | FL-ls |
| K02921 | Genetic information proc | Ribosome | M00177 | Ribosome, eukaryotes [PATH:map03010] [BR:ko03011] | RP-L37Ae, RPL37A | large subunit ribosomal protein L37Ae | FL-ls |
| K02924 | Genetic information proc | Ribosome | M00177 | Ribosome, eukaryotes [PATH:map03010] [BR:ko03011] | RP-L39e, RPL39 | large subunit ribosomal protein L39e | FL-ls |
| K02929 | Genetic information proc | Ribosome | M00177 | Ribosome, eukaryotes [PATH:map03010] [BR:ko03011] | RP-L44e, RPL44 | large subunit ribosomal protein L44e | FL-ls |
| K02936 | Genetic information proc | Ribosome | M00177 | Ribosome, eukaryotes [PATH:map03010] [BR:ko03011] | RP-L7Ae, RPL7A | large subunit ribosomal protein L7Ae | FL-ls |
| K02962 | Genetic information proc | Ribosome | M00177 | Ribosome, eukaryotes [PATH:map03010] [BR:ko03011] | RP-S17e, RPS17 | small subunit ribosomal protein S17e | FL-ls |
| K02975 | Genetic information proc | Ribosome | M00177 | Ribosome, eukaryotes [PATH:map03010] [BR:ko03011] | RP-S25e, RPS25 | small subunit ribosomal protein S25e | FL-ls |
| K02976 | Genetic information proc | Ribosome | M00177 | Ribosome, eukaryotes [PATH:map03010] [BR:ko03011] | RP-S26e, RPS26 | small subunit ribosomal protein S26e | FL-ls |
| K02977 | Genetic information proc | Ribosome | M00177 | Ribosome, eukaryotes [PATH:map03010] [BR:ko03011] | RP-S27Ae, RPS27A | small subunit ribosomal protein S27Ae | FL-ls |
| K02978 | Genetic information proc | Ribosome | M00177 | Ribosome, eukaryotes [PATH:map03010] [BR:ko03011] | RP-S27e, RPS27 | small subunit ribosomal protein S27e | FL-ls |
| K02979 | Genetic information proc | Ribosome | M00177 | Ribosome, eukaryotes [PATH:map03010] [BR:ko03011] | RP-S28e, RPS28 | small subunit ribosomal protein S28e | FL-ls |
| K02983 | Genetic information proc | Ribosome | M00177 | Ribosome, eukaryotes [PATH:map03010] [BR:ko03011] | RP-S30e, RPS30 | small subunit ribosomal protein S30e | FL-ls |
| K02984 | Genetic information proc | Ribosome | M00177 | Ribosome, eukaryotes [PATH:map03010] [BR:ko03011] | RP-S3Ae, RPS3A | small subunit ribosomal protein S3Ae | FL-ls |
| K02987 | Genetic information proc | Ribosome | M00177 | Ribosome, eukaryotes [PATH:map03010] [BR:ko03011] | RP-S4e, RPS4 | small subunit ribosomal protein S4e | FL-ls |
| K02995 | Genetic information proc | Ribosome | M00177 | Ribosome, eukaryotes [PATH:map03010] [BR:ko03011] | RP-S8e, RPS8 | small subunit ribosomal protein S8e | FL-ls |
| K02863 | Genetic information proc | Ribosome | M00178 | Ribosome, bacteria [PATH:map03010] [BR:ko03011] | RP-L1, MRPL1, rplA | large subunit ribosomal protein L1 | FL-ls |
| K02864 | Genetic information proc | Ribosome | M00178 | Ribosome, bacteria [PATH:map03010] [BR:ko03011] | RP-L10, MRPL10, r | large subunit ribosomal protein L10 | FL-ls |
| K02867 | Genetic information proc | Ribosome | M00178 | Ribosome, bacteria [PATH:map03010] [BR:ko03011] | RP-L11, MRPL11, r | large subunit ribosomal protein L11 | FL-ls |
| K02871 | Genetic information proc | Ribosome | M00178 | Ribosome, bacteria [PATH:map03010] [BR:ko03011] | RP-L13, MRPL13, r | large subunit ribosomal protein L13 | FL-ls |
| K02874 | Genetic information proc | Ribosome | M00178 | Ribosome, bacteria [PATH:map03010] [BR:ko03011] | RP-L14, MRPL14, r | large subunit ribosomal protein L14 | FL-ls |
| K02876 | Genetic information proc | Ribosome | M00178 | Ribosome, bacteria [PATH:map03010] [BR:ko03011] | RP-L15, MRPL15, r | large subunit ribosomal protein L15 | FL-ls |
| K02881 | Genetic information proc | Ribosome | M00178 | Ribosome, bacteria [PATH:map03010] [BR:ko03011] | RP-L18, MRPL18, r | large subunit ribosomal protein L18 | FL-ls |
| K02886 | Genetic information proc | Ribosome | M00178 | Ribosome, bacteria [PATH:map03010] [BR:ko03011] | RP-L2, MRPL2, rplB | large subunit ribosomal protein L2 | FL-ls |
| K02890 | Genetic information proc | Ribosome | M00178 | Ribosome, bacteria [PATH:map03010] [BR:ko03011] | RP-L22, MRPL22, r | large subunit ribosomal protein L22 | FL-ls |
| K02892 | Genetic information proc | Ribosome | M00178 | Ribosome, bacteria [PATH:map03010] [BR:ko03011] | RP-L23, MRPL23, r | large subunit ribosomal protein L23 | FL-ls |
| K02895 | Genetic information proc | Ribosome | M00178 | Ribosome, bacteria [PATH:map03010] [BR:ko03011] | RP-L24, MRPL24, r | large subunit ribosomal protein L24 | FL-ls |
| K02904 | Genetic information proc | Ribosome | M00178 | Ribosome, bacteria [PATH:map03010] [BR:ko03011] | RP-L29, rpmC | large subunit ribosomal protein L29 | FL-ls |
| K02906 | Genetic information proc | Ribosome | M00178 | Ribosome, bacteria [PATH:map03010] [BR:ko03011] | RP-L3, MRPL3, rplC | large subunit ribosomal protein L3 | FL-ls |
| K02907 | Genetic information proc | Ribosome | M00178 | Ribosome, bacteria [PATH:map03010] [BR:ko03011] | RP-L30, MRPL30, r | large subunit ribosomal protein L30 | FL-ls |
| K02931 | Genetic information proc | Ribosome | M00178 | Ribosome, bacteria [PATH:map03010] [BR:ko03011] | RP-L5, MRPL5, rplE | large subunit ribosomal protein L5 | FL-ls |
| K02933 | Genetic information proc | Ribosome | M00178 | Ribosome, bacteria [PATH:map03010] [BR:ko03011] | RP-L6, MRPL6, rplF | large subunit ribosomal protein L6 | FL-ls |
| K02946 | Genetic information proc | Ribosome | M00178 | Ribosome, bacteria [PATH:map03010] [BR:ko03011] | RP-S10, MRPS10, r | small subunit ribosomal protein S10 | FL-ls |
| K02948 | Genetic information proc | Ribosome | M00178 | Ribosome, bacteria [PATH:map03010] [BR:ko03011] | RP-S11, MRPS11, r | small subunit ribosomal protein S11 | FL-ls |
| K02950 | Genetic information proc | Ribosome | M00178 | Ribosome, bacteria [PATH:map03010] [BR:ko03011] | RP-S12, MRPS12, r | small subunit ribosomal protein S12 | FL-ls |
| K02952 | Genetic information proc | Ribosome | M00178 | Ribosome, bacteria [PATH:map03010] [BR:ko03011] | RP-S13, rpsM | small subunit ribosomal protein S13 | FL-ls |
| K02956 | Genetic information proc | Ribosome | M00178 | Ribosome, bacteria [PATH:map03010] [BR:ko03011] | RP-S15, MRPS15, r | small subunit ribosomal protein S15 | FL-ls |
| K02961 | Genetic information proc | Ribosome | M00178 | Ribosome, bacteria [PATH:map03010] [BR:ko03011] | RP-S17, MRPS17, r | small subunit ribosomal protein S17 | FL-ls |
| K02965 | Genetic information proc | Ribosome | M00178 | Ribosome, bacteria [PATH:map03010] [BR:ko03011] | RP-S19, rpsS | small subunit ribosomal protein S19 | FL-ls |
| K02967 | Genetic information proc | Ribosome | M00178 | Ribosome, bacteria [PATH:map03010] [BR:ko03011] | RP-S2, MRPS2, rps | small subunit ribosomal protein S2 | FL-ls |
| K02982 | Genetic information proc | Ribosome | M00178 | Ribosome, bacteria [PATH:map03010] [BR:ko03011] | RP-S3, rpsC | small subunit ribosomal protein S3 | FL-ls |
| K02986 | Genetic information proc | Ribosome | M00178 | Ribosome, bacteria [PATH:map03010] [BR:ko03011] | RP-S4, rpsD | small subunit ribosomal protein S4 | FL-ls |
| K02988 | Genetic information proc | Ribosome | M00178 | Ribosome, bacteria [PATH:map03010] [BR:ko03011] | RP-S5, MRPS5, rps | small subunit ribosomal protein S5 | FL-ls |
| K02992 | Genetic information proc | Ribosome | M00178 | Ribosome, bacteria [PATH:map03010] [BR:ko03011] | RP-S7, MRPS7, rps | small subunit ribosomal protein S7 | FL-ls |

| | | | | | | | |
|---|---|---|---|---|---|---|---|
| K02994 | Genetic information proc | Ribosome | M00178 | Ribosome, bacteria [PATH:map03010] [BR:ko03011] | RP-S8, rpsH | small subunit ribosomal protein S8 | FL-ls |
| K02996 | Genetic information proc | Ribosome | M00178 | Ribosome, bacteria [PATH:map03010] [BR:ko03011] | RP-S9, MRPS9, rps | small subunit ribosomal protein S9 | FL-ls |
| K02863 | Genetic information proc | Ribosome | M00179 | Ribosome, archaea [PATH:map03010] [BR:ko03011] | RP-L1, MRPL1, rplA | large subunit ribosomal protein L1 | FL-ls |
| K02864 | Genetic information proc | Ribosome | M00179 | Ribosome, archaea [PATH:map03010] [BR:ko03011] | RP-L10, MRPL10, r | large subunit ribosomal protein L10 | FL-ls |
| K02866 | Genetic information proc | Ribosome | M00179 | Ribosome, archaea [PATH:map03010] [BR:ko03011] | RP-L10e, RPL10 | large subunit ribosomal protein L10e | FL-ls |
| K02867 | Genetic information proc | Ribosome | M00179 | Ribosome, archaea [PATH:map03010] [BR:ko03011] | RP-L11, MRPL11, r | large subunit ribosomal protein L11 | FL-ls |
| K02869 | Genetic information proc | Ribosome | M00179 | Ribosome, archaea [PATH:map03010] [BR:ko03011] | RP-L12, rpl12 | large subunit ribosomal protein L12 | FL-ls |
| K02871 | Genetic information proc | Ribosome | M00179 | Ribosome, archaea [PATH:map03010] [BR:ko03011] | RP-L13, MRPL13, r | large subunit ribosomal protein L13 | FL-ls |
| K02874 | Genetic information proc | Ribosome | M00179 | Ribosome, archaea [PATH:map03010] [BR:ko03011] | RP-L14, MRPL14, r | large subunit ribosomal protein L14 | FL-ls |
| K02876 | Genetic information proc | Ribosome | M00179 | Ribosome, archaea [PATH:map03010] [BR:ko03011] | RP-L15, MRPL15, r | large subunit ribosomal protein L15 | FL-ls |
| K02877 | Genetic information proc | Ribosome | M00179 | Ribosome, archaea [PATH:map03010] [BR:ko03011] | RP-L15e, RPL15 | large subunit ribosomal protein L15e | FL-ls |
| K02881 | Genetic information proc | Ribosome | M00179 | Ribosome, archaea [PATH:map03010] [BR:ko03011] | RP-L18, MRPL18, r | large subunit ribosomal protein L18 | FL-ls |
| K02883 | Genetic information proc | Ribosome | M00179 | Ribosome, archaea [PATH:map03010] [BR:ko03011] | RP-L18e, RPL18 | large subunit ribosomal protein L18e | FL-ls |
| K02885 | Genetic information proc | Ribosome | M00179 | Ribosome, archaea [PATH:map03010] [BR:ko03011] | RP-L19e, RPL19 | large subunit ribosomal protein L19e | FL-ls |
| K02886 | Genetic information proc | Ribosome | M00179 | Ribosome, archaea [PATH:map03010] [BR:ko03011] | RP-L2, MRPL2, rplB | large subunit ribosomal protein L2 | FL-ls |
| K02889 | Genetic information proc | Ribosome | M00179 | Ribosome, archaea [PATH:map03010] [BR:ko03011] | RP-L21e, RPL21 | large subunit ribosomal protein L21e | FL-ls |
| K02890 | Genetic information proc | Ribosome | M00179 | Ribosome, archaea [PATH:map03010] [BR:ko03011] | RP-L22, MRPL22, r | large subunit ribosomal protein L22 | FL-ls |
| K02892 | Genetic information proc | Ribosome | M00179 | Ribosome, archaea [PATH:map03010] [BR:ko03011] | RP-L23, MRPL23, r | large subunit ribosomal protein L23 | FL-ls |
| K02895 | Genetic information proc | Ribosome | M00179 | Ribosome, archaea [PATH:map03010] [BR:ko03011] | RP-L24, MRPL24, r | large subunit ribosomal protein L24 | FL-ls |
| K02896 | Genetic information proc | Ribosome | M00179 | Ribosome, archaea [PATH:map03010] [BR:ko03011] | RP-L24e, RPL24 | large subunit ribosomal protein L24e | FL-ls |
| K02904 | Genetic information proc | Ribosome | M00179 | Ribosome, archaea [PATH:map03010] [BR:ko03011] | RP-L29, rpmC | large subunit ribosomal protein L29 | FL-ls |
| K02906 | Genetic information proc | Ribosome | M00179 | Ribosome, archaea [PATH:map03010] [BR:ko03011] | RP-L3, MRPL3, rplC | large subunit ribosomal protein L3 | FL-ls |
| K02907 | Genetic information proc | Ribosome | M00179 | Ribosome, archaea [PATH:map03010] [BR:ko03011] | RP-L30, MRPL30, r | large subunit ribosomal protein L30 | FL-ls |
| K02910 | Genetic information proc | Ribosome | M00179 | Ribosome, archaea [PATH:map03010] [BR:ko03011] | RP-L31e, RPL31 | large subunit ribosomal protein L31e | FL-ls |
| K02912 | Genetic information proc | Ribosome | M00179 | Ribosome, archaea [PATH:map03010] [BR:ko03011] | RP-L32e, RPL32 | large subunit ribosomal protein L32e | FL-ls |
| K02921 | Genetic information proc | Ribosome | M00179 | Ribosome, archaea [PATH:map03010] [BR:ko03011] | RP-L37Ae, RPL37A | large subunit ribosomal protein L37Ae | FL-ls |
| K02924 | Genetic information proc | Ribosome | M00179 | Ribosome, archaea [PATH:map03010] [BR:ko03011] | RP-L39e, RPL39 | large subunit ribosomal protein L39e | FL-ls |
| K02929 | Genetic information proc | Ribosome | M00179 | Ribosome, archaea [PATH:map03010] [BR:ko03011] | RP-L44e, RPL44 | large subunit ribosomal protein L44e | FL-ls |
| K02931 | Genetic information proc | Ribosome | M00179 | Ribosome, archaea [PATH:map03010] [BR:ko03011] | RP-L5, MRPL5, rplE | large subunit ribosomal protein L5 | FL-ls |
| K02933 | Genetic information proc | Ribosome | M00179 | Ribosome, archaea [PATH:map03010] [BR:ko03011] | RP-L6, MRPL6, rplF | large subunit ribosomal protein L6 | FL-ls |
| K02936 | Genetic information proc | Ribosome | M00179 | Ribosome, archaea [PATH:map03010] [BR:ko03011] | RP-L7Ae, RPL7A | large subunit ribosomal protein L7Ae | FL-ls |
| K02946 | Genetic information proc | Ribosome | M00179 | Ribosome, archaea [PATH:map03010] [BR:ko03011] | RP-S10, MRPS10, r | small subunit ribosomal protein S10 | FL-ls |
| K02948 | Genetic information proc | Ribosome | M00179 | Ribosome, archaea [PATH:map03010] [BR:ko03011] | RP-S11, MRPS11, r | small subunit ribosomal protein S11 | FL-ls |
| K02950 | Genetic information proc | Ribosome | M00179 | Ribosome, archaea [PATH:map03010] [BR:ko03011] | RP-S12, MRPS12, r | small subunit ribosomal protein S12 | FL-ls |
| K02952 | Genetic information proc | Ribosome | M00179 | Ribosome, archaea [PATH:map03010] [BR:ko03011] | RP-S13, rpsM | small subunit ribosomal protein S13 | FL-ls |
| K02956 | Genetic information proc | Ribosome | M00179 | Ribosome, archaea [PATH:map03010] [BR:ko03011] | RP-S15, MRPS15, r | small subunit ribosomal protein S15 | FL-ls |
| K02961 | Genetic information proc | Ribosome | M00179 | Ribosome, archaea [PATH:map03010] [BR:ko03011] | RP-S17, MRPS17, r | small subunit ribosomal protein S17 | FL-ls |
| K02962 | Genetic information proc | Ribosome | M00179 | Ribosome, archaea [PATH:map03010] [BR:ko03011] | RP-S17e, RPS17 | small subunit ribosomal protein S17e | FL-ls |
| K02965 | Genetic information proc | Ribosome | M00179 | Ribosome, archaea [PATH:map03010] [BR:ko03011] | RP-S19, rpsS | small subunit ribosomal protein S19 | FL-ls |
| K02967 | Genetic information proc | Ribosome | M00179 | Ribosome, archaea [PATH:map03010] [BR:ko03011] | RP-S2, MRPS2, rps | small subunit ribosomal protein S2 | FL-ls |
| K02975 | Genetic information proc | Ribosome | M00179 | Ribosome, archaea [PATH:map03010] [BR:ko03011] | RP-S25e, RPS25 | small subunit ribosomal protein S25e | FL-ls |
| K02976 | Genetic information proc | Ribosome | M00179 | Ribosome, archaea [PATH:map03010] [BR:ko03011] | RP-S26e, RPS26 | small subunit ribosomal protein S26e | FL-ls |
| K02977 | Genetic information proc | Ribosome | M00179 | Ribosome, archaea [PATH:map03010] [BR:ko03011] | RP-S27Ae, RPS27A | small subunit ribosomal protein S27Ae | FL-ls |
| K02978 | Genetic information proc | Ribosome | M00179 | Ribosome, archaea [PATH:map03010] [BR:ko03011] | RP-S27e, RPS27 | small subunit ribosomal protein S27e | FL-ls |
| K02979 | Genetic information proc | Ribosome | M00179 | Ribosome, archaea [PATH:map03010] [BR:ko03011] | RP-S28e, RPS28 | small subunit ribosomal protein S28e | FL-ls |
| K02982 | Genetic information proc | Ribosome | M00179 | Ribosome, archaea [PATH:map03010] [BR:ko03011] | RP-S3, rpsC | small subunit ribosomal protein S3 | FL-ls |
| K02983 | Genetic information proc | Ribosome | M00179 | Ribosome, archaea [PATH:map03010] [BR:ko03011] | RP-S30e, RPS30 | small subunit ribosomal protein S30e | FL-ls |
| K02984 | Genetic information proc | Ribosome | M00179 | Ribosome, archaea [PATH:map03010] [BR:ko03011] | RP-S3Ae, RPS3A | small subunit ribosomal protein S3Ae | FL-ls |
| K02986 | Genetic information proc | Ribosome | M00179 | Ribosome, archaea [PATH:map03010] [BR:ko03011] | RP-S4, rpsD | small subunit ribosomal protein S4 | FL-ls |
| K02987 | Genetic information proc | Ribosome | M00179 | Ribosome, archaea [PATH:map03010] [BR:ko03011] | RP-S4e, RPS4 | small subunit ribosomal protein S4e | FL-ls |
| K02988 | Genetic information proc | Ribosome | M00179 | Ribosome, archaea [PATH:map03010] [BR:ko03011] | RP-S5, MRPS5, rps | small subunit ribosomal protein S5 | FL-ls |
| K02992 | Genetic information proc | Ribosome | M00179 | Ribosome, archaea [PATH:map03010] [BR:ko03011] | RP-S7, MRPS7, rps | small subunit ribosomal protein S7 | FL-ls |
| K02994 | Genetic information proc | Ribosome | M00179 | Ribosome, archaea [PATH:map03010] [BR:ko03011] | RP-S8, rpsH | small subunit ribosomal protein S8 | FL-ls |
| K02995 | Genetic information proc | Ribosome | M00179 | Ribosome, archaea [PATH:map03010] [BR:ko03011] | RP-S8e, RPS8 | small subunit ribosomal protein S8e | FL-ls |
| K02996 | Genetic information proc | Ribosome | M00179 | Ribosome, archaea [PATH:map03010] [BR:ko03011] | RP-S9, MRPS9, rps | small subunit ribosomal protein S9 | FL-ls |
| K02879 | Genetic information proc | Ribosome | M00178 | Ribosome, bacteria [PATH:map03010] [BR:ko03011] | RP-L17, MRPL17, r | large subunit ribosomal protein L17 | PA-ls |
| K02884 | Genetic information proc | Ribosome | M00178 | Ribosome, bacteria [PATH:map03010] [BR:ko03011] | RP-L19, MRPL19, r | large subunit ribosomal protein L19 | PA-ls |
| K02887 | Genetic information proc | Ribosome | M00178 | Ribosome, bacteria [PATH:map03010] [BR:ko03011] | RP-L20, MRPL20, r | large subunit ribosomal protein L20 | PA-ls |
| K02888 | Genetic information proc | Ribosome | M00178 | Ribosome, bacteria [PATH:map03010] [BR:ko03011] | RP-L21, MRPL21, r | large subunit ribosomal protein L21 | PA-ls |
| K02897 | Genetic information proc | Ribosome | M00178 | Ribosome, bacteria [PATH:map03010] [BR:ko03011] | RP-L25, rplY | large subunit ribosomal protein L25 | PA-ls |

| | | | | | | | |
|---|---|---|---|---|---|---|---|
| K02899 | Genetic information proc | Ribosome | M00178 | Ribosome, bacteria [PATH:map03010] [BR:ko03011] | RP-L27, MRPL27, rp | large subunit ribosomal protein L27 | PA-ls |
| K02902 | Genetic information proc | Ribosome | M00178 | Ribosome, bacteria [PATH:map03010] [BR:ko03011] | RP-L28, MRPL28, rp | large subunit ribosomal protein L28 | PA-ls |
| K02909 | Genetic information proc | Ribosome | M00178 | Ribosome, bacteria [PATH:map03010] [BR:ko03011] | RP-L31, rpmE | large subunit ribosomal protein L31 | PA-ls |
| K02911 | Genetic information proc | Ribosome | M00178 | Ribosome, bacteria [PATH:map03010] [BR:ko03011] | RP-L32, MRPL32, rp | large subunit ribosomal protein L32 | PA-ls |
| K02913 | Genetic information proc | Ribosome | M00178 | Ribosome, bacteria [PATH:map03010] [BR:ko03011] | RP-L33, MRPL33, rp | large subunit ribosomal protein L33 | PA-ls |
| K02914 | Genetic information proc | Ribosome | M00178 | Ribosome, bacteria [PATH:map03010] [BR:ko03011] | RP-L34, MRPL34, rp | large subunit ribosomal protein L34 | PA-ls |
| K02916 | Genetic information proc | Ribosome | M00178 | Ribosome, bacteria [PATH:map03010] [BR:ko03011] | RP-L35, MRPL35, rp | large subunit ribosomal protein L35 | PA-ls |
| K02919 | Genetic information proc | Ribosome | M00178 | Ribosome, bacteria [PATH:map03010] [BR:ko03011] | RP-L36, MRPL36, rp | large subunit ribosomal protein L36 | PA-ls |
| K02926 | Genetic information proc | Ribosome | M00178 | Ribosome, bacteria [PATH:map03010] [BR:ko03011] | RP-L4, MRPL4, rplD | large subunit ribosomal protein L4 | PA-ls |
| K02945 | Genetic information proc | Ribosome | M00178 | Ribosome, bacteria [PATH:map03010] [BR:ko03011] | RP-S1, rpsA | small subunit ribosomal protein S1 | PA-ls |
| K02959 | Genetic information proc | Ribosome | M00178 | Ribosome, bacteria [PATH:map03010] [BR:ko03011] | RP-S16, MRPS16, r | small subunit ribosomal protein S16 | PA-ls |
| K02963 | Genetic information proc | Ribosome | M00178 | Ribosome, bacteria [PATH:map03010] [BR:ko03011] | RP-S18, MRPS18, r | small subunit ribosomal protein S18 | PA-ls |
| K02968 | Genetic information proc | Ribosome | M00178 | Ribosome, bacteria [PATH:map03010] [BR:ko03011] | RP-S20, rpsT | small subunit ribosomal protein S20 | PA-ls |
| K02990 | Genetic information proc | Ribosome | M00178 | Ribosome, bacteria [PATH:map03010] [BR:ko03011] | RP-S6, MRPS6, rps | small subunit ribosomal protein S6 | PA-ls |
| K02891 | Genetic information proc | Ribosome | M00177 | Ribosome, eukaryotes [PATH:map03010] [BR:ko03011] | RP-L22e, RPL22 | large subunit ribosomal protein L22e | shared |
| K02893 | Genetic information proc | Ribosome | M00177 | Ribosome, eukaryotes [PATH:map03010] [BR:ko03011] | RP-L23Ae, RPL23A | large subunit ribosomal protein L23Ae | shared |
| K02908 | Genetic information proc | Ribosome | M00177 | Ribosome, eukaryotes [PATH:map03010] [BR:ko03011] | RP-L30e, RPL30 | large subunit ribosomal protein L30e | shared |
| K02930 | Genetic information proc | Ribosome | M00177 | Ribosome, eukaryotes [PATH:map03010] [BR:ko03011] | RP-L4e, RPL4 | large subunit ribosomal protein L4e | shared |
| K02966 | Genetic information proc | Ribosome | M00177 | Ribosome, eukaryotes [PATH:map03010] [BR:ko03011] | RP-S19e, RPS19 | small subunit ribosomal protein S19e | shared |
| K02974 | Genetic information proc | Ribosome | M00177 | Ribosome, eukaryotes [PATH:map03010] [BR:ko03011] | RP-S24e, RPS24 | small subunit ribosomal protein S24e | shared |
| K02997 | Genetic information proc | Ribosome | M00177 | Ribosome, eukaryotes [PATH:map03010] [BR:ko03011] | RP-S9e, RPS9 | small subunit ribosomal protein S9e | shared |
| K02878 | Genetic information proc | Ribosome | M00178 | Ribosome, bacteria [PATH:map03010] [BR:ko03011] | RP-L16, MRPL16, rp | large subunit ribosomal protein L16 | shared |
| K02935 | Genetic information proc | Ribosome | M00178 | Ribosome, bacteria [PATH:map03010] [BR:ko03011] | RP-L7, MRPL12, rpl | large subunit ribosomal protein L7/L12 | shared |
| K02939 | Genetic information proc | Ribosome | M00178 | Ribosome, bacteria [PATH:map03010] [BR:ko03011] | RP-L9, MRPL9, rplI | large subunit ribosomal protein L9 | shared |
| K02954 | Genetic information proc | Ribosome | M00178 | Ribosome, bacteria [PATH:map03010] [BR:ko03011] | RP-S14, MRPS14, r | small subunit ribosomal protein S14 | shared |
| K02970 | Genetic information proc | Ribosome | M00178 | Ribosome, bacteria [PATH:map03010] [BR:ko03011] | RP-S21, MRPS21, r | small subunit ribosomal protein S21 | shared |
| K02908 | Genetic information proc | Ribosome | M00179 | Ribosome, archaea [PATH:map03010] [BR:ko03011] | RP-L30e, RPL30 | large subunit ribosomal protein L30e | shared |
| K02930 | Genetic information proc | Ribosome | M00179 | Ribosome, archaea [PATH:map03010] [BR:ko03011] | RP-L4e, RPL4 | large subunit ribosomal protein L4e | shared |
| K02944 | Genetic information proc | Ribosome | M00179 | Ribosome, archaea [PATH:map03010] [BR:ko03011] | RP-LX, rplX | large subunit ribosomal protein LX | shared |
| K02954 | Genetic information proc | Ribosome | M00179 | Ribosome, archaea [PATH:map03010] [BR:ko03011] | RP-S14, MRPS14, r | small subunit ribosomal protein S14 | shared |
| K02966 | Genetic information proc | Ribosome | M00179 | Ribosome, archaea [PATH:map03010] [BR:ko03011] | RP-S19e, RPS19 | small subunit ribosomal protein S19e | shared |
| K02974 | Genetic information proc | Ribosome | M00179 | Ribosome, archaea [PATH:map03010] [BR:ko03011] | RP-S24e, RPS24 | small subunit ribosomal protein S24e | shared |
| K07573 | Genetic information proc | RNA processing | M00390 | Exosome, archaea [PATH:map03018] | CSL4, EXOSC1 | exosome complex component CSL4 | FL-ls |
| K03679 | Genetic information proc | RNA processing | M00390 | Exosome, archaea [PATH:map03018] | RRP4, EXOSC2 | exosome complex component RRP4 | FL-ls |
| K11600 | Genetic information proc | RNA processing | M00390 | Exosome, archaea [PATH:map03018] | RRP41, EXOSC4, S | exosome complex component RRP41 | FL-ls |
| K12589 | Genetic information proc | RNA processing | M00390 | Exosome, archaea [PATH:map03018] | RRP42, EXOSC7 | exosome complex component RRP42 | FL-ls |
| K07573 | Genetic information proc | RNA processing | M00391 | Exosome, eukaryotes [PATH:map03018] | CSL4, EXOSC1 | exosome complex component CSL4 | FL-ls |
| K03679 | Genetic information proc | RNA processing | M00391 | Exosome, eukaryotes [PATH:map03018] | RRP4, EXOSC2 | exosome complex component RRP4 | FL-ls |
| K11600 | Genetic information proc | RNA processing | M00391 | Exosome, eukaryotes [PATH:map03018] | RRP41, EXOSC4, S | exosome complex component RRP41 | FL-ls |
| K12589 | Genetic information proc | RNA processing | M00391 | Exosome, eukaryotes [PATH:map03018] | RRP42, EXOSC7 | exosome complex component RRP42 | FL-ls |
| K11131 | Genetic information proc | RNA processing | M00425 | H/ACA ribonucleoprotein complex [PATH:map03008] [BR | DKC1, NOLA4, CBF | H/ACA ribonucleoprotein complex subunit 4 [EC:5.4.99.- | FL-ls |
| K08300 | Genetic information proc | RNA processing | M00394 | RNA degradosome [PATH:map03018] | rne | ribonuclease E [EC:3.1.26.12] | PA-ls |
| K12599 | Genetic information proc | RNA processing | M00392 | Ski complex [PATH:map03018] | SKI2, SKIV2L | antiviral helicase SKI2 [EC:3.6.4.-] | shared |
| K12600 | Genetic information proc | RNA processing | M00392 | Ski complex [PATH:map03018] | SKI3, TTC37 | superkiller protein 3 | shared |
| K03514 | Genetic information proc | RNA processing | M00393 | TRAMP complex [PATH:map03018] | PAPD5_7, TRF4 | non-canonical poly(A) RNA polymerase PAPD5/7 [EC:2. | shared |
| K12597 | Genetic information proc | RNA processing | M00393 | TRAMP complex [PATH:map03018] | AIR1_2 | protein AIR1/2 | shared |
| K03732 | Genetic information proc | RNA processing | M00394 | RNA degradosome [PATH:map03018] | rhlB | ATP-dependent RNA helicase RhlB [EC:3.6.4.13] | shared |
| K11130 | Genetic information proc | RNA processing | M00425 | H/ACA ribonucleoprotein complex [PATH:map03008] [BR | NOP10, NOLA3 | H/ACA ribonucleoprotein complex subunit 3 | shared |
| K12172 | Genetic information proc | RNA processing | M00427 | Nuclear pore complex [PATH:map03013] | RANBP2, NUP358 | E3 SUMO-protein ligase RanBP2 [EC:2.3.2.-] | shared |
| K14306 | Genetic information proc | RNA processing | M00427 | Nuclear pore complex [PATH:map03013] | NUP62, NSP1 | nuclear pore complex protein Nup62 | shared |
| K14319 | Genetic information proc | RNA processing | M00427 | Nuclear pore complex [PATH:map03013] | RANGAP1 | Ran GTPase-activating protein 1 | shared |
| K12160 | Genetic information proc | RNA processing | M00427 | Nuclear pore complex [PATH:map03013] | SUMO, SMT3 | small ubiquitin-related modifier | shared |
| K12854 | Genetic information proc | Spliceosome | M00354 | Spliceosome, U4/U6.U5 tri-snRNP [PATH:map03040] [BI | SNRNP200, BRR2 | pre-mRNA-splicing helicase BRR2 [EC:3.6.4.13] | PA-ls |
| K12854 | Genetic information proc | Spliceosome | M00355 | Spliceosome, 35S U5-snRNP [PATH:map03040] [BR:ko0 | SNRNP200, BRR2 | pre-mRNA-splicing helicase BRR2 [EC:3.6.4.13] | PA-ls |
| K11093 | Genetic information proc | Spliceosome | M00351 | Spliceosome, U1-snRNP [PATH:map03040] [BR:ko03041 | SNRP70 | U1 small nuclear ribonucleoprotein 70kDa | shared |
| K12825 | Genetic information proc | Spliceosome | M00352 | Spliceosome, U2-snRNP [PATH:map03040] [BR:ko03041 | SF3A1, SAP114 | splicing factor 3A subunit 1 | shared |
| K12860 | Genetic information proc | Spliceosome | M00353 | Spliceosome, Prp19/CDC5L complex [PATH:map03040] | CDC5L, CDC5, CEF | pre-mRNA-splicing factor CDC5/CEF1 | shared |
| K12863 | Genetic information proc | Spliceosome | M00353 | Spliceosome, Prp19/CDC5L complex [PATH:map03040] | CWC15 | protein CWC15 | shared |
| K12858 | Genetic information proc | Spliceosome | M00354 | Spliceosome, U4/U6.U5 tri-snRNP [PATH:map03040] [BI | DDX23, PRP28 | ATP-dependent RNA helicase DDX23/PRP28 [EC:3.6.4.1 | shared |
| K12846 | Genetic information proc | Spliceosome | M00354 | Spliceosome, U4/U6.U5 tri-snRNP [PATH:map03040] [BI | SNRNP27 | U4/U6.U5 tri-snRNP-associated protein 3 | shared |

| | | | | | | | |
|---|---|---|---|---|---|---|---|
| K12869 | Genetic information proce | Spliceosome | M00355 | Spliceosome, 35S U5-snRNP [PATH:map03040] [BR:ko0 | CRN, CRNKL1, CLF | crooked neck | shared |
| K12860 | Genetic information proce | Spliceosome | M00355 | Spliceosome, 35S U5-snRNP [PATH:map03040] [BR:ko0 | CDC5L, CDC5, CEF | pre-mRNA-splicing factor CDC5/CEF1 | shared |
| K12863 | Genetic information proce | Spliceosome | M00355 | Spliceosome, 35S U5-snRNP [PATH:map03040] [BR:ko0 | CWC15 | protein CWC15 | shared |
| K01001 | Glycan metabolism | Glycan biosynthesis | M00055 | N-glycan precursor biosynthesis [PATH:map00510 map0 | ALG7 | UDP-N-acetylglucosamine--dolichyl-phosphate N-acetylg | FL-ls |
| K00737 | Glycan metabolism | Glycan biosynthesis | M00075 | N-glycan biosynthesis, complex type [PATH:map00510 | MGAT3 | beta-1,4-mannosyl-glycoprotein beta-1,4-N-acetylglucos | PA-ls |
| K00729 | Glycan metabolism | Glycan biosynthesis | M00055 | N-glycan precursor biosynthesis [PATH:map00510 map0 | ALG5 | dolichyl-phosphate beta-glucosyltransferase [EC:2.4.1.1 | shared |
| K03857 | Glycan metabolism | Glycan biosynthesis | M00065 | GPI-anchor biosynthesis, core oligosaccharide [PATH:m | PIGA, GPI3 | phosphatidylinositol glycan, class A [EC:2.4.1.198] | shared |
| K05286 | Glycan metabolism | Glycan biosynthesis | M00065 | GPI-anchor biosynthesis, core oligosaccharide [PATH:m | PIGB | phosphatidylinositol glycan, class B [EC:2.4.1.-] | shared |
| K01197 | Glycan metabolism | Glycosaminoglycan metabolism | M00076 | Dermatan sulfate degradation [PATH:map00531 map011 | hya | hyaluronoglucosaminidase [EC:3.2.1.35] | shared |
| K01197 | Glycan metabolism | Glycosaminoglycan metabolism | M00077 | Chondroitin sulfate degradation [PATH:map00531 map01 | hya | hyaluronoglucosaminidase [EC:3.2.1.35] | shared |
| K03272 | Glycan metabolism | Lipopolysaccharide metabolism | M00064 | ADP-L-glycero-D-manno-heptose biosynthesis [PATH:ma | gmhC, hldE, waaE, | D-beta-D-heptose 7-phosphate kinase / D-beta-D-heptos | FL-ls |
| K02843 | Glycan metabolism | Lipopolysaccharide metabolism | M00080 | Lipopolysaccharide biosynthesis, inner core => outer co | waaF, rfaF | heptosyltransferase II [EC:2.4.-.-] | FL-ls |
| K02527 | Glycan metabolism | Lipopolysaccharide metabolism | M00060 | Lipopolysaccharide biosynthesis, KDO2-lipid A [PATH:m | kdtA, waaA | 3-deoxy-D-manno-octulosonic-acid transferase [EC:2.4. | shared |
| K02517 | Glycan metabolism | Lipopolysaccharide metabolism | M00060 | Lipopolysaccharide biosynthesis, KDO2-lipid A [PATH:m | lpxL, htrB | Kdo2-lipid IVA lauroyltransferase [EC:2.3.1.241] | shared |
| K00748 | Glycan metabolism | Lipopolysaccharide metabolism | M00060 | Lipopolysaccharide biosynthesis, KDO2-lipid A [PATH:m | lpxB | lipid-A-disaccharide synthase [EC:2.4.1.182] | shared |
| K00912 | Glycan metabolism | Lipopolysaccharide metabolism | M00060 | Lipopolysaccharide biosynthesis, KDO2-lipid A [PATH:m | lpxK | tetraacyldisaccharide 4'-kinase [EC:2.7.1.130] | shared |
| K03269 | Glycan metabolism | Lipopolysaccharide metabolism | M00060 | Lipopolysaccharide biosynthesis, KDO2-lipid A [PATH:m | lpxH | UDP-2,3-diacylglucosamine hydrolase [EC:3.6.1.54] | shared |
| K02536 | Glycan metabolism | Lipopolysaccharide metabolism | M00060 | Lipopolysaccharide biosynthesis, KDO2-lipid A [PATH:m | lpxD | UDP-3-O-[3-hydroxymyristoyl] glucosamine N-acyltransf | shared |
| K02535 | Glycan metabolism | Lipopolysaccharide metabolism | M00060 | Lipopolysaccharide biosynthesis, KDO2-lipid A [PATH:m | lpxC | UDP-3-O-[3-hydroxymyristoyl] N-acetylglucosamine dea | shared |
| K01627 | Glycan metabolism | Lipopolysaccharide metabolism | M00063 | CMP-KDO biosynthesis [PATH:map00540 map01100] | kdsA | 2-dehydro-3-deoxyphosphooctonate aldolase (KDO 8-P | shared |
| K03270 | Glycan metabolism | Lipopolysaccharide metabolism | M00063 | CMP-KDO biosynthesis [PATH:map00540 map01100] | kdsC | 3-deoxy-D-manno-octulosonate 8-phosphate phosphatas | shared |
| K00979 | Glycan metabolism | Lipopolysaccharide metabolism | M00063 | CMP-KDO biosynthesis [PATH:map00540 map01100] | kdsB | 3-deoxy-manno-octulosonate cytidylyltransferase (CMP- | shared |
| K06041 | Glycan metabolism | Lipopolysaccharide metabolism | M00063 | CMP-KDO biosynthesis [PATH:map00540 map01100] | kdsD, kpsF | arabinose-5-phosphate isomerase [EC:5.3.1.13] | shared |
| K03274 | Glycan metabolism | Lipopolysaccharide metabolism | M00064 | ADP-L-glycero-D-manno-heptose biosynthesis [PATH:ma | gmhD, rfaD | ADP-L-glycero-D-manno-heptose 6-epimerase [EC:5.1.3 | shared |
| K03273 | Glycan metabolism | Lipopolysaccharide metabolism | M00064 | ADP-L-glycero-D-manno-heptose biosynthesis [PATH:ma | gmhB | D-glycero-D-manno-heptose 1,7-bisphosphate phosphat | shared |
| K03271 | Glycan metabolism | Lipopolysaccharide metabolism | M00064 | ADP-L-glycero-D-manno-heptose biosynthesis [PATH:ma | gmhA, lpcA | D-sedoheptulose 7-phosphate isomerase [EC:5.3.1.28] | shared |
| K02527 | Glycan metabolism | Lipopolysaccharide metabolism | M00080 | Lipopolysaccharide biosynthesis, inner core => outer co | kdtA, waaA | 3-deoxy-D-manno-octulosonic-acid transferase [EC:2.4. | shared |
| K02848 | Glycan metabolism | Lipopolysaccharide metabolism | M00080 | Lipopolysaccharide biosynthesis, inner core => outer co | waaP, rfaP | heptose I phosphotransferase [EC:2.7.1.-] | shared |
| K02841 | Glycan metabolism | Lipopolysaccharide metabolism | M00080 | Lipopolysaccharide biosynthesis, inner core => outer co | waaC, rfaC | heptosyltransferase I [EC:2.4.-.-] | shared |
| K02849 | Glycan metabolism | Lipopolysaccharide metabolism | M00080 | Lipopolysaccharide biosynthesis, inner core => outer co | waaQ, rfaQ | heptosyltransferase III [EC:2.4.-.-] | shared |
| K02844 | Glycan metabolism | Lipopolysaccharide metabolism | M00080 | Lipopolysaccharide biosynthesis, inner core => outer co | waaG, rfaG | UDP-glucose:(heptosyl)LPS alpha-1,3-glucosyltransfera | shared |
| K02371 | Lipid metabolism | Fatty acid biosynthesis and degrada | M00083 | Fatty acid biosynthesis, elongation [PATH:map00061 ma | fabK | enoyl-[acyl-carrier protein] reductase II [EC:1.3.1.9] | FL-ls |
| K00645 | Lipid metabolism | Fatty acid biosynthesis and degrada | M00082 | Fatty acid biosynthesis, initiation [PATH:map00061 map | fabD | [acyl-carrier-protein] S-malonyltransferase [EC:2.3.1.39] | PA-ls |
| K00648 | Lipid metabolism | Fatty acid biosynthesis and degrada | M00082 | Fatty acid biosynthesis, initiation [PATH:map00061 map | fabH | 3-oxoacyl-[acyl-carrier-protein] synthase III [EC:2.3.1.1 | PA-ls |
| K00667 | Lipid metabolism | Fatty acid biosynthesis and degrada | M00082 | Fatty acid biosynthesis, initiation [PATH:map00061 map | FAS2 | fatty acid synthase subunit alpha, fungi type [EC:2.3.1. | PA-ls |
| K01716 | Lipid metabolism | Fatty acid biosynthesis and degrada | M00083 | Fatty acid biosynthesis, elongation [PATH:map00061 ma | fabA | 3-hydroxyacyl-[acyl-carrier protein] dehydratase / trans- | PA-ls |
| K00667 | Lipid metabolism | Fatty acid biosynthesis and degrada | M00083 | Fatty acid biosynthesis, elongation [PATH:map00061 ma | FAS2 | fatty acid synthase subunit alpha, fungi type [EC:2.3.1. | PA-ls |
| K10780 | Lipid metabolism | Fatty acid biosynthesis and degrada | M00083 | Fatty acid biosynthesis, elongation [PATH:map00061 ma | fabL | enoyl-[acyl-carrier protein] reductase III [EC:1.3.1.104] | shared |
| K06445 | Lipid metabolism | Fatty acid biosynthesis and degrada | M00087 | beta-Oxidation [PATH:map00071 map01212 map01100] | fadE | acyl-CoA dehydrogenase [EC:1.3.99.-] | shared |
| K09479 | Lipid metabolism | Fatty acid biosynthesis and degrada | M00087 | beta-Oxidation [PATH:map00071 map01212 map01100] | ACADVL | very long chain acyl-CoA dehydrogenase [EC:1.3.8.9] | shared |
| K00635 | Lipid metabolism | Lipid metabolism | M00089 | Triacylglycerol biosynthesis [PATH:map00561 map0110 | tgs, wax-dgat | diacylglycerol O-acyltransferase / wax synthase [EC:2.3 | PA-ls |
| K00570 | Lipid metabolism | Lipid metabolism | M00091 | Phosphatidylcholine (PC) biosynthesis, PE => PC [PATH | pmtA | phosphatidylethanolamine/phosphatidyl-N-methylethanol | PA-ls |
| K01613 | Lipid metabolism | Lipid metabolism | M00093 | Phosphatidylethanolamine (PE) biosynthesis, PA => PS | psd, PISD | phosphatidylserine decarboxylase [EC:4.1.1.65] | PA-ls |
| K01046 | Lipid metabolism | Lipid metabolism | M00098 | Acylglycerol degradation [PATH:map00561 map01100] | lip, TGL2 | triacylglycerol lipase [EC:3.1.1.3] | PA-ls |
| K00866 | Lipid metabolism | Lipid metabolism | M00090 | Phosphatidylcholine (PC) biosynthesis, choline => PC [F | CKI1 | choline kinase [EC:2.7.1.32] | shared |
| K00551 | Lipid metabolism | Lipid metabolism | M00091 | Phosphatidylcholine (PC) biosynthesis, PE => PC [PATH | PEMT | phosphatidylethanolamine/phosphatidyl-N-methylethanol | shared |
| K04708 | Lipid metabolism | Lipid metabolism | M00094 | Ceramide biosynthesis [PATH:map00600 map01100] | E1.1.1.102 | 3-dehydrosphinganine reductase [EC:1.1.1.102] | shared |
| K04708 | Lipid metabolism | Lipid metabolism | M00099 | Sphingosine biosynthesis [PATH:map00600 map01100] | E1.1.1.102 | 3-dehydrosphinganine reductase [EC:1.1.1.102] | shared |
| K10526 | Lipid metabolism | Lipid metabolism | M00113 | Jasmonic acid biosynthesis [PATH:map00592 map0110 | OPCL1 | OPC-8:0 CoA ligase 1 [EC:6.2.1.-] | shared |
| K05917 | Lipid metabolism | Sterol biosynthesis | M00101 | Cholesterol biosynthesis, squalene 2,3-epoxide => chol | CYP51 | sterol 14alpha-demethylase [EC:1.14.14.154] | PA-ls |
| K09828 | Lipid metabolism | Sterol biosynthesis | M00101 | Cholesterol biosynthesis, squalene 2,3-epoxide => chol | DHCR24, DWF1 | Delta24-sterol reductase [EC:1.3.1.72 1.3.1.-] | shared |
| K00227 | Lipid metabolism | Sterol biosynthesis | M00101 | Cholesterol biosynthesis, squalene 2,3-epoxide => chol | SC5DL, ERG3 | Delta7-sterol 5-desaturase [EC:1.14.19.20] | shared |
| K07750 | Lipid metabolism | Sterol biosynthesis | M00101 | Cholesterol biosynthesis, squalene 2,3-epoxide => chol | MESO1, ERG25 | methylsterol monooxygenase [EC:1.14.18.9] | shared |
| K07748 | Lipid metabolism | Sterol biosynthesis | M00101 | Cholesterol biosynthesis, squalene 2,3-epoxide => chol | E1.1.1.170, NSDHL | sterol-4alpha-carboxylate 3-dehydrogenase (decarboxyl | shared |
| K00223 | Lipid metabolism | Sterol biosynthesis | M00102 | Ergocalciferol biosynthesis [PATH:map00100 map01100 | ERG4 | Delta24(24(1))-sterol reductase [EC:1.3.1.71] | shared |
| K00227 | Lipid metabolism | Sterol biosynthesis | M00102 | Ergocalciferol biosynthesis [PATH:map00100 map01100 | SC5DL, ERG3 | Delta7-sterol 5-desaturase [EC:1.14.19.20] | shared |
| K00559 | Lipid metabolism | Sterol biosynthesis | M00102 | Ergocalciferol biosynthesis [PATH:map00100 map01100 | E2.1.1.41, SMT1, E | sterol 24-C-methyltransferase [EC:2.1.1.41] | shared |
| K01872 | Metabolism | Aminoacyl tRNA | M00359 | Aminoacyl-tRNA biosynthesis, eukaryotes [PATH:map00 | AARS, alaS | alanyl-tRNA synthetase [EC:6.1.1.7] | FL-ls |
| K01887 | Metabolism | Aminoacyl tRNA | M00359 | Aminoacyl-tRNA biosynthesis, eukaryotes [PATH:map00 | RARS, argS | arginyl-tRNA synthetase [EC:6.1.1.19] | FL-ls |
| K01876 | Metabolism | Aminoacyl tRNA | M00359 | Aminoacyl-tRNA biosynthesis, eukaryotes [PATH:map00 | aspS | aspartyl-tRNA synthetase [EC:6.1.1.12] | FL-ls |

| | | | | | | | |
|---|---|---|---|---|---|---|---|
| K01883 | Metabolism | Aminoacyl tRNA | M00359 | Aminoacyl-tRNA biosynthesis, eukaryotes [PATH:map00 | CARS, cysS | cysteinyl-tRNA synthetase [EC:6.1.1.16] | FL-ls |
| K01886 | Metabolism | Aminoacyl tRNA | M00359 | Aminoacyl-tRNA biosynthesis, eukaryotes [PATH:map00 | QARS, glnS | glutaminyl-tRNA synthetase [EC:6.1.1.18] | FL-ls |
| K01880 | Metabolism | Aminoacyl tRNA | M00359 | Aminoacyl-tRNA biosynthesis, eukaryotes [PATH:map00 | GARS, glyS1 | glycyl-tRNA synthetase [EC:6.1.1.14] | FL-ls |
| K01892 | Metabolism | Aminoacyl tRNA | M00359 | Aminoacyl-tRNA biosynthesis, eukaryotes [PATH:map00 | HARS, hisS | histidyl-tRNA synthetase [EC:6.1.1.21] | FL-ls |
| K01870 | Metabolism | Aminoacyl tRNA | M00359 | Aminoacyl-tRNA biosynthesis, eukaryotes [PATH:map00 | IARS, ileS | isoleucyl-tRNA synthetase [EC:6.1.1.5] | FL-ls |
| K01869 | Metabolism | Aminoacyl tRNA | M00359 | Aminoacyl-tRNA biosynthesis, eukaryotes [PATH:map00 | LARS, leuS | leucyl-tRNA synthetase [EC:6.1.1.4] | FL-ls |
| K01889 | Metabolism | Aminoacyl tRNA | M00359 | Aminoacyl-tRNA biosynthesis, eukaryotes [PATH:map00 | FARSA, pheS | phenylalanyl-tRNA synthetase alpha chain [EC:6.1.1.20] | FL-ls |
| K01890 | Metabolism | Aminoacyl tRNA | M00359 | Aminoacyl-tRNA biosynthesis, eukaryotes [PATH:map00 | FARSB, pheT | phenylalanyl-tRNA synthetase beta chain [EC:6.1.1.20] | FL-ls |
| K01875 | Metabolism | Aminoacyl tRNA | M00359 | Aminoacyl-tRNA biosynthesis, eukaryotes [PATH:map00 | SARS, serS | seryl-tRNA synthetase [EC:6.1.1.11] | FL-ls |
| K01868 | Metabolism | Aminoacyl tRNA | M00359 | Aminoacyl-tRNA biosynthesis, eukaryotes [PATH:map00 | TARS, thrS | threonyl-tRNA synthetase [EC:6.1.1.3] | FL-ls |
| K01867 | Metabolism | Aminoacyl tRNA | M00359 | Aminoacyl-tRNA biosynthesis, eukaryotes [PATH:map00 | WARS, trpS | tryptophanyl-tRNA synthetase [EC:6.1.1.2] | FL-ls |
| K01866 | Metabolism | Aminoacyl tRNA | M00359 | Aminoacyl-tRNA biosynthesis, eukaryotes [PATH:map00 | YARS, tyrS | tyrosyl-tRNA synthetase [EC:6.1.1.1] | FL-ls |
| K01873 | Metabolism | Aminoacyl tRNA | M00359 | Aminoacyl-tRNA biosynthesis, eukaryotes [PATH:map00 | VARS, valS | valyl-tRNA synthetase [EC:6.1.1.9] | FL-ls |
| K01872 | Metabolism | Aminoacyl tRNA | M00360 | Aminoacyl-tRNA biosynthesis, prokaryotes [PATH:map0 | AARS, alaS | alanyl-tRNA synthetase [EC:6.1.1.7] | FL-ls |
| K01887 | Metabolism | Aminoacyl tRNA | M00360 | Aminoacyl-tRNA biosynthesis, prokaryotes [PATH:map0 | RARS, argS | arginyl-tRNA synthetase [EC:6.1.1.19] | FL-ls |
| K01876 | Metabolism | Aminoacyl tRNA | M00360 | Aminoacyl-tRNA biosynthesis, prokaryotes [PATH:map0 | aspS | aspartyl-tRNA synthetase [EC:6.1.1.12] | FL-ls |
| K01883 | Metabolism | Aminoacyl tRNA | M00360 | Aminoacyl-tRNA biosynthesis, prokaryotes [PATH:map0 | CARS, cysS | cysteinyl-tRNA synthetase [EC:6.1.1.16] | FL-ls |
| K01886 | Metabolism | Aminoacyl tRNA | M00360 | Aminoacyl-tRNA biosynthesis, prokaryotes [PATH:map0 | QARS, glnS | glutaminyl-tRNA synthetase [EC:6.1.1.18] | FL-ls |
| K01880 | Metabolism | Aminoacyl tRNA | M00360 | Aminoacyl-tRNA biosynthesis, prokaryotes [PATH:map0 | GARS, glyS1 | glycyl-tRNA synthetase [EC:6.1.1.14] | FL-ls |
| K01892 | Metabolism | Aminoacyl tRNA | M00360 | Aminoacyl-tRNA biosynthesis, prokaryotes [PATH:map0 | HARS, hisS | histidyl-tRNA synthetase [EC:6.1.1.21] | FL-ls |
| K01870 | Metabolism | Aminoacyl tRNA | M00360 | Aminoacyl-tRNA biosynthesis, prokaryotes [PATH:map0 | IARS, ileS | isoleucyl-tRNA synthetase [EC:6.1.1.5] | FL-ls |
| K01869 | Metabolism | Aminoacyl tRNA | M00360 | Aminoacyl-tRNA biosynthesis, prokaryotes [PATH:map0 | LARS, leuS | leucyl-tRNA synthetase [EC:6.1.1.4] | FL-ls |
| K04566 | Metabolism | Aminoacyl tRNA | M00360 | Aminoacyl-tRNA biosynthesis, prokaryotes [PATH:map0 | lysK | lysyl-tRNA synthetase, class I [EC:6.1.1.6] | FL-ls |
| K09698 | Metabolism | Aminoacyl tRNA | M00360 | Aminoacyl-tRNA biosynthesis, prokaryotes [PATH:map0 | gltX | nondiscriminating glutamyl-tRNA synthetase [EC:6.1.1.2 | FL-ls |
| K01889 | Metabolism | Aminoacyl tRNA | M00360 | Aminoacyl-tRNA biosynthesis, prokaryotes [PATH:map0 | FARSA, pheS | phenylalanyl-tRNA synthetase alpha chain [EC:6.1.1.20] | FL-ls |
| K01890 | Metabolism | Aminoacyl tRNA | M00360 | Aminoacyl-tRNA biosynthesis, prokaryotes [PATH:map0 | FARSB, pheT | phenylalanyl-tRNA synthetase beta chain [EC:6.1.1.20] | FL-ls |
| K01875 | Metabolism | Aminoacyl tRNA | M00360 | Aminoacyl-tRNA biosynthesis, prokaryotes [PATH:map0 | SARS, serS | seryl-tRNA synthetase [EC:6.1.1.11] | FL-ls |
| K01868 | Metabolism | Aminoacyl tRNA | M00360 | Aminoacyl-tRNA biosynthesis, prokaryotes [PATH:map0 | TARS, thrS | threonyl-tRNA synthetase [EC:6.1.1.3] | FL-ls |
| K01867 | Metabolism | Aminoacyl tRNA | M00360 | Aminoacyl-tRNA biosynthesis, prokaryotes [PATH:map0 | WARS, trpS | tryptophanyl-tRNA synthetase [EC:6.1.1.2] | FL-ls |
| K01866 | Metabolism | Aminoacyl tRNA | M00360 | Aminoacyl-tRNA biosynthesis, prokaryotes [PATH:map0 | YARS, tyrS | tyrosyl-tRNA synthetase [EC:6.1.1.1] | FL-ls |
| K01873 | Metabolism | Aminoacyl tRNA | M00360 | Aminoacyl-tRNA biosynthesis, prokaryotes [PATH:map0 | VARS, valS | valyl-tRNA synthetase [EC:6.1.1.9] | FL-ls |
| K01878 | Metabolism | Aminoacyl tRNA | M00360 | Aminoacyl-tRNA biosynthesis, prokaryotes [PATH:map0 | glyQ | glycyl-tRNA synthetase alpha chain [EC:6.1.1.14] | PA-ls |
| K01879 | Metabolism | Aminoacyl tRNA | M00360 | Aminoacyl-tRNA biosynthesis, prokaryotes [PATH:map0 | glyS | glycyl-tRNA synthetase beta chain [EC:6.1.1.14] | PA-ls |
| K01893 | Metabolism | Aminoacyl tRNA | M00359 | Aminoacyl-tRNA biosynthesis, eukaryotes [PATH:map00 | NARS, asnS | asparaginyl-tRNA synthetase [EC:6.1.1.22] | shared |
| K04567 | Metabolism | Aminoacyl tRNA | M00359 | Aminoacyl-tRNA biosynthesis, eukaryotes [PATH:map00 | KARS, lysS | lysyl-tRNA synthetase, class II [EC:6.1.1.6] | shared |
| K01881 | Metabolism | Aminoacyl tRNA | M00359 | Aminoacyl-tRNA biosynthesis, eukaryotes [PATH:map00 | PARS, proS | prolyl-tRNA synthetase [EC:6.1.1.15] | shared |
| K01893 | Metabolism | Aminoacyl tRNA | M00360 | Aminoacyl-tRNA biosynthesis, prokaryotes [PATH:map0 | NARS, asnS | asparaginyl-tRNA synthetase [EC:6.1.1.22] | shared |
| K01884 | Metabolism | Aminoacyl tRNA | M00360 | Aminoacyl-tRNA biosynthesis, prokaryotes [PATH:map0 | cysS1 | cysteinyl-tRNA synthetase, unknown class [EC:6.1.1.16 | shared |
| K04567 | Metabolism | Aminoacyl tRNA | M00360 | Aminoacyl-tRNA biosynthesis, prokaryotes [PATH:map0 | KARS, lysS | lysyl-tRNA synthetase, class II [EC:6.1.1.6] | shared |
| K09759 | Metabolism | Aminoacyl tRNA | M00360 | Aminoacyl-tRNA biosynthesis, prokaryotes [PATH:map0 | aspC, aspS | nondiscriminating aspartyl-tRNA synthetase [EC:6.1.1.2 | shared |
| K01881 | Metabolism | Aminoacyl tRNA | M00360 | Aminoacyl-tRNA biosynthesis, prokaryotes [PATH:map0 | PARS, proS | prolyl-tRNA synthetase [EC:6.1.1.15] | shared |
| K00972 | Metabolism | Nucleotide sugar | M00361 | Nucleotide sugar biosynthesis, eukaryotes [PATH:map0 | UAP1 | UDP-N-acetylglucosamine/UDP-N-acetylgalactosamine d | FL-ls |
| K00972 | Metabolism | Nucleotide sugar | M00362 | Nucleotide sugar biosynthesis, prokaryotes [PATH:map0 | UAP1 | UDP-N-acetylglucosamine/UDP-N-acetylgalactosamine d | FL-ls |
| K04042 | Metabolism | Nucleotide sugar | M00362 | Nucleotide sugar biosynthesis, prokaryotes [PATH:map0 | glmU | bifunctional UDP-N-acetylglucosamine pyrophosphorylas | PA-ls |
| K01791 | Metabolism | Nucleotide sugar | M00362 | Nucleotide sugar biosynthesis, prokaryotes [PATH:map0 | wecB | UDP-N-acetylglucosamine 2-epimerase (non-hydrolysing | shared |
| K11528 | Metabolism | Nucleotide sugar | M00362 | Nucleotide sugar biosynthesis, prokaryotes [PATH:map0 | glmU | UDP-N-acetylglucosamine pyrophosphorylase [EC:2.7.7 | shared |
| K00767 | Metabolism of cofactors | Cofactor and vitamin metabolism | M00115 | NAD biosynthesis, aspartate => NAD [PATH:map00760 | nadC, QPRT | nicotinate-nucleotide pyrophosphorylase (carboxylating) | FL-ls |
| K02548 | Metabolism of cofactors | Cofactor and vitamin metabolism | M00116 | Menaquinone biosynthesis, chorismate => menaquinol [ | menA | 1,4-dihydroxy-2-naphthoate polyprenyltransferase [EC:2 | FL-ls |
| K03183 | Metabolism of cofactors | Cofactor and vitamin metabolism | M00116 | Menaquinone biosynthesis, chorismate => menaquinol [ | ubiE | demethylmenaquinone methyltransferase / 2-methoxy-6- | FL-ls |
| K03182 | Metabolism of cofactors | Cofactor and vitamin metabolism | M00117 | Ubiquinone biosynthesis, prokaryotes, chorismate => ub | ubiD | 4-hydroxy-3-polyprenylbenzoate decarboxylase [EC:4.1 | FL-ls |
| K03183 | Metabolism of cofactors | Cofactor and vitamin metabolism | M00117 | Ubiquinone biosynthesis, prokaryotes, chorismate => ub | ubiE | demethylmenaquinone methyltransferase / 2-methoxy-6- | FL-ls |
| K00077 | Metabolism of cofactors | Cofactor and vitamin metabolism | M00119 | Pantothenate biosynthesis, valine/L-aspartate => panto | panE, apbA | 2-dehydropantoate 2-reductase [EC:1.1.1.169] | FL-ls |
| K00606 | Metabolism of cofactors | Cofactor and vitamin metabolism | M00119 | Pantothenate biosynthesis, valine/L-aspartate => panto | panB | 3-methyl-2-oxobutanoate hydroxymethyltransferase [EC | FL-ls |
| K13038 | Metabolism of cofactors | Cofactor and vitamin metabolism | M00120 | Coenzyme A biosynthesis, pantothenate => CoA [PATH | coaBC, dfp | phosphopantothenoylcysteine decarboxylase / phospho | FL-ls |
| K01845 | Metabolism of cofactors | Cofactor and vitamin metabolism | M00121 | Heme biosynthesis, glutamate => heme [PATH:map0086 | hemL | glutamate-1-semialdehyde 2,1-aminomutase [EC:5.4.3.8 | FL-ls |
| K02492 | Metabolism of cofactors | Cofactor and vitamin metabolism | M00121 | Heme biosynthesis, glutamate => heme [PATH:map0086 | hemA | glutamyl-tRNA reductase [EC:1.2.1.70] | FL-ls |
| K01698 | Metabolism of cofactors | Cofactor and vitamin metabolism | M00121 | Heme biosynthesis, glutamate => heme [PATH:map0086 | hemB, ALAD | porphobilinogen synthase [EC:4.2.1.24] | FL-ls |
| K02233 | Metabolism of cofactors | Cofactor and vitamin metabolism | M00122 | Cobalamin biosynthesis, cobinamide => cobalamin [PATH | E2.7.8.26, cobS, co | adenosylcobinamide-GDP ribazoletransferase [EC:2.7.8 | FL-ls |
| K02227 | Metabolism of cofactors | Cofactor and vitamin metabolism | M00122 | Cobalamin biosynthesis, cobinamide => cobalamin [PAT | cbiB, cobD | adenosylcobinamide-phosphate synthase [EC:6.3.1.10] | FL-ls |
| K02232 | Metabolism of cofactors | Cofactor and vitamin metabolism | M00122 | Cobalamin biosynthesis, cobinamide => cobalamin [PAT | cobQ, cbiP | adenosylcobyric acid synthase [EC:6.3.5.10] | FL-ls |

| | | | | | | | |
|---|---|---|---|---|---|---|---|
| K00798 | Metabolism of cofactors | Cofactor and vitamin metabolism | M00122 | Cobalamin biosynthesis, cobinamide => cobalamin [PATI | MMAB, pduO | cob(I)alamin adenosyltransferase [EC:2.5.1.17] | FL-ls |
| K00833 | Metabolism of cofactors | Cofactor and vitamin metabolism | M00123 | Biotin biosynthesis, pimeloyl-ACP/CoA => biotin [PATH:r | bioA | adenosylmethionine---8-amino-7-oxononanoate aminotra | FL-ls |
| K01012 | Metabolism of cofactors | Cofactor and vitamin metabolism | M00123 | Biotin biosynthesis, pimeloyl-ACP/CoA => biotin [PATH:r | bioB | biotin synthase [EC:2.8.1.6] | FL-ls |
| K01935 | Metabolism of cofactors | Cofactor and vitamin metabolism | M00123 | Biotin biosynthesis, pimeloyl-ACP/CoA => biotin [PATH:r | bioD | dethiobiotin synthetase [EC:6.3.3.3] | FL-ls |
| K02858 | Metabolism of cofactors | Cofactor and vitamin metabolism | M00125 | Riboflavin biosynthesis, GTP => riboflavin/FMN/FAD [PA | ribB, RIB3 | 3,4-dihydroxy 2-butanone 4-phosphate synthase [EC:4. | FL-ls |
| K00794 | Metabolism of cofactors | Cofactor and vitamin metabolism | M00125 | Riboflavin biosynthesis, GTP => riboflavin/FMN/FAD [PA | ribH, RIB4 | 6,7-dimethyl-8-ribityllumazine synthase [EC:2.5.1.78] | FL-ls |
| K00793 | Metabolism of cofactors | Cofactor and vitamin metabolism | M00125 | Riboflavin biosynthesis, GTP => riboflavin/FMN/FAD [PA | ribE, RIB5 | riboflavin synthase [EC:2.5.1.9] | FL-ls |
| K00796 | Metabolism of cofactors | Cofactor and vitamin metabolism | M00126 | Tetrahydrofolate biosynthesis, GTP => THF [PATH:map0 | folP | dihydropteroate synthase [EC:2.5.1.15] | FL-ls |
| K01495 | Metabolism of cofactors | Cofactor and vitamin metabolism | M00126 | Tetrahydrofolate biosynthesis, GTP => THF [PATH:map0 | GCH1, folE | GTP cyclohydrolase IA [EC:3.5.4.16] | FL-ls |
| K03147 | Metabolism of cofactors | Cofactor and vitamin metabolism | M00127 | Thiamine biosynthesis, AIR => thiamine-P/thiamine-2P [ | thiC | phosphomethylpyrimidine synthase [EC:4.1.99.17] | FL-ls |
| K00946 | Metabolism of cofactors | Cofactor and vitamin metabolism | M00127 | Thiamine biosynthesis, AIR => thiamine-P/thiamine-2P [ | thiL | thiamine-monophosphate kinase [EC:2.7.4.16] | FL-ls |
| K00833 | Metabolism of cofactors | Cofactor and vitamin metabolism | M00573 | Biotin biosynthesis, BioI pathway, long-chain-acyl-ACP | bioA | adenosylmethionine---8-amino-7-oxononanoate aminotra | FL-ls |
| K01012 | Metabolism of cofactors | Cofactor and vitamin metabolism | M00573 | Biotin biosynthesis, BioI pathway, long-chain-acyl-ACP | bioB | biotin synthase [EC:2.8.1.6] | FL-ls |
| K01935 | Metabolism of cofactors | Cofactor and vitamin metabolism | M00573 | Biotin biosynthesis, BioI pathway, long-chain-acyl-ACP | bioD | dethiobiotin synthetase [EC:6.3.3.3] | FL-ls |
| K00833 | Metabolism of cofactors | Cofactor and vitamin metabolism | M00577 | Biotin biosynthesis, BioW pathway, pimelate => pimeloyl | bioA | adenosylmethionine---8-amino-7-oxononanoate aminotra | FL-ls |
| K01012 | Metabolism of cofactors | Cofactor and vitamin metabolism | M00577 | Biotin biosynthesis, BioW pathway, pimelate => pimeloyl | bioB | biotin synthase [EC:2.8.1.6] | FL-ls |
| K01935 | Metabolism of cofactors | Cofactor and vitamin metabolism | M00577 | Biotin biosynthesis, BioW pathway, pimelate => pimeloyl | bioD | dethiobiotin synthetase [EC:6.3.3.3] | FL-ls |
| K00796 | Metabolism of cofactors | Cofactor and vitamin metabolism | M00841 | Tetrahydrofolate biosynthesis, mediated by PTPS, GTP | folP | dihydropteroate synthase [EC:2.5.1.15] | FL-ls |
| K01495 | Metabolism of cofactors | Cofactor and vitamin metabolism | M00841 | Tetrahydrofolate biosynthesis, mediated by PTPS, GTP | GCH1, folE | GTP cyclohydrolase IA [EC:3.5.4.16] | FL-ls |
| K01737 | Metabolism of cofactors | Cofactor and vitamin metabolism | M00842 | Tetrahydrobiopterin biosynthesis, GTP => BH4 [PATH:m | queD, ptpS, PTS | 6-pyruvoyltetrahydropterin/6-carboxytetrahydropterin sy | FL-ls |
| K01495 | Metabolism of cofactors | Cofactor and vitamin metabolism | M00842 | Tetrahydrobiopterin biosynthesis, GTP => BH4 [PATH:m | GCH1, folE | GTP cyclohydrolase IA [EC:3.5.4.16] | FL-ls |
| K01737 | Metabolism of cofactors | Cofactor and vitamin metabolism | M00843 | L-threo-Tetrahydrobiopterin biosynthesis, GTP => L-thre | queD, ptpS, PTS | 6-pyruvoyltetrahydropterin/6-carboxytetrahydropterin sy | FL-ls |
| K01495 | Metabolism of cofactors | Cofactor and vitamin metabolism | M00843 | L-threo-Tetrahydrobiopterin biosynthesis, GTP => L-thre | GCH1, folE | GTP cyclohydrolase IA [EC:3.5.4.16] | FL-ls |
| K01845 | Metabolism of cofactors | Cofactor and vitamin metabolism | M00846 | Siroheme biosynthesis, glutamate => siroheme [PATH:m | hemL | glutamate-1-semialdehyde 2,1-aminomutase [EC:5.4.3.8 | FL-ls |
| K02492 | Metabolism of cofactors | Cofactor and vitamin metabolism | M00846 | Siroheme biosynthesis, glutamate => siroheme [PATH:m | hemA | glutamyl-tRNA reductase [EC:1.2.1.70] | FL-ls |
| K01698 | Metabolism of cofactors | Cofactor and vitamin metabolism | M00846 | Siroheme biosynthesis, glutamate => siroheme [PATH:m | hemB, ALAD | porphobilinogen synthase [EC:4.2.1.24] | FL-ls |
| K02304 | Metabolism of cofactors | Cofactor and vitamin metabolism | M00846 | Siroheme biosynthesis, glutamate => siroheme [PATH:m | MET8 | precorrin-2 dehydrogenase / sirohydrochlorin ferrochelat | FL-ls |
| K02303 | Metabolism of cofactors | Cofactor and vitamin metabolism | M00846 | Siroheme biosynthesis, glutamate => siroheme [PATH:m | cobA | uroporphyrin-III C-methyltransferase [EC:2.1.1.107] | FL-ls |
| K00969 | Metabolism of cofactors | Cofactor and vitamin metabolism | M00115 | NAD biosynthesis, aspartate => NAD [PATH:map00760 | nadD | nicotinate-nucleotide adenylyltransferase [EC:2.7.7.18] | PA-ls |
| K03185 | Metabolism of cofactors | Cofactor and vitamin metabolism | M00117 | Ubiquinone biosynthesis, prokaryotes, chorismate => ub | ubiH | 2-octaprenyl-6-methoxyphenol hydroxylase [EC:1.14.13 | PA-ls |
| K00568 | Metabolism of cofactors | Cofactor and vitamin metabolism | M00117 | Ubiquinone biosynthesis, prokaryotes, chorismate => ub | ubiG | 2-polyprenyl-6-hydroxyphenyl methylase / 3-demethylub | PA-ls |
| K00859 | Metabolism of cofactors | Cofactor and vitamin metabolism | M00120 | Coenzyme A biosynthesis, pantothenate => CoA [PATH | coaE | dephospho-CoA kinase [EC:2.7.1.24] | PA-ls |
| K00867 | Metabolism of cofactors | Cofactor and vitamin metabolism | M00120 | Coenzyme A biosynthesis, pantothenate => CoA [PATH | coaA | type I pantothenate kinase [EC:2.7.1.33] | PA-ls |
| K03525 | Metabolism of cofactors | Cofactor and vitamin metabolism | M00120 | Coenzyme A biosynthesis, pantothenate => CoA [PATH | coaX | type III pantothenate kinase [EC:2.7.1.33] | PA-ls |
| K00228 | Metabolism of cofactors | Cofactor and vitamin metabolism | M00121 | Heme biosynthesis, glutamate => heme [PATH:map0086 | CPOX, hemF | coproporphyrinogen III oxidase [EC:1.3.3.3] | PA-ls |
| K02495 | Metabolism of cofactors | Cofactor and vitamin metabolism | M00121 | Heme biosynthesis, glutamate => heme [PATH:map0086 | hemN, hemZ | oxygen-independent coproporphyrinogen III oxidase [EC | PA-ls |
| K01772 | Metabolism of cofactors | Cofactor and vitamin metabolism | M00121 | Heme biosynthesis, glutamate => heme [PATH:map0086 | hemH, FECH | protoporphyrin/coproporphyrin ferrochelatase [EC:4.99.1 | PA-ls |
| K00231 | Metabolism of cofactors | Cofactor and vitamin metabolism | M00121 | Heme biosynthesis, glutamate => heme [PATH:map0086 | PPOX, hemY | protoporphyrinogen/coproporphyrinogen III oxidase [EC: | PA-ls |
| K01599 | Metabolism of cofactors | Cofactor and vitamin metabolism | M00121 | Heme biosynthesis, glutamate => heme [PATH:map0086 | hemE, UROD | uroporphyrinogen decarboxylase [EC:4.1.1.37] | PA-ls |
| K01719 | Metabolism of cofactors | Cofactor and vitamin metabolism | M00121 | Heme biosynthesis, glutamate => heme [PATH:map0086 | hemD, UROS | uroporphyrinogen-III synthase [EC:4.2.1.75] | PA-ls |
| K00652 | Metabolism of cofactors | Cofactor and vitamin metabolism | M00123 | Biotin biosynthesis, pimeloyl-ACP/CoA => biotin [PATH:r | bioF | 8-amino-7-oxononanoate synthase [EC:2.3.1.47] | PA-ls |
| K00275 | Metabolism of cofactors | Cofactor and vitamin metabolism | M00124 | Pyridoxal biosynthesis, erythrose-4P => pyridoxal-5P [P | pdxH, PNPO | pyridoxamine 5'-phosphate oxidase [EC:1.4.3.5] | PA-ls |
| K14652 | Metabolism of cofactors | Cofactor and vitamin metabolism | M00125 | Riboflavin biosynthesis, GTP => riboflavin/FMN/FAD [PA | ribBA | 3,4-dihydroxy 2-butanone 4-phosphate synthase / GTP | PA-ls |
| K11753 | Metabolism of cofactors | Cofactor and vitamin metabolism | M00125 | Riboflavin biosynthesis, GTP => riboflavin/FMN/FAD [PA | ribF | riboflavin kinase / FMN adenylyltransferase [EC:2.7.1.2 | PA-ls |
| K00950 | Metabolism of cofactors | Cofactor and vitamin metabolism | M00126 | Tetrahydrofolate biosynthesis, GTP => THF [PATH:map0 | folK | 2-amino-4-hydroxy-6-hydroxymethyldihydropteridine dipl | PA-ls |
| K01633 | Metabolism of cofactors | Cofactor and vitamin metabolism | M00126 | Tetrahydrofolate biosynthesis, GTP => THF [PATH:map0 | folB | 7,8-dihydroneopterin aldolase/epimerase/oxygenase [E | PA-ls |
| K00878 | Metabolism of cofactors | Cofactor and vitamin metabolism | M00127 | Thiamine biosynthesis, AIR => thiamine-P/thiamine-2P [ | thiM | hydroxyethylthiazole kinase [EC:2.7.1.50] | PA-ls |
| K00788 | Metabolism of cofactors | Cofactor and vitamin metabolism | M00127 | Thiamine biosynthesis, AIR => thiamine-P/thiamine-2P [ | thiE | thiamine-phosphate pyrophosphorylase [EC:2.5.1.3] | PA-ls |
| K00652 | Metabolism of cofactors | Cofactor and vitamin metabolism | M00573 | Biotin biosynthesis, BioI pathway, long-chain-acyl-ACP | bioF | 8-amino-7-oxononanoate synthase [EC:2.3.1.47] | PA-ls |
| K00652 | Metabolism of cofactors | Cofactor and vitamin metabolism | M00577 | Biotin biosynthesis, BioW pathway, pimelate => pimeloyl | bioF | 8-amino-7-oxononanoate synthase [EC:2.3.1.47] | PA-ls |
| K14652 | Metabolism of cofactors | Cofactor and vitamin metabolism | M00840 | Tetrahydrofolate biosynthesis, mediated by ribA and trp | ribBA | 3,4-dihydroxy 2-butanone 4-phosphate synthase / GTP | PA-ls |
| K01633 | Metabolism of cofactors | Cofactor and vitamin metabolism | M00840 | Tetrahydrofolate biosynthesis, mediated by ribA and trp | folB | 7,8-dihydroneopterin aldolase/epimerase/oxygenase [E | PA-ls |
| K00950 | Metabolism of cofactors | Cofactor and vitamin metabolism | M00841 | Tetrahydrofolate biosynthesis, mediated by PTPS, GTP | folK | 2-amino-4-hydroxy-6-hydroxymethyldihydropteridine dipl | PA-ls |
| K02302 | Metabolism of cofactors | Cofactor and vitamin metabolism | M00846 | Siroheme biosynthesis, glutamate => siroheme [PATH:m | cysG | uroporphyrin-III C-methyltransferase / precorrin-2 dehyd | PA-ls |
| K01719 | Metabolism of cofactors | Cofactor and vitamin metabolism | M00846 | Siroheme biosynthesis, glutamate => siroheme [PATH:m | hemD, UROS | uroporphyrinogen-III synthase [EC:4.2.1.75] | PA-ls |
| K05928 | Metabolism of cofactors | Cofactor and vitamin metabolism | M00112 | Tocopherol/tocotorienol biosynthesis [PATH:map00130 | E2.1.1.95 | tocopherol O-methyltransferase [EC:2.1.1.95] | shared |
| K01950 | Metabolism of cofactors | Cofactor and vitamin metabolism | M00115 | NAD biosynthesis, aspartate => NAD [PATH:map00760 | E6.3.5.1, NADSYN1 | NAD+ synthase (glutamine-hydrolysing) [EC:6.3.5.1] | shared |
| K01916 | Metabolism of cofactors | Cofactor and vitamin metabolism | M00115 | NAD biosynthesis, aspartate => NAD [PATH:map00760 | nadE | NAD+ synthase [EC:6.3.1.5] | shared |
| K03517 | Metabolism of cofactors | Cofactor and vitamin metabolism | M00115 | NAD biosynthesis, aspartate => NAD [PATH:map00760 | nadA | quinolinate synthase [EC:2.5.1.72] | shared |
| K02551 | Metabolism of cofactors | Cofactor and vitamin metabolism | M00116 | Menaquinone biosynthesis, chorismate => menaquinol [ | menD | 2-succinyl-5-enolpyruvyl-6-hydroxy-3-cyclohexene-1-car | shared |

| | | | | | | | |
|---|---|---|---|---|---|---|---|
| K08680 | Metabolism of cofactors | Cofactor and vitamin metabolism | M00116 | Menaquinone biosynthesis, chorismate => menaquinol [ | menH | 2-succinyl-6-hydroxy-2,4-cyclohexadiene-1-carboxylate | shared |
| K01661 | Metabolism of cofactors | Cofactor and vitamin metabolism | M00116 | Menaquinone biosynthesis, chorismate => menaquinol [ | menB | naphthoate synthase [EC:4.1.3.36] | shared |
| K02549 | Metabolism of cofactors | Cofactor and vitamin metabolism | M00116 | Menaquinone biosynthesis, chorismate => menaquinol [ | menC | O-succinylbenzoate synthase [EC:4.2.1.113] | shared |
| K01911 | Metabolism of cofactors | Cofactor and vitamin metabolism | M00116 | Menaquinone biosynthesis, chorismate => menaquinol [ | menE | O-succinylbenzoic acid---CoA ligase [EC:6.2.1.26] | shared |
| K03184 | Metabolism of cofactors | Cofactor and vitamin metabolism | M00117 | Ubiquinone biosynthesis, prokaryotes, chorismate => ub | ubiF | 3-demethoxyubiquinol 3-hydroxylase [EC:1.14.99.60] | shared |
| K03179 | Metabolism of cofactors | Cofactor and vitamin metabolism | M00117 | Ubiquinone biosynthesis, prokaryotes, chorismate => ub | ubiA | 4-hydroxybenzoate polyprenyltransferase [EC:2.5.1.39] | shared |
| K03181 | Metabolism of cofactors | Cofactor and vitamin metabolism | M00117 | Ubiquinone biosynthesis, prokaryotes, chorismate => ub | ubiC | chorismate--pyruvate lyase [EC:4.1.3.40] | shared |
| K00954 | Metabolism of cofactors | Cofactor and vitamin metabolism | M00120 | Coenzyme A biosynthesis, pantothenate => CoA [PATH | E2.7.7.3A, coaD, k | pantetheine-phosphate adenylyltransferase [EC:2.7.7.3] | shared |
| K02201 | Metabolism of cofactors | Cofactor and vitamin metabolism | M00120 | Coenzyme A biosynthesis, pantothenate => CoA [PATH | E2.7.7.3B | pantetheine-phosphate adenylyltransferase [EC:2.7.7.3] | shared |
| K01598 | Metabolism of cofactors | Cofactor and vitamin metabolism | M00120 | Coenzyme A biosynthesis, pantothenate => CoA [PATH | PPCDC, coaC | phosphopantothenoylcysteine decarboxylase [EC:4.1.1. | shared |
| K01749 | Metabolism of cofactors | Cofactor and vitamin metabolism | M00121 | Heme biosynthesis, glutamate => heme [PATH:map0086 | hemC, HMBS | hydroxymethylbilane synthase [EC:2.5.1.61] | shared |
| K00230 | Metabolism of cofactors | Cofactor and vitamin metabolism | M00121 | Heme biosynthesis, glutamate => heme [PATH:map0086 | hemG | menaquinone-dependent protoporphyrinogen oxidase [E | shared |
| K13542 | Metabolism of cofactors | Cofactor and vitamin metabolism | M00121 | Heme biosynthesis, glutamate => heme [PATH:map0086 | cobA-hemD | uroporphyrinogen III methyltransferase / synthase [EC:2 | shared |
| K13543 | Metabolism of cofactors | Cofactor and vitamin metabolism | M00121 | Heme biosynthesis, glutamate => heme [PATH:map0086 | hemDX | uroporphyrinogen III methyltransferase / synthase [EC:2 | shared |
| K02231 | Metabolism of cofactors | Cofactor and vitamin metabolism | M00122 | Cobalamin biosynthesis, cobinamide => cobalamin [PAT | cobP, cobU | adenosylcobinamide kinase / adenosylcobinamide-phos | shared |
| K02226 | Metabolism of cofactors | Cofactor and vitamin metabolism | M00122 | Cobalamin biosynthesis, cobinamide => cobalamin [PAT | cobC, phpB | alpha-ribazole phosphatase [EC:3.1.3.73] | shared |
| K02225 | Metabolism of cofactors | Cofactor and vitamin metabolism | M00122 | Cobalamin biosynthesis, cobinamide => cobalamin [PAT | cobC1, cobC | cobalamin biosynthetic protein CobC | shared |
| K00768 | Metabolism of cofactors | Cofactor and vitamin metabolism | M00122 | Cobalamin biosynthesis, cobinamide => cobalamin [PAT | E2.4.2.21, cobU, c | nicotinate-nucleotide--dimethylbenzimidazole phosphorib | shared |
| K00097 | Metabolism of cofactors | Cofactor and vitamin metabolism | M00124 | Pyridoxal biosynthesis, erythrose-4P => pyridoxal-5P [P | pdxA | 4-hydroxythreonine-4-phosphate dehydrogenase [EC:1.1 | shared |
| K03472 | Metabolism of cofactors | Cofactor and vitamin metabolism | M00124 | Pyridoxal biosynthesis, erythrose-4P => pyridoxal-5P [P | epd | D-erythrose 4-phosphate dehydrogenase [EC:1.2.1.72] | shared |
| K03473 | Metabolism of cofactors | Cofactor and vitamin metabolism | M00124 | Pyridoxal biosynthesis, erythrose-4P => pyridoxal-5P [P | pdxB | erythronate-4-phosphate dehydrogenase [EC:1.1.1.290] | shared |
| K03474 | Metabolism of cofactors | Cofactor and vitamin metabolism | M00124 | Pyridoxal biosynthesis, erythrose-4P => pyridoxal-5P [P | pdxJ | pyridoxine 5-phosphate synthase [EC:2.6.99.2] | shared |
| K00082 | Metabolism of cofactors | Cofactor and vitamin metabolism | M00125 | Riboflavin biosynthesis, GTP => riboflavin/FMN/FAD [PA | ribD2 | 5-amino-6-(5-phosphoribosylamino)uracil reductase [EC: | shared |
| K00953 | Metabolism of cofactors | Cofactor and vitamin metabolism | M00125 | Riboflavin biosynthesis, GTP => riboflavin/FMN/FAD [PA | FLAD1 | FAD synthetase [EC:2.7.7.2] | shared |
| K13941 | Metabolism of cofactors | Cofactor and vitamin metabolism | M00126 | Tetrahydrofolate biosynthesis, GTP => THF [PATH:map0 | folKP | 2-amino-4-hydroxy-6-hydroxymethyldihydropteridine diph | shared |
| K11754 | Metabolism of cofactors | Cofactor and vitamin metabolism | M00126 | Tetrahydrofolate biosynthesis, GTP => THF [PATH:map0 | folC | dihydrofolate synthase / folylpolyglutamate synthase [E | shared |
| K13940 | Metabolism of cofactors | Cofactor and vitamin metabolism | M00126 | Tetrahydrofolate biosynthesis, GTP => THF [PATH:map0 | sulD | dihydroneopterin aldolase / 2-amino-4-hydroxy-6-hydrox | shared |
| K08310 | Metabolism of cofactors | Cofactor and vitamin metabolism | M00126 | Tetrahydrofolate biosynthesis, GTP => THF [PATH:map0 | nudB, ntpA | dihydroneopterin triphosphate diphosphatase [EC:3.6.1. | shared |
| K09007 | Metabolism of cofactors | Cofactor and vitamin metabolism | M00126 | Tetrahydrofolate biosynthesis, GTP => THF [PATH:map0 | folE2 | GTP cyclohydrolase IB [EC:3.5.4.16] | shared |
| K14153 | Metabolism of cofactors | Cofactor and vitamin metabolism | M00127 | Thiamine biosynthesis, AIR => thiamine-P/thiamine-2P [ | thiDE | hydroxymethylpyrimidine kinase / phosphomethylpyrimid | shared |
| K00941 | Metabolism of cofactors | Cofactor and vitamin metabolism | M00127 | Thiamine biosynthesis, AIR => thiamine-P/thiamine-2P [ | thiD | hydroxymethylpyrimidine/phosphomethylpyrimidine kinas | shared |
| K14154 | Metabolism of cofactors | Cofactor and vitamin metabolism | M00127 | Thiamine biosynthesis, AIR => thiamine-P/thiamine-2P [ | THI6 | thiamine-phosphate diphosphorylase / hydroxyethylthiaz | shared |
| K06127 | Metabolism of cofactors | Cofactor and vitamin metabolism | M00128 | Ubiquinone biosynthesis, eukaryotes, 4-hydroxybenzoat | COQ5 | 2-methoxy-6-polyprenyl-1,4-benzoquinol methylase [EC: | shared |
| K06134 | Metabolism of cofactors | Cofactor and vitamin metabolism | M00128 | Ubiquinone biosynthesis, eukaryotes, 4-hydroxybenzoat | COQ7 | 3-demethoxyubiquinol 3-hydroxylase [EC:1.14.99.60] | shared |
| K06125 | Metabolism of cofactors | Cofactor and vitamin metabolism | M00128 | Ubiquinone biosynthesis, eukaryotes, 4-hydroxybenzoat | COQ2 | 4-hydroxybenzoate polyprenyltransferase [EC:2.5.1.39] | shared |
| K06126 | Metabolism of cofactors | Cofactor and vitamin metabolism | M00128 | Ubiquinone biosynthesis, eukaryotes, 4-hydroxybenzoat | COQ6 | ubiquinone biosynthesis monooxygenase Coq6 [EC:1.14 | shared |
| K00288 | Metabolism of cofactors | Cofactor and vitamin metabolism | M00141 | C1-unit interconversion, eukaryotes [PATH:map00670 m | MTHFD | methylenetetrahydrofolate dehydrogenase (NADP+) / me | shared |
| K13403 | Metabolism of cofactors | Cofactor and vitamin metabolism | M00141 | C1-unit interconversion, eukaryotes [PATH:map00670 m | MTHFD2 | methylenetetrahydrofolate dehydrogenase(NAD+) / 5,10- | shared |
| K02169 | Metabolism of cofactors | Cofactor and vitamin metabolism | M00572 | Pimeloyl-ACP biosynthesis, BioC-BioH pathway, malonyl | bioC | malonyl-CoA O-methyltransferase [EC:2.1.1.197] | shared |
| K02170 | Metabolism of cofactors | Cofactor and vitamin metabolism | M00572 | Pimeloyl-ACP biosynthesis, BioC-BioH pathway, malonyl | bioH | pimeloyl-[acyl-carrier protein] methyl ester esterase [EC | shared |
| K01906 | Metabolism of cofactors | Cofactor and vitamin metabolism | M00577 | Biotin biosynthesis, BioW pathway, pimelate => pimeloyl | bioW | 6-carboxyhexanoate--CoA ligase [EC:6.2.1.14] | shared |
| K14974 | Metabolism of cofactors | Cofactor and vitamin metabolism | M00622 | Nicotinate degradation, nicotinate => fumarate [PATH:m | nicC | 6-hydroxynicotinate 3-monooxygenase [EC:1.14.13.114] | shared |
| K13941 | Metabolism of cofactors | Cofactor and vitamin metabolism | M00840 | Tetrahydrofolate biosynthesis, mediated by ribA and trp | folKP | 2-amino-4-hydroxy-6-hydroxymethyldihydropteridine diph | shared |
| K11754 | Metabolism of cofactors | Cofactor and vitamin metabolism | M00841 | Tetrahydrofolate biosynthesis, mediated by PTPS, GTP | folC | dihydrofolate synthase / folylpolyglutamate synthase [E | shared |
| K01749 | Metabolism of cofactors | Cofactor and vitamin metabolism | M00846 | Siroheme biosynthesis, glutamate => siroheme [PATH:m | hemC, HMBS | hydroxymethylbilane synthase [EC:2.5.1.61] | shared |
| K02496 | Metabolism of cofactors | Cofactor and vitamin metabolism | M00846 | Siroheme biosynthesis, glutamate => siroheme [PATH:m | hemX | uroporphyrin-III C-methyltransferase [EC:2.1.1.107] | shared |
| K13542 | Metabolism of cofactors | Cofactor and vitamin metabolism | M00846 | Siroheme biosynthesis, glutamate => siroheme [PATH:m | cobA-hemD | uroporphyrinogen III methyltransferase / synthase [EC:2 | shared |
| K13543 | Metabolism of cofactors | Cofactor and vitamin metabolism | M00846 | Siroheme biosynthesis, glutamate => siroheme [PATH:m | hemDX | uroporphyrinogen III methyltransferase / synthase [EC:2 | shared |
| K01588 | Nucleotide metabolism | Purine metabolism | M00048 | Inosine monophosphate biosynthesis, PRPP + glutamine | purE | 5-(carboxyamino)imidazole ribonucleotide mutase [EC:5. | FL-ls |
| K01589 | Nucleotide metabolism | Purine metabolism | M00048 | Inosine monophosphate biosynthesis, PRPP + glutamine | purK | 5-(carboxyamino)imidazole ribonucleotide synthase [EC: | FL-ls |
| K06863 | Nucleotide metabolism | Purine metabolism | M00048 | Inosine monophosphate biosynthesis, PRPP + glutamine | purP | 5-formaminoimidazole-4-carboxamide-1-(beta)-D-ribofura | FL-ls |
| K13713 | Nucleotide metabolism | Purine metabolism | M00048 | Inosine monophosphate biosynthesis, PRPP + glutamine | purCD | fusion protein PurCD [EC:6.3.2.6 6.3.4.13] | FL-ls |
| K01945 | Nucleotide metabolism | Purine metabolism | M00048 | Inosine monophosphate biosynthesis, PRPP + glutamine | purD | phosphoribosylamine---glycine ligase [EC:6.3.4.13] | FL-ls |
| K01923 | Nucleotide metabolism | Purine metabolism | M00048 | Inosine monophosphate biosynthesis, PRPP + glutamine | purC | phosphoribosylaminoimidazole-succinocarboxamide syn | FL-ls |
| K01933 | Nucleotide metabolism | Purine metabolism | M00048 | Inosine monophosphate biosynthesis, PRPP + glutamine | purM | phosphoribosylformylglycinamidine cyclo-ligase [EC:6.3. | FL-ls |
| K01952 | Nucleotide metabolism | Purine metabolism | M00048 | Inosine monophosphate biosynthesis, PRPP + glutamine | purL, PFAS | phosphoribosylformylglycinamidine synthase [EC:6.3.5. | FL-ls |
| K00939 | Nucleotide metabolism | Purine metabolism | M00049 | Adenine ribonucleotide biosynthesis, IMP => ADP,ATP [ | adk, AK | adenylate kinase [EC:2.7.4.3] | FL-ls |
| K01466 | Nucleotide metabolism | Purine metabolism | M00546 | Purine degradation, xanthine => urea [PATH:map00230 | allB | allantoinase [EC:3.5.2.5] | FL-ls |
| K00087 | Nucleotide metabolism | Purine metabolism | M00546 | Purine degradation, xanthine => urea [PATH:map00230 | ygeS, xdhA | xanthine dehydrogenase molybdenum-binding subunit [E | FL-ls |
| K00942 | Nucleotide metabolism | Purine metabolism | M00050 | Guanine ribonucleotide biosynthesis IMP => GDP,GTP [ | E2.7.4.8, gmk | guanylate kinase [EC:2.7.4.8] | PA-ls |

| | | | | | | | |
|---|---|---|---|---|---|---|---|
| K07127 | Nucleotide metabolism | Purine metabolism | M00546 | Purine degradation, xanthine => urea [PATH:map00230 | uraH, pucM, hiuH | 5-hydroxyisourate hydrolase [EC:3.5.2.17] | PA-ls |
| K01477 | Nucleotide metabolism | Purine metabolism | M00546 | Purine degradation, xanthine => urea [PATH:map00230 | alc, ALLC | allantoicase [EC:3.5.3.4] | PA-ls |
| K13482 | Nucleotide metabolism | Purine metabolism | M00546 | Purine degradation, xanthine => urea [PATH:map00230 | xdhB | xanthine dehydrogenase large subunit [EC:1.17.1.4] | PA-ls |
| K13481 | Nucleotide metabolism | Purine metabolism | M00546 | Purine degradation, xanthine => urea [PATH:map00230 | xdhA | xanthine dehydrogenase small subunit [EC:1.17.1.4] | PA-ls |
| K11177 | Nucleotide metabolism | Purine metabolism | M00546 | Purine degradation, xanthine => urea [PATH:map00230 | yagR | xanthine dehydrogenase YagR molybdenum-binding sub | PA-ls |
| K11178 | Nucleotide metabolism | Purine metabolism | M00546 | Purine degradation, xanthine => urea [PATH:map00230 | yagS | xanthine dehydrogenase YagS FAD-binding subunit [EC | PA-ls |
| K11788 | Nucleotide metabolism | Purine metabolism | M00048 | Inosine monophosphate biosynthesis, PRPP + glutamine | ADE5 | phosphoribosylamine--glycine ligase / phosphoribosylfor | shared |
| K11808 | Nucleotide metabolism | Purine metabolism | M00048 | Inosine monophosphate biosynthesis, PRPP + glutamine | ADE2 | phosphoribosylaminoimidazole carboxylase [EC:4.1.1.21 | shared |
| K01587 | Nucleotide metabolism | Purine metabolism | M00048 | Inosine monophosphate biosynthesis, PRPP + glutamine | PAICS | phosphoribosylaminoimidazole carboxylase / phosphorib | shared |
| K13485 | Nucleotide metabolism | Purine metabolism | M00546 | Purine degradation, xanthine => urea [PATH:map00230 | PRHOXNB, URAD | 2-oxo-4-hydroxy-4-carboxy-5-ureidoimidazoline decarbox | shared |
| K16840 | Nucleotide metabolism | Purine metabolism | M00546 | Purine degradation, xanthine => urea [PATH:map00230 | hpxQ | 2-oxo-4-hydroxy-4-carboxy-5-ureidoimidazoline decarbox | shared |
| K13484 | Nucleotide metabolism | Purine metabolism | M00546 | Purine degradation, xanthine => urea [PATH:map00230 | TTHL | 5-hydroxyisourate hydrolase / 2-oxo-4-hydroxy-4-carbox | shared |
| K16842 | Nucleotide metabolism | Purine metabolism | M00546 | Purine degradation, xanthine => urea [PATH:map00230 | hpxB | allantoinase [EC:3.5.2.5] | shared |
| K13479 | Nucleotide metabolism | Purine metabolism | M00546 | Purine degradation, xanthine => urea [PATH:map00230 | ygeT, xdhB | xanthine dehydrogenase FAD-binding subunit [EC:1.17.1 | shared |
| K13483 | Nucleotide metabolism | Purine metabolism | M00546 | Purine degradation, xanthine => urea [PATH:map00230 | yagT | xanthine dehydrogenase YagT iron-sulfur-binding subuni | shared |
| K06016 | Nucleotide metabolism | Pyrimidine metabolism | M00046 | Pyrimidine degradation, uracil => beta-alanine, thymine = | pydC | beta-ureidopropionase / N-carbamoyl-L-amino-acid hydro | FL-ls |
| K00254 | Nucleotide metabolism | Pyrimidine metabolism | M00051 | Uridine monophosphate biosynthesis, glutamine (+ PRPP | DHODH, pyrD | dihydroorotate dehydrogenase [EC:1.3.5.2] | FL-ls |
| K00762 | Nucleotide metabolism | Pyrimidine metabolism | M00051 | Uridine monophosphate biosynthesis, glutamine (+ PRPP | pyrE | orotate phosphoribosyltransferase [EC:2.4.2.10] | FL-ls |
| K01591 | Nucleotide metabolism | Pyrimidine metabolism | M00051 | Uridine monophosphate biosynthesis, glutamine (+ PRPP | pyrF | orotidine-5'-phosphate decarboxylase [EC:4.1.1.23] | FL-ls |
| K00945 | Nucleotide metabolism | Pyrimidine metabolism | M00052 | Pyrimidine ribonucleotide biosynthesis, UMP => UDP/UT | cmk | CMP/dCMP kinase [EC:2.7.4.25] | FL-ls |
| K01937 | Nucleotide metabolism | Pyrimidine metabolism | M00052 | Pyrimidine ribonucleotide biosynthesis, UMP => UDP/UT | pyrG, CTPS | CTP synthase [EC:6.3.4.2] | FL-ls |
| K00943 | Nucleotide metabolism | Pyrimidine metabolism | M00053 | Pyrimidine deoxyribonuleotide biosynthesis, CDP/CTP = | tmk, DTYMK | dTMP kinase [EC:2.7.4.9] | FL-ls |
| K01465 | Nucleotide metabolism | Pyrimidine metabolism | M00051 | Uridine monophosphate biosynthesis, glutamine (+ PRPP | URA4, pyrC | dihydroorotase [EC:3.5.2.3] | PA-ls |
| K01494 | Nucleotide metabolism | Pyrimidine metabolism | M00053 | Pyrimidine deoxyribonuleotide biosynthesis, CDP/CTP = | dcd | dCTP deaminase [EC:3.5.4.13] | PA-ls |
| K01520 | Nucleotide metabolism | Pyrimidine metabolism | M00053 | Pyrimidine deoxyribonuleotide biosynthesis, CDP/CTP = | dut, DUT | dUTP pyrophosphatase [EC:3.6.1.23] | PA-ls |
| K00226 | Nucleotide metabolism | Pyrimidine metabolism | M00051 | Uridine monophosphate biosynthesis, glutamine (+ PRPP | pyrD | dihydroorotate dehydrogenase (fumarate) [EC:1.3.98.1] | shared |
| K07550 | Xenobiotics biodegradatio | Aromatics degradation | M00418 | Toluene degradation, anaerobic, toluene => benzoyl-CoA | bbsB | benzoylsuccinyl-CoA thiolase BbsB subunit [EC:2.3.1.-] | FL-ls |
| K07543 | Xenobiotics biodegradatio | Aromatics degradation | M00418 | Toluene degradation, anaerobic, toluene => benzoyl-CoA | bbsE | benzylsuccinate CoA-transferase BbsE subunit [EC:2.8. | FL-ls |
| K07544 | Xenobiotics biodegradatio | Aromatics degradation | M00418 | Toluene degradation, anaerobic, toluene => benzoyl-CoA | bbsF | benzylsuccinate CoA-transferase BbsF subunit [EC:2.8. | FL-ls |
| K10616 | Xenobiotics biodegradatio | Aromatics degradation | M00419 | Cymene degradation, p-cymene => p-cumate [PATH:map | cymAa | p-cymene methyl-monooxygenase [EC:1.14.15.25] | PA-ls |
| K14727 | Xenobiotics biodegradatio | Aromatics degradation | M00568 | Catechol ortho-cleavage, catechol => 3-oxoadipate [PAT | pcaL | 3-oxoadipate enol-lactonase / 4-carboxymuconolactone | PA-ls |
| K03464 | Xenobiotics biodegradatio | Aromatics degradation | M00568 | Catechol ortho-cleavage, catechol => 3-oxoadipate [PAT | catC | muconolactone D-isomerase [EC:5.3.3.4] | PA-ls |
| K07546 | Xenobiotics biodegradatio | Aromatics degradation | M00418 | Toluene degradation, anaerobic, toluene => benzoyl-CoA | bbsH | E-phenylitaconyl-CoA hydratase [EC:4.2.1.-] | shared |
| K10618 | Xenobiotics biodegradatio | Aromatics degradation | M00419 | Cymene degradation, p-cymene => p-cumate [PATH:map | cymC | p-cumic aldehyde dehydrogenase | shared |
| K14584 | Xenobiotics biodegradatio | Aromatics degradation | M00534 | Naphthalene degradation, naphthalene => salicylate [PA | nahD | 2-hydroxychromene-2-carboxylate isomerase [EC:5.99.1 | shared |
| K10620 | Xenobiotics biodegradatio | Aromatics degradation | M00539 | Cumate degradation, p-cumate => 2-oxopent-4-enoate + | cmtB | 2,3-dihydroxy-2,3-dihydro-p-cumate dehydrogenase [EC | shared |
| K10619 | Xenobiotics biodegradatio | Aromatics degradation | M00539 | Cumate degradation, p-cumate => 2-oxopent-4-enoate + | cmtAb | p-cumate 2,3-dioxygenase subunit alpha [EC:1.14.12.25 | shared |
| K07535 | Xenobiotics biodegradatio | Aromatics degradation | M00540 | Benzoate degradation, cyclohexanecarboxylic acid =>pi | badH | 2-hydroxycyclohexanecarboxyl-CoA dehydrogenase [EC | shared |
| K07536 | Xenobiotics biodegradatio | Aromatics degradation | M00540 | Benzoate degradation, cyclohexanecarboxylic acid =>pi | badI | 2-ketocyclohexanecarboxyl-CoA hydrolase [EC:3.1.2.-] | shared |
| K04117 | Xenobiotics biodegradatio | Aromatics degradation | M00540 | Benzoate degradation, cyclohexanecarboxylic acid =>pi | aliB | cyclohexanecarboxyl-CoA dehydrogenase [EC:1.3.99.-] | shared |
| K04116 | Xenobiotics biodegradatio | Aromatics degradation | M00540 | Benzoate degradation, cyclohexanecarboxylic acid =>pi | aliA | cyclohexanecarboxylate-CoA ligase [EC:6.2.1.-] | shared |
| K10222 | Xenobiotics biodegradatio | Aromatics degradation | M00543 | Biphenyl degradation, biphenyl => 2-oxopent-4-enoate + | bphD | 2,6-dioxo-6-phenylhexa-3-enoate hydrolase [EC:3.7.1.8] | shared |
| K08689 | Xenobiotics biodegradatio | Aromatics degradation | M00543 | Biphenyl degradation, biphenyl => 2-oxopent-4-enoate + | bphAa, bphA1, bphA | biphenyl 2,3-dioxygenase subunit alpha [EC:1.14.12.18] | shared |
| K08690 | Xenobiotics biodegradatio | Aromatics degradation | M00543 | Biphenyl degradation, biphenyl => 2-oxopent-4-enoate + | bphB | cis-2,3-dihydrobiphenyl-2,3-diol dehydrogenase [EC:1.3. | shared |
| K01055 | Xenobiotics biodegradatio | Aromatics degradation | M00568 | Catechol ortho-cleavage, catechol => 3-oxoadipate [PAT | pcaD | 3-oxoadipate enol-lactonase [EC:3.1.1.24] | shared |
| K04102 | Xenobiotics biodegradatio | Aromatics degradation | M00623 | Phthalate degradation, phthalate => protocatechuate [PA | pht5 | 4,5-dihydroxyphthalate decarboxylase [EC:4.1.1.55] | shared |
